# The Influence of Force Field Perturbations on Symmetric and Asymmetric Bimanual Reaching

**DOI:** 10.1101/2025.08.28.672982

**Authors:** Jaskanwaljeet Kaur, Ramesh Balasubramaniam

## Abstract

Bimanual symmetric and asymmetric reaching movements require the motor system to coordinate the two limbs under varying spatial and physical demands. While prior research has shown a strong tendency for temporal synchronization between the hands, little is known about how bimanual reaching is organized when each limb is subjected to distinct types of force fields, such as viscous or elastic loads. This study systematically investigated how matched (viscous/viscous, elastic/elastic) and mismatched (e.g., viscous/no load, elastic/no load, viscous/elastic) load conditions influence key timing and kinematic parameters during bimanual reaching tasks across different spatial configurations and force magnitudes. Results show that kinematic measures for each arm, including peak velocity, peak acceleration, and peak deceleration, exhibit significant differences across both matched and mismatched load conditions, reflecting clear adaptations in limb behavior. In contrast, temporal aspects of movement—specifically the timepoints of peak velocity, peak acceleration, peak deceleration, and movement end time—remain coupled under matched loads but become decoupled under mismatched loads, with each limb operating more independently regardless of the spatial configuration of the task. Together, these findings clarify how the motor system balances the need for coordinated bimanual performance with the flexibility required to adapt to varying load conditions. This study extends prior work by demonstrating that not all bimanual reaches are temporally synchronized; instead, task-specific factors such as load type and symmetry critically shape how the limbs coordinate, revealing the motor system’s capacity for flexible, context-dependent control.

## Introduction

Efficiently coordinating both hands is integral to performing daily tasks, such as pouring milk into a cup while stabilizing the container, highlighting the interdependence and cooperation between the limbs. Previous studies by Kelso et al., 1979; 1983 have shown strong coupling between the limbs during bimanual movements, with near synchronous onsets, peak velocities, and movement terminations, despite variations in hand speeds. The spatiotemporal characteristics of bimanual movements reveal robust interactions between the left and right limbs (Swinnen, 2002). Numerous studies on bimanual tasks have shown temporal synchrony in the movements of both hands. For instance, Keele (1986) observed that participants tended to synchronize their hands during bimanual aiming tasks, even without explicit instructions to do so. This synchronization extended to movement initiation and duration. Similarly, Jeannerod (1984) found synchronization not only in movement onset and duration but also in the timing of maximum hand velocity and grip aperture during bimanual prehension tasks. The inclination to synchronize both hands has been noted across tasks of varying difficulty levels (Kelso et al., 1979; Kelso et al., 1983).

While several studies have established a fundamental tendency for temporal synchronization between the two hands (Kelso et al., 1979; Kelso et al., 1983; Jeannerod, 1984; Jackson et al., 1999; Diedrichsen et al., 2001), bimanual aiming and prehension tasks with precision demands have revealed instances of asynchronous coordination. That is, the level of precision required can influence the temporal coordination between the hands. This observation suggests that the temporal synchrony or asynchrony between the limb movements is strongly influenced by contextual factors (Balakrishnan & Hinckley, 2000). A study investigating bimanual prehension tasks, where both hands reached to grasp two objects, showed discrepancy in timing attributed to differences in target distances, aligning with expectations from typical unimanual movements towards varying distances (Bingham et al., 2008). The degree of asynchrony increased as task difficulty increased. Furthermore, even when tasks complexities and movement distances were identical, initial synchronization during the acceleration phases transitions into disparate arrival times at the targets. This terminal-phase asynchrony has been attributed to the need for each hand to adjust independently to task-specific demands or perturbations. Studies applying sudden disturbances or changes in the movement environment during prehension tasks have shown that subtle adjustments in movement trajectories— such as changes in target location or load conditions—can impact interlimb coordination. More substantial changes in movement demands or direction can lead to sustained interference between the hands, suggesting that the motor system’s ability to maintain coordinated performance depends on the nature and magnitude of the challenge (Mason & Grabowski, 2010).

Taken together, these findings point to a nuanced understanding of how the two hands work together during bimanual tasks: while the motor system exhibits a strong tendency toward temporal synchrony, this coupling is not rigid or uniform across all situations. As Riek et al. (2003) discovered, synchrony at the start or end points of movement does not guarantee synchronous control throughout the entire reach; one hand may momentarily pause or adjust to allow the other to catch up, resulting in coordinated endpoints despite underlying differences in movement profiles. These studies suggest that the control of bimanual movements is highly task-dependent, shaped by the specific constraints, demands, and adjustments required within a given context. Importantly, the system flexibly shifts between tightly linked and more independent control strategies depending on what the task requires.

While prior research has consistently manipulated task complexity through changes in object size (Bruyn & Mason, 2009), movement distance (Carnahan, 1998; Bozzacchi et al., 2017), and subtle perturbations (Mason & Grabowski, 2010; Brunfeldt et al., 2021), the specific influence of applied force fields—such as viscous and elastic loads—on bimanual reaching movements has not been explored. Therefore, the primary objective of the current study is to investigate how the motor system coordinates the two hands during bimanual reaching when subjected to different load conditions, particularly focusing on targets at varying distances. We aim to explore how applying viscous and elastic loads affects movement performance, as these types of force fields may influence how the motor system coordinates the limbs during bimanual reaching. Specifically, we examine how a combination of matched or mismatched load conditions influences key movement parameters—include movement end time, peak velocity, timepoint at peak velocity, peak acceleration, timepoint at peak acceleration, peak deceleration and timepoint at peak deceleration—during bimanual reaching tasks across varying spatial configurations. Additionally, we aim to examine reaction time, movement time, and total response time; however, we anticipate that these measures will remain relatively stable across load conditions, as the instructions emphasized performing reaches as quickly and accurately as possible, encouraging consistent movement strategies across load conditions. This allows us to isolate how applied loads shape the execution phase of movement without necessarily altering the initiation or planning phases, providing a clearer picture of how the motor system flexibly adjusts bimanual performance under complex task conditions.

Building on prior research, we hypothesized that the introduction of applied force fields—specifically viscous and elastic loads—will significantly influence movement parameters during bimanual reaching tasks across varying spatial configurations. Specifically, we expected notable changes in peak velocity, acceleration, and deceleration when these loads are applied, while the timing of these measures (i.e., time to peak velocity, time to peak acceleration, time to peak deceleration) would remain relatively stable, reflecting the system’s effort to preserve overall temporal coordination despite the differing load conditions. We predicted that viscous loads, which apply velocity-dependent resistance, will reduce peak velocity and acceleration, as the motor system compensates for the damping effect that opposes rapid movement. In contrast, elastic loads, which introduce position-dependent resistance, were expected to alter the movement profile by increasing resistance near the end of the reach, potentially delaying when peak acceleration or deceleration occurs, without necessarily reducing their overall magnitude. Furthermore, we anticipated that mismatched load conditions—where one limb experiences viscous resistance and the other elastic resistance—would impose the greatest challenge for maintaining coordinated bimanual performance, requiring limb-specific adjustments and resulting in longer movement durations and reduced interlimb synchrony. Overall, these predictions aim to clarify how the motor system balances the need for coordinated performance against differing load conditions, offering insights into the flexible control strategies used during bimanual reaching under complex task demands.

## Methods

### Participants

Forty-three healthy participants took part in the study. Data from thirteen participants were excluded due to failure to follow task instructions. As a result, data from thirty participants were analyzed (age: 22.93 ± 4.13, 20 Female). The experiment was approved by the Institutional Review Board of the University of California, Merced, and conducted in accordance with the Declaration of Helsinki. All participants provided written informed consent prior to participation and were at least 18 years old, with normal or corrected-to-normal vision.

### Handedness Measurements

Handedness was evaluated using the 4-item Edinburgh Handedness Inventory (EHI)–Short Form (Veale, 2014). The EHI assesses hand dominance in daily activities (e.g., writing, throwing). The laterality quotient (LQ) of hand dominance ranges from −100 (left-handed) to 100 (right-handed), with an LQ between −100 and −61 indicating left-handedness, −60 to 60 indicating mixed-handedness, and 61 to 100 indicating right-handedness. In the present study, 100% of the participants were right-handed (*N* = 30).

### Apparatus

The experiment was performed using the Kinarm upper-limb robotic exoskeleton (BKIN Technologies Ltd., Kingston, ON, Canada). The Kinarm exoskeleton has been widely used in both non-human primate research and human studies, as well as in clinical assessments to diagnose stroke and other motor deficits (Bansil et al., 2012). Operating at sampling rates of up to 1000 Hz, the system enables accurate and precise quantification of motor and sensory characteristics in both healthy and neurologically impaired participants (Dukelow et al., 2010; Kenzie et al., 2014).

The Kinarm device includes a height-adjustable chair with bilateral arm and hand support platforms, connected to the operator’s computer. A monitor and an underlying screen display the task being performed (see Figure 1a for a task schematic), allowing participants to view and interact with visual stimuli within the same workspace where their arms move. This setup enables two-dimensional planar movements while supporting the arms to reduce fatigue, facilitating accurate measurement of bimanual coordination. Additionally, the Kinarm system can apply controlled perturbations or force fields to the arms, allowing researchers to investigate how such perturbations impact planned motor movements (Brown et al., 2007). During task performance, participants’ hand movements were continuously recorded using Dexterit-E software at a sampling rate of 1000 Hz (3.8v, BKIN Technologies Ltd., Ontario, Canada). All data were automatically saved in c3d file format, containing hand position coordinates (x, y) and derived measures of movement velocity and acceleration in the transverse plane.

**Figure 1.**
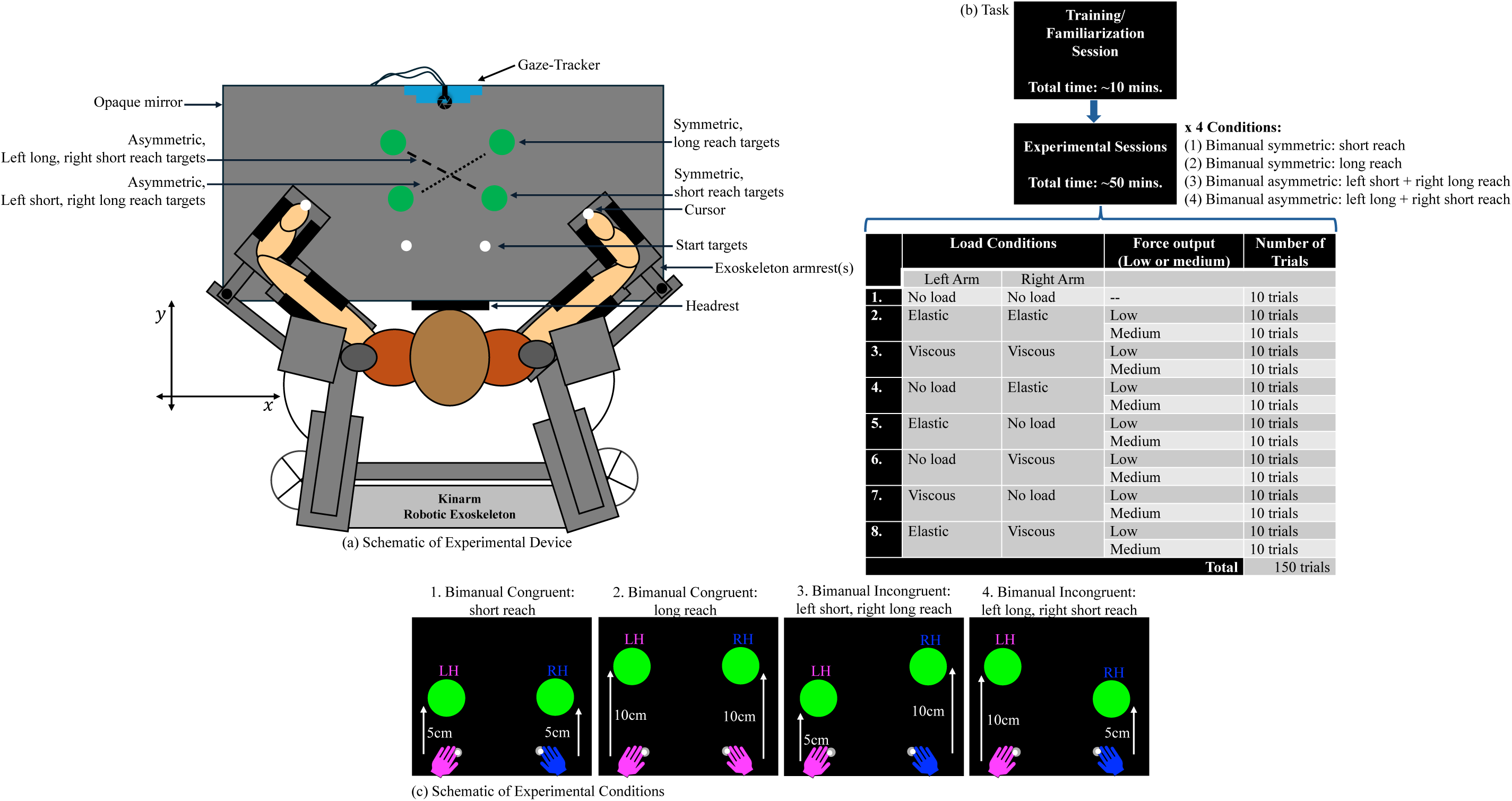
Experimental setup. **(a)** The Kinarm robotic exoskeleton with a seated participant. **(b)** Task design depicts the order of events in the bimanual reaching experiment. A training/familiarization session was completed so that participants could become accustomed to the bimanual movements performed during the task. The experiment consisted of a randomized set of trials across symmetric and asymmetric reaching conditions. Each condition included a No Load/No Load baseline (10 trials per condition) and force-loaded trials with either low force (elastic load = 30 N, viscous load = −15 N) or medium force (elastic load = 36.250 N, viscous load = −20 N). The reaching conditions were: (1) Symmetric long reach, (2) Symmetric short reach, (3) Asymmetric reach (left long, right short), and (4) Asymmetric reach (left short, right long). The force condition was randomized across trials. In total, participants completed 680 trials (40 baseline trials + 640 force trials). **(c)** Schematic showing the location of the 4 reaching movements participants performed during the experiment.

### Symmetric and Asymmetric Reaching Movements

We developed a bimanual reaching task using Simulink (R2015a, The MathWorks, USA) and Dexterit-E to investigate how perturbing the motor system with force fields—specifically viscous and/or elastic forces—affects bimanual reaching during symmetric and asymmetric conditions. The symmetric reaching movements included: a short reach, where participants moved both hands together from the start position toward a target 5 cm away; and a long reach, where participants moved both hands together toward a target 10 cm away. The asymmetric reaching movements involved: a left short reach, right long reach, where participants moved their left hand 5 cm and their right hand 10 cm toward the targets; and a left long reach, right short reach, where participants moved their left hand 10 cm and their right hand 5 cm toward the targets. During these movements, force fields were applied to the participants’ hands, with two possible types: viscous or elastic force field. The viscous force is velocity-dependent, acting like a dampener or friction on the limb, where an increase in velocity results in increased resistance. The elastic force, on the other hand, is position-dependent, with lateral force scaling relative to the hand’s position from the start. The force applied by the Kinarm robot was always orthogonal to the direction of movement, as described below:

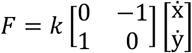

where *F* represents the force vector in the horizontal plane, and ẋ and ẏ denote the hand velocities in the horizontal plane. The variable *k* indicates the viscosity of the force field. In the low force condition, the viscous (velocity-dependent) force field has *k* = −15 N/m, and the elastic (position-dependent) force field has *k* = 30 N/m. In the medium force condition, the viscous (velocity-dependent) force field has *k* = −20 N/m, and the elastic (position-dependent) force field has *k* = 36.250 N/m.

As shown in Figure 1c, two target circles (representing hand positions) were displayed 5 cm apart. The target size and distance were determined based on pilot testing with adult participants (>18 years). For both symmetric and asymmetric trials, the experimental setup regarding the starting position remained constant, but the movement patterns varied depending on the type of reach being performed. Before beginning the task, participants underwent training on the reaching movements they would be performing during the experiment—these training trials were conducted without any force fields applied. Participants initially aligned their index fingers at a white target (the starting position). Once their hands were positioned at the start, randomized target locations were presented, with both the reaching movements and the force fields randomized across trials. The participants’ task was to reach both targets as quickly and accurately as possible.

A total of 8 load conditions were used in the experiment: 1 baseline (no load) condition and 7 each for low force and medium force outputs. These conditions were classified into 4 main reaching movements, as described earlier and shown in Figure 1b. The total number of trials was 680, consisting of 40 baseline trials (10 trials per condition) and 640 force trials (randomized across low and medium force conditions). Participants were not informed about the specifics of the force fields applied during the experiment; instead, they were encouraged to perform the reach as quickly and accurately as possible, or to the best of their ability, despite the perturbations. Pauses occurred between each trial, and after a 500 ms delay, the white targets reappeared for hand positioning. Participants aligned their hands to these starting targets before continuing with the experiment.

### Data Processing and Analysis

All raw data for each trial were saved in .c3d files per participant and imported into MATLAB for analysis. Hand kinematics were computed using proprietary MATLAB functions provided by BKIN Technologies and filtered using a 3rd-order, double-pass Butterworth filter with a 10 Hz, 3 dB cutoff—resulting in a 6th-order zero-lag filter. These processed data files were then used for offline data analysis. A custom MATLAB script was used to extract key dependent variables on a per trial basis.

As an initial step, measures of reaction time (RT), movement time (MT), and total response time (ResT) were computed to confirm consistent timing of responses across experimental conditions. Reaction time was calculated as the interval between the onset of the stimulus and the initiation of movement. Movement onset was defined as the moment when the limb’s velocity reached 5% of that trial’s peak velocity. Movement time for each limb was measured as the duration from movement onset to when the hand reached the target. Finally, total response time for each limb was calculated as the time from stimulus onset to the moment the hand reached the target. Full results for these measures are reported in the supplementary materials. Specifically, RT data visualizations are presented in supplementary figure 17, with linear mixed-effects (LME) results in supplementary table 1 and post hoc comparisons in supplementary tables 2 and 3. MT outcomes are shown in supplementary figure 18, with corresponding LME results in supplementary table 4 and post hoc comparisons in supplementary tables 5 and 6. Finally, ResT results are summarized in supplementary figure 19, with LME outcomes in supplementary table 7 and post hoc analyses in supplementary tables 8 and 9.

The primary variables analyzed in the main manuscript included Movement End Time (ME), defined as the timepoint at which the hand’s motion ceases; Peak Velocity (PV), corresponding to the maximum speed achieved during the reach; and Timepoint to Peak Velocity (TPV), which refers to the time at which peak velocity occurs. Additionally, Peak Acceleration (PA), defined as the maximum rate of change in velocity, and Timepoint to Peak Acceleration (TPA), the time at which peak acceleration occurs, were analyzed. Peak Deceleration (PD), representing the greatest rate of velocity decrease during the movement, and Timepoint to Peak Deceleration (TPD), the time at which maximum deceleration occurs, were also extracted and analyzed.

Kinematic data were processed across all trials using a custom MATLAB pipeline. Raw position signals from the saved Kinarm data were used to calculate hand velocity and acceleration. Velocity data were converted to centimeters per second (cm/s) and smoothed using a moving average filter to reduce high-frequency noise. Peak velocity was defined as the maximum value for the left hand and the minimum for the right hand, reflecting the direction-specific nature of the movement. The corresponding TPV was recorded in milliseconds. Acceleration was computed by taking the first derivative of the smoothed velocity signal and applying the same moving average smoothing. TPA was also recorded relative to movement onset. To estimate peak deceleration, the second most prominent acceleration peak—opposite in direction to the initial movement burst—was identified. For the right hand, this corresponded to the second-largest positive acceleration; for the left hand, the second-largest negative acceleration. This approach helped isolate true deceleration events, minimizing confounds with initial acceleration. The TPD was recorded accordingly.

Movement end time was defined as the moment when the participant’s hand first entered the target zone during the reaching task. This timepoint was marked by the Kinarm system as the participant reached the designated targets. The system continuously monitored the hand’s position relative to the target, and movement end time was recorded when the hand’s position fell within the predefined acceptance radius of the target for each trial. These time markers were used as the operational definition of movement end time in the subsequent analyses. All kinematic variables were extracted within a post-movement onset window ranging from 150 to 4500ms, selected to capture the full time course of the bimanual reaching movement—from initial acceleration to final movement termination— while excluding early signal noise and post-movement artifacts. Sub-windows were applied for specific variables as appropriate (e.g., 150-1000ms for peak acceleration and deceleration, 150-2000ms for peak velocity, and up to 4500ms for movement endpoints). All extracted variables—including PV, PA, PD and their respective timepoints (TPV, TPA, TPD), as well as movement endpoints (ME)—were saved as condition- and trial-specific .csv files. These files were subsequently imported into R for data visualization and statistical analysis.

### Statistical Analysis

All statistical analyses were performed using R (Version 4.0.3). Linear mixed-effects (LME) regression models were fitted using the lme4 package (Bates et al., 2015). To analyze the ME, PV, TPV, PA, TPA, PD, and TPD, we utilized the LME models which explicitly accounted for the variation in our data contributed to by trial and participant. Corrections for multiple comparisons were applied using Tukey-adjusted pairwise contrasts of estimated marginal means from the full linear mixed-effects model. For each combination of load condition, reach configuration, and force level, we tested whether the right and left hands differed significantly in performance. Positive values indicate greater or earlier values for the right hand, and negative values reflect a left-hand advantage. To further analyze right-left hand differences, we ran a second LME model using trial-wise difference scores (Right - Left) as the dependent variable. This model included load condition, reach symmetry, and force magnitude as fixed effects, with subject and trial as random intercepts. By collapsing across hand, this approach allowed us to directly assess how asymmetries varied systematically across task manipulations, independent of baseline hand effects. Corrections for multiple comparisons were applied using false discovery rate (FDR)-adjusted post hoc tests, which control the expected proportion of false positives among significant results (Chen et al., 2017). These analyses enabled both targeted and comprehensive evaluations of how force field perturbations modulated motor behavior during bimanual reaching.

## Results

### Kinematic Profiles of Reaching Movements

To characterize limb movement performance across conditions, we first visualized hand displacement, velocity, and acceleration profiles. Figures 2a–d present individual trial data overlaid with average kinematic trajectories for the no-load (baseline) condition. Full trajectories for all reaching movements and force field conditions are provided in the supplemental materials. These visualizations show how hand position (displacement), velocity, and acceleration evolved over time for both congruent and incongruent reaching movements. The displacement, velocity, and acceleration profiles for all other loaded conditions—viscous/viscous, elastic/elastic, no load/elastic, elastic/no load, no load/viscous, viscous/no load, viscous/elastic, and elastic/viscous—are shown in the supplementary figures. Low force conditions (Elastic = 30 N, Viscous = −15 N) are displayed in Supplementary Figures S1-S8, while medium force conditions (Elastic = 36.25 N, Viscous = −20 N) are shown in Supplementary Figures S9-S16.

**Figure 2.**
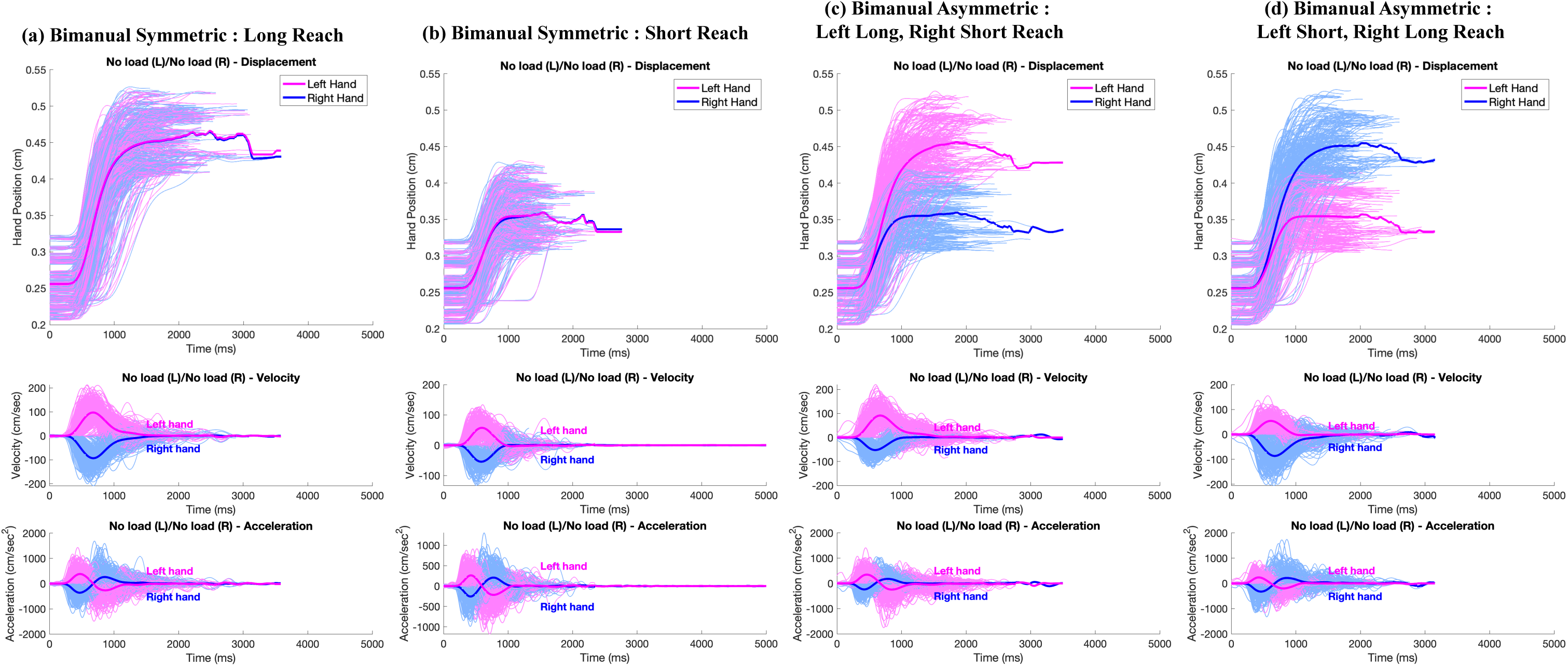
Kinematic Profiles. The displacement, velocity, and acceleration over time for the baseline (no load) condition across all four reaching movements are presented. Individual data points from all 30 participants are shown in light magenta and light blue for the left- and right-hand movements, respectively. The dark magenta and dark blue lines represent the averages of these individual data points for each hand.

### Reaction Time (RT), Movement Time (MT), Total Response Time (ResT)

To confirm that movement initiation and execution times were consistent across experimental conditions, we first analyzed reaction time (RT), movement time (MT), and total response time (ResT). We anticipated minimal differences; moreover, the task design emphasized reaching to the targets as quickly and accurately as possible, encouraging participants to balance speed and accuracy in a way that constrained movement variability despite differences in load and reaching conditions.

RT, defined as the interval from stimulus onset to movement initiation, showed no consistent effects of load conditions, hand or their interaction across either the symmetric or asymmetric reaching tasks under both low force (30N Elastic, −15N Viscous) and medium force (36.25N Elastic, −20N Viscous) conditions. Although a few isolated significant differences emerged for specific load combinations, these effects did not generalize across reach types or force levels, suggesting that they were likely incidental rather than systematic. MT, representation the duration between the initiation and completion of movement, similarly remained statistically stable across load conditions and hands. Finally, ResT, the sum of RT and MT, confirmed this pattern, with overall movement duration remaining comparable across all experimental manipulations. Occasional differences were observed but were small in magnitude and inconsistent across varying loads and reaching conditions.

Together, these results demonstrate that participants maintained synchronized timing of reach initiation and completion across all loads and reach conditions. Thus, any differences found in movement kinematics are unlikely to be caused by changes in overall movement timing but rather reflect true effects of the experimental manipulations on movement dynamics. Full statistical models and supplemental figures for RT, MT, and ResT are provided in the supplementary materials.

### Movement End Time (ME)

We first analyzed movement end time (ME), defined as the total time (in milliseconds) from stimulus onset to the moment when the participant’s hand first entered the target zone, as recorded by the Kinarm system when the hand crossed into the predefined acceptance radius for each trial. This measure captures when the hand crosses the predefined spatial endpoint threshold, signaling the completion of the reach. Figure 3a shows the total movement time to endpoint completion across all loads and reach configurations. To assess these effects, we fit a LME model with fixed effects of load configuration (Condition), hand (left or right), symmetry type (matched vs. mismatched reach), and force magnitude (low vs. medium), including the Condition x Hand interaction. Random intercepts were included for participant and trial number (Table 1). In addition, we calculated the difference in ME between the right and left hands for each trial and fit a second LME model using these difference scores as the dependent variable. This model output included fixed effects of load configuration (Condition), symmetry type (matched vs. mismatched reach), and force magnitude (low vs. medium). Figure 3b presents the results of this difference-based model, and the full statistical output is provided in supplementary table 10.

**Figure 3.**
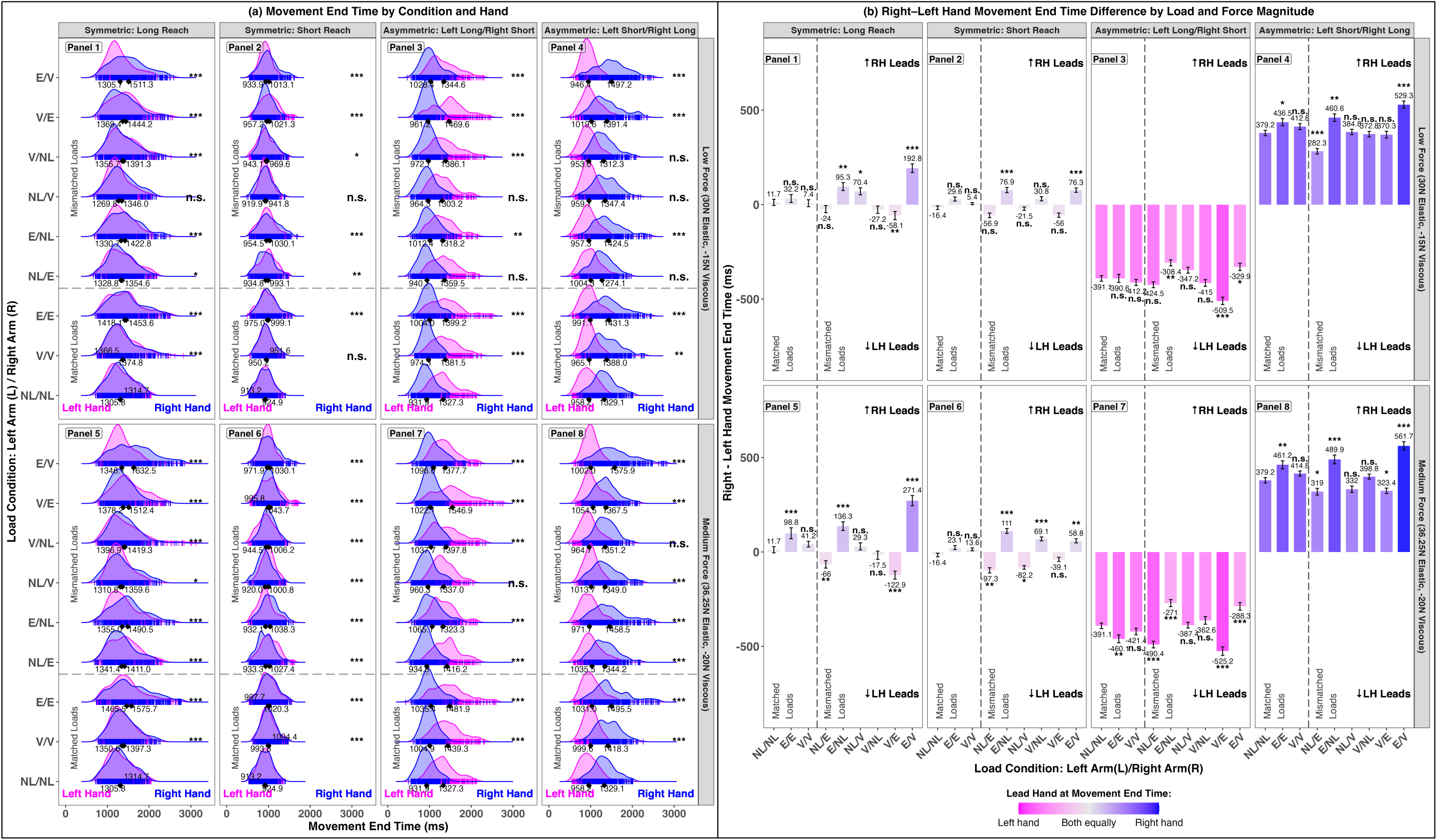
Movement End Time (ME) During Reaching Movements. (a) ME across load conditions and reach types. Panels 1-4 show symmetric long reach, symmetric short reach, asymmetric left-long/right-short reach, and asymmetric left-short/right-long reach under the low force condition (30N Elastic, −15N Viscous). Panels 5-8 show the same reach types under the medium force condition (36.25N Elastic, −20N Viscous). The gray dashed line separates matched and mismatched load conditions. Asterisks indicate statistical significance relative to the no load (NL/NL) baseline. Ridge density plots illustrate the distribution of ME values, with curves representing the overall shape of the data and vertical bars indicating individual trials. Diamonds mark the mean ME for each hand. (b) Right-left hand differences in ME. Panel layout matches (a). Positive values indicate a right-hand lead; negative values indicate a left-hand lead. Statistical significance is shown relative to the NL/NL baseline. The gradient legend represents movement end (ME) synchronization: bright pink indicates the left hand’s reach ended earlier, bright blue indicates the right hand’s reach ended earlier, and intermediate shades reflect more synchronized movement end times between hands.

**Table 1.** Results from the linear mixed-effects analysis of movement end time (ME) across symmetric and asymmetric bimanual reaching configurations. Fixed effects estimates (β), standard errors (SE), and p-values are reported. The model included random intercepts for participant and trial number and included fixed effects for Load Condition × Hand, Reach Configuration, and Force Magnitude. Significant effects (p < 0.05) are highlighted in bold.

|  | Linear Mixed-Effects Model Results: Movement End Time (ME) |  |  |
| --- | --- | --- | --- |
| Fixed effects<br>L=left arm/R=right arm) | $\beta$ | SE | $p(\chi^2)$ |
| Intercept<br>(No Load (L)/No Load (R)) | 1326.74 | 34.16 | <0.001 |
| Viscous (L)/Viscous (R) | 53.84 | 7.68 | <0.001 |
| Elastic (L)/ Elastic (R) | 91.36 | 7.69 | <0.001 |
| No Load (L)/Elastic (R) | 71.30 | 7.68 | <0.001 |
| Elastic (L)/No Load (R) | 12.95 | 7.69 | 0.092 |
| No Load (L)/Viscous (R) | 18.82 | 7.70 | 0.015 |
| Viscous (L)/No Load (R) | 42.76 | 7.65 | <0.001 |
| Viscous (L)/Elastic (R) | 135.39 | 7.68 | <0.001 |
| Elastic (L)/Viscous (R) | 27.28 | 7.69 | <0.001 |
| Hand [Right] | -5.05 | 7.63 | 0.508 |
| Reach Configuration [Symmetric Short Reach] | -418.48 | 3.64 | <0.001 |
| Reach Configuration [Asymmetric Left-Long/Right-Short Reach] | -201.84 | 3.64 | <0.001 |
| Reach Configuration [Asymmetric Left-Short/Right-Long Reach] | -200.30 | 3.64 | <0.001 |
| Force Magnitude [Medium Force: 36.25N Elastic, -20N Viscous] | 32.48 | 2.57 | <0.001 |
| Viscous (L)/Viscous (R) x Hand [Right] | 11.61 | 10.81 | 0.283 |
| Elastic (L)/ Elastic (R) x Hand [Right] | 40.47 | 10.83 | <0.001 |
| No Load (L)/Elastic (R) x Hand [Right] | -64.15 | 10.82 | <0.001 |
| Elastic (L)/No Load (R) x Hand [Right] | 106.45 | 10.82 | <0.001 |
| No Load (L)/Viscous (R) x Hand [Right] | 7.79 | 10.84 | 0.472 |
| Viscous (L)/No Load (R) x Hand [Right] | 8.23 | 10.78 | 0.445 |
| Viscous (L)/Elastic (R) x Hand [Right] | -76.98 | 10.81 | <0.001 |
| Elastic (L)/Viscous (R) x Hand [Right] | 149.93 | 10.84 | <0.001 |
| Random effects | Groups |  | SD |
|  | Trial Number | Intercept | 29.81 |
|  | Participant | Intercept | 184.05 |
|  | Residual |  | 259.67 |
|  | Observations: 41373 |  |  |
| Full Model: ME ~ (Load Condition x Hand) + Reach Configuration + Force Magnitude + (1 \| Trial Number) + (1 \| Participant) |  |  |  |

Overall, as shown in Table 1, the average time to reach movement end under baseline conditions (no load/no load) was 1326.74ms (SE = 34.16, p < .001). Among matched load conditions, viscous/viscous (53.83ms, SE = 7.68, p < .001) and elastic/elastic (91.36ms, SE = 7.69, p < .001) both showed significant increases relative to baseline, whereas among mismatched load conditions, no load/elastic (71.30ms, SE = 7.68, p < .001), no load/viscous (18.82ms, SE = 7.70, p = .015), viscous/no load (42.7ms, SE = 7.65, p < .001), viscous/elastic (135.39ms, SE = 7.68, p < .001), and elastic/viscous (27.28ms, SE = 7.69, p < .001) also showed significant increases; elastic/no load showed a smaller, non-significant trend (12.95ms, SE = 7.69, p = .092). This suggests that both matched and mismatched load conditions generally caused the total movement time to increase relative to the baseline (no load/no load), meaning it took participants longer to complete their reaches under most load manipulations. Symmetry type had a robust effect: symmetric short reaches were completed much faster (−418.48ms, SE = 3.64, p < .001), and both asymmetric reach types (left-long/right-short: −201.84ms, SE = 3.64, p < .001; left-short/right-long: −200.30ms, SE = 3.64, p < .001) also led to shorter movement end times relative to symmetric long reaches. Medium force produced a modest but reliable increase in duration (32.4ms, SE = 2.57, p < .001).

Importantly, condition x hand interactions revealed that load-related left-right differences were particularly pronounced under mismatched load conditions, with significant effects observed for no load/elastic x hand (−64.15ms, SE = 10.82, p < .001), elastic/no load x hand (106.45ms, SE = 10.82, p < .001), viscous/elastic x hand (76.98ms, SE = 10.81, p < .001), and elastic/viscous x hand (149.93ms, SE = 10.84, p < .001). Interestingly, even the matched condition elastic/elastic showed a significant interaction (40.47ms, SE = 10.83, p < .001), whereas other matched load interactions (viscous/viscous, no load/viscous, viscous/no load) were non-significant. Together, these results indicate that mismatched load conditions, in particular, amplify hand-specific adjustments in movement end timing, reflecting the motor system’s need to flexibly coordinate the timing demands of the two hands under unequal load constraints.

Lastly, while we report this reduced model for clarity, we also fit a full-factorial model including all Condition x Hand x Symmetry x Force interactions to generate estimated marginal means and visualize interaction effects. Post hoc pairwise comparisons between left and right hands were conducted within each load condition, reach type, and force level (Supplementary Table 11). These comparisons revealed significant timing asymmetries in several mismatched conditions, particularly under incongruent reach configurations, with temporal differences exceeding ±500ms in some cases. For a more targeted analysis of these asymmetries, we computed the difference in time to peak velocity between the right and left hands and fit a separate full-factorial mixed-effects model using Condition x Symmetry x Force as fixed effects (since Hand was already differenced). To assess whether the temporal degree of separation between the hands varied systematically across load conditions, we compared these right–left difference scores to the No Load/No Load (NL/NL) baseline (Supplementary Table 12). Positive or negative values reflect which hand led, while larger values indicate a greater temporal degree of separation. Several mismatched configurations produced significantly greater separations than NL/NL, especially when both reach symmetry and force magnitude were imbalanced. Together, these post-hoc analyses offer important detail on how particular load pairings shape bilateral coordination at the moment of movement completion.

### Peak Velocity (PV)

Figure 4a illustrates the peak velocities (PV) observed during reaching movements across all load and reach configurations. PV reflects the maximum hand speed achieved during the reach and provides insight into how force output adapts under varying mechanical demands. To assess differences across conditions, we fit a linear mixed-effects model with fixed effects of load configuration (Condition), hand (left or right), symmetry type (symmetric vs. asymmetric reach), and force magnitude (low vs. medium), including the Condition x Hand interaction. Random intercepts were included for subject and trial (Table 2a). In addition, we calculated the difference in PV between the right and left hands for each trial and fit a second LME model using these difference scores as the dependent variable. This model output included fixed effects of load configuration (Condition), symmetry type (matched vs. mismatched reach), and force magnitude (low vs. medium). Figure 4b presents the results of this difference-based model, and the full statistical output is provided in supplementary table 13a.

**Figure 4.**
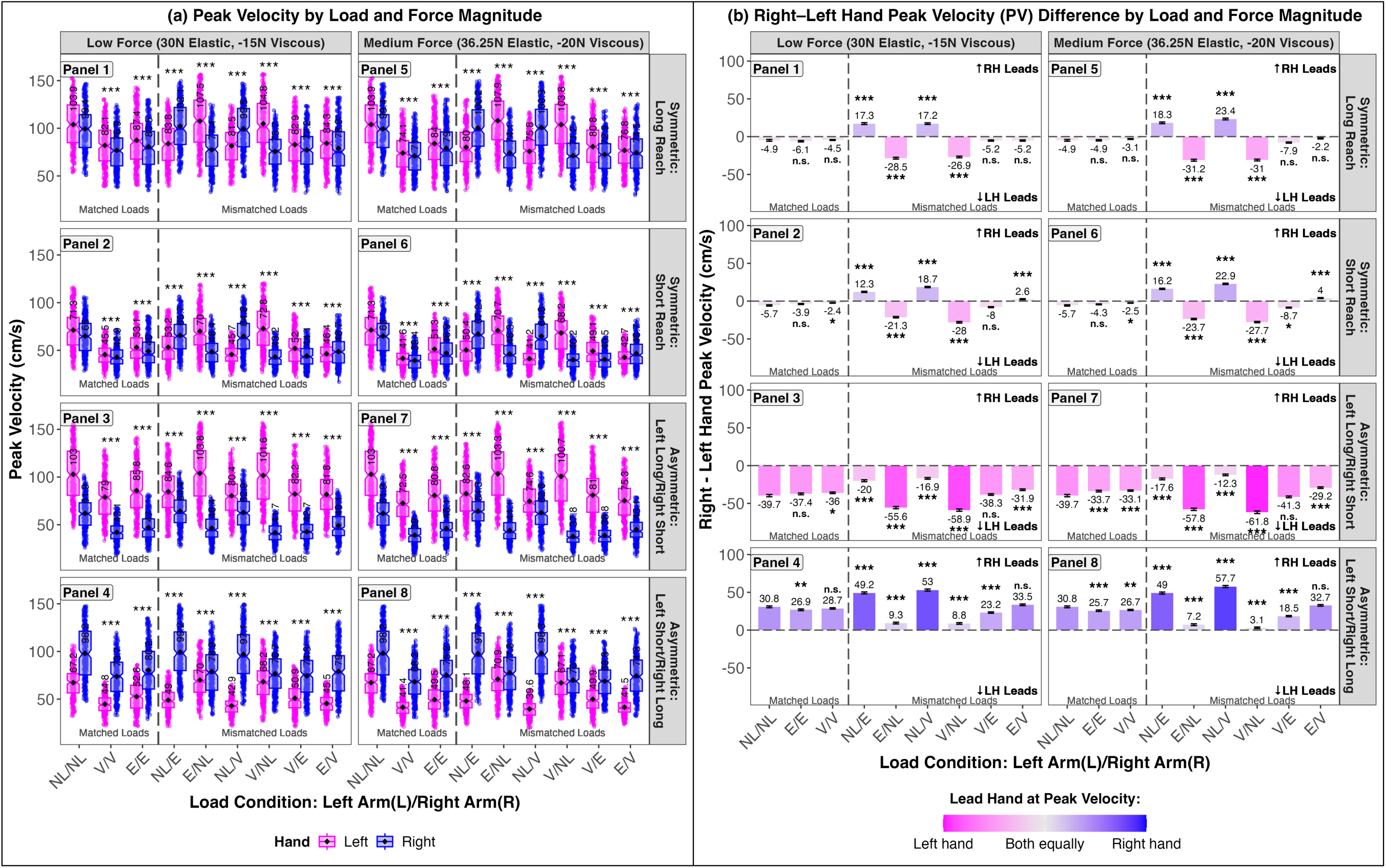
Peak Velocity (PV) During Reaching Movements. (a) PV across load conditions and reach types. Panels 1-4 show symmetric long reach, symmetric short reach, asymmetric left-long/right-short reach, and asymmetric left-short/right-long reach under the low force condition (30N Elastic, −15N Viscous). Panels 5-8 show the same reach types under the medium force condition (36.25N Elastic, −20N Viscous). The gray dashed line separates matched from mismatched load conditions. Asterisks indicate statistical significance relative to the no load (NL/NL) baseline. Boxplots include notches for medians, solid black dots for means, and whiskers for the full data range. (b) Right-left hand differences in peak velocity. Panels match the layout in (a). Positive values indicate a right-hand lead; negative values indicate a left-hand lead. Statistical differences are shown relative to the NL/NL baseline. The gradient legend reflects PV synchronization: bright pink indicates stronger left-hand involvement, bright blue indicates stronger right-hand involvement, and intermediate shades reflect more synchronized contributions from both hands.

**Table 2.** Results from the linear mixed-effects analysis of (a) peak velocity (PV) and (b) timepoint at peak velocity (TPV) across symmetric and asymmetric bimanual reaching configurations. Fixed effects estimates (β), standard errors (SE), and p-values are reported. The model included random intercepts for participant and trial number and included fixed effects for Load Condition × Hand, Reach Configuration, and Force Magnitude. Significant effects (p < 0.05) are highlighted in bold.

|  | Linear Mixed-Effects Model Results: PV & TPV |  |  |  |  |  |
| --- | --- | --- | --- | --- | --- | --- |
|  | (a) Peak Velocity (PV) |  |  | (b) Timepoint at Peak Velocity (TPV) |  |  |
| Fixed effects<br>L=left arm/R=right arm) | $\beta$ | SE | $p(\chi^2)$ | $\beta$ | SE | $p(\chi^2)$ |
| Intercept<br>(No Load (L)/No Load (R)) | 108.32 | 2.72 | <0.001 | 721.37 | 17.81 | <0.001 |
| Viscous (L)/Viscous (R) | -27.81 | 0.54 | <0.001 | 195.21 | 4.62 | <0.001 |
| Elastic (L)/ Elastic (R) | -20.05 | 0.54 | <0.001 | 48.30 | 4.62 | <0.001 |
| No Load (L)/Elastic (R) | -19.85 | 0.55 | <0.001 | 54.69 | 4.68 | <0.001 |
| Elastic (L)/No Load (R) | -0.04 | 0.55 | 0.935 | 7.15 | 4.67 | 0.126 |
| No Load (L)/Viscous (R) | -26.51 | 0.55 | <0.001 | 195.48 | 4.66 | <0.001 |
| Viscous (L)/No Load (R) | -2.39 | 0.54 | <0.001 | 4.64 | 4.66 | 0.319 |
| Viscous (L)/Elastic (R) | -21.89 | 0.54 | <0.001 | 59.49 | 4.62 | <0.001 |
| Elastic (L)/Viscous (R) | -26.18 | 0.54 | <0.001 | 194.03 | 4.62 | <0.001 |
| Hand [Right] | -5.23 | 0.55 | <0.001 | 0.68 | 5.17 | 0.896 |
| Reach Configuration [Symmetric Short Reach] | -34.16 | 0.25 | <0.001 | -130.20 | 1.92 | <0.001 |
| Reach Configuration [Asymmetric Left-Long/Right-Short Reach] | -18.42 | 0.26 | <0.001 | -49.83 | 1.96 | <0.001 |
| Reach Configuration [Asymmetric Left-Short/Right-Long Reach] | -18.91 | 0.26 | <0.001 | -52.02 | 1.96 | <0.001 |
| Force Magnitude [Medium Force: 36.25N Elastic, -20N Viscous] | -2.69 | 0.18 | <0.001 | 17.06 | 1.37 | <0.001 |
| Viscous (L)/Viscous (R) x Hand [Right] | 1.76 | 0.76 | 0.021 | -4.09 | 6.49 | 0.529 |
| Elastic (L)/ Elastic (R) x Hand [Right] | 0.56 | 0.76 | 0.465 | -7.38 | 6.49 | 0.256 |
| No Load (L)/Elastic (R) x Hand [Right] | 20.03 | 0.77 | <0.001 | -61.71 | 6.57 | <0.001 |
| Elastic (L)/No Load (R) x Hand [Right] | -19.24 | 0.77 | <0.001 | 41.03 | 6.56 | <0.001 |
| No Load (L)/Viscous (R) x Hand [Right] | 25.16 | 0.77 | <0.001 | -202.50 | 6.55 | <0.001 |
| Viscous (L)/No Load (R) x Hand [Right] | -21.97 | 0.77 | <0.001 | 188.01 | 6.54 | <0.001 |
| Viscous (L)/Elastic (R) x Hand [Right] | -3.17 | 0.76 | <0.001 | 130.45 | 6.50 | <0.001 |
| Elastic (L)/Viscous (R) x Hand [Right] | 5.62 | 0.76 | <0.001 | -148.25 | 6.49 | <0.001 |
| Random effects | Groups |  | SD | Groups |  | SD |
|  | Trial Number | Intercept | 1.70 | Trial Number | Intercept | 12.51 |
|  | Participant | Intercept | 14.72 | Participant | Intercept | 95.09 |
|  | Residual |  | 17.72 | Residual |  | 132.55 |
|  | Observations: 39775 |  |  | Observations: 38159 |  |  |
| Full Model: PV or TPV ~ (Load Condition x Hand) + Reach Configuration + Force Magnitude + (1 \| Trial Number) + (1 \| Participant) |  |  |  |  |  |  |

The model—seen in Table 2a— revealed that PV was significantly influenced by load configuration. Compared to the neutral no load/no load baseline, both matched-load conditions (viscous/viscous: −27.81cm/s, SE = 0.54, p < .001; elastic/elastic: −20.05cm/s, SE = 0.54, p < .001) and several mismatched-load conditions (no load/elastic: −19.85cm/s, SE = 0.55, p < .001; no load/viscous: −26.51cm/s, SE = 0.55, p < .001; viscous/elastic: −21.89cm/s, SE = 0.54, p < .001; elastic/viscous: −26.18cm/s, SE = 0.54, p < .001) produced significant reductions in PV. Elastic/no load showed no significant difference (−0.04cm/s, SE = 0.55, p = .935), while viscous/no load showed a small but reliable reduction (−2.39cm/s, SE = 0.54, p < .001). Symmetry type exerted a strong influence: symmetric short-reach movements were significantly faster than symmetric long-reach trials (−34.16cm/s, SE = 0.25, p < .001), and both asymmetric reaches—left-long/right-short (−18.42cm/s, SE = 0.26, p < .001) and left-short/right-long (−18.91cm/s, SE = 0.26, p < .001)—also produced faster peak velocities. Additionally, medium force trials introduced a small but significant slowing compared to low force (−2.69 cm/s, SE = 0.18, p < .001).

A main effect of hand revealed that right-hand reaches were slightly slower overall (−5.23cm/s, SE = 0.55, p < .001). Several significant condition x hand interactions emerged, indicating that lateralized effects were condition specific. Notably, the viscous/viscous matched-load condition showed a small but significant right-hand advantage (1.76cm/s, SE = 0.76, p = .021), while the elastic/elastic matched-load condition showed no significant hand difference (0.56cm/s, SE = 0.76, p = .465). In contrast, all mismatched-load conditions showed robust hand-dependent effects: under no load/elastic, the right hand was substantially faster (20.03cm/s, p < .001), while under elastic/no load, the right hand was significantly slower (−19.24cm/s, p < .001). Similar asymmetries were observed in no load/viscous (25.16cm/s), viscous/no load (−21.97cm/s), viscous/elastic (−3.17cm/s), and elastic/viscous (5.62cm/s), all p < .001.

In summary, peak velocity was shaped by load configuration, symmetry type, and their interaction with handedness. Both matched and mismatched loads imposed distinct costs on force output, but mismatched-load conditions triggered particularly strong hand-dependent asymmetries, highlighting the adaptive role of lateralization under mechanical imbalance. Symmetric short reaches yielded the fastest peak velocities, while medium force slightly reduced maximum movement speed. Together, these findings reveal the complex interplay between biomechanical demands and sensorimotor control in regulating force production.

Lastly, while we report this reduced model for clarity, we also fit a full-factorial model including all Condition x Hand x Symmetry x Force interactions to generate estimated marginal means and visualize interaction effects. Post-hoc pairwise comparisons between left and right hands were conducted within each load condition, reach type, and force level (Supplementary Table 14). These comparisons revealed significant peak velocity asymmetries in several mismatched conditions, particularly under asymmetric reach configurations. In many of these cases, the right hand exhibited a consistent performance advantage, with differences exceeding ±50 cm/s. For a more targeted analysis of these asymmetries, we computed the difference in peak velocity between the right and left hands and fit a separate full-factorial mixed-effects model using Condition x Symmetry x Force as fixed effects (since Hand was already differenced). To assess whether the degree of separation in peak velocity between the hands varied systematically across load conditions, we compared these right-left difference scores to the No Load/No Load (NL/NL) baseline (Supplementary Table 15). Positive or negative values reflect which hand achieved higher velocity, while larger values indicate a greater degree of performance asymmetry. Several mismatched configurations produced significantly greater asymmetries than NL/NL, especially when both reach symmetry and force magnitude were imbalanced. In summary, the post-hoc results highlight that although left/right differences appear across several conditions, the most robust and meaningful lateralized effects are driven by mismatched load pairings, with matched-load asymmetries emerging only under specific conditions involving higher force magnitudes.

### Timepoint at Peak Velocity (TPV)

Figure 5 illustrates the timepoint at which peak velocity (TPV) occurred during reaching movements across all load and reach configurations. TPV reflects the moment of maximum hand speed and provides insight into how movement timing adapts under varying mechanical demands. To assess differences across conditions, we fit a linear mixed-effects model with fixed effects of load configuration (Condition), hand (left or right), symmetry type (symmetric vs. asymmetric reach), and force magnitude (low vs. medium), including the Condition x Hand interaction (Table 2b). Random intercepts were included for subject and trial. To further assess asymmetries, we computed right-left TPV differences on each trial and fit a second model using these difference scores as the dependent variable. This model included Condition, reach symmetry, and force magnitude as fixed effects. Figure 5b summarizes the results of this difference-based model, with full statistical output available in Supplementary Table 13b.

**Figure 5.**
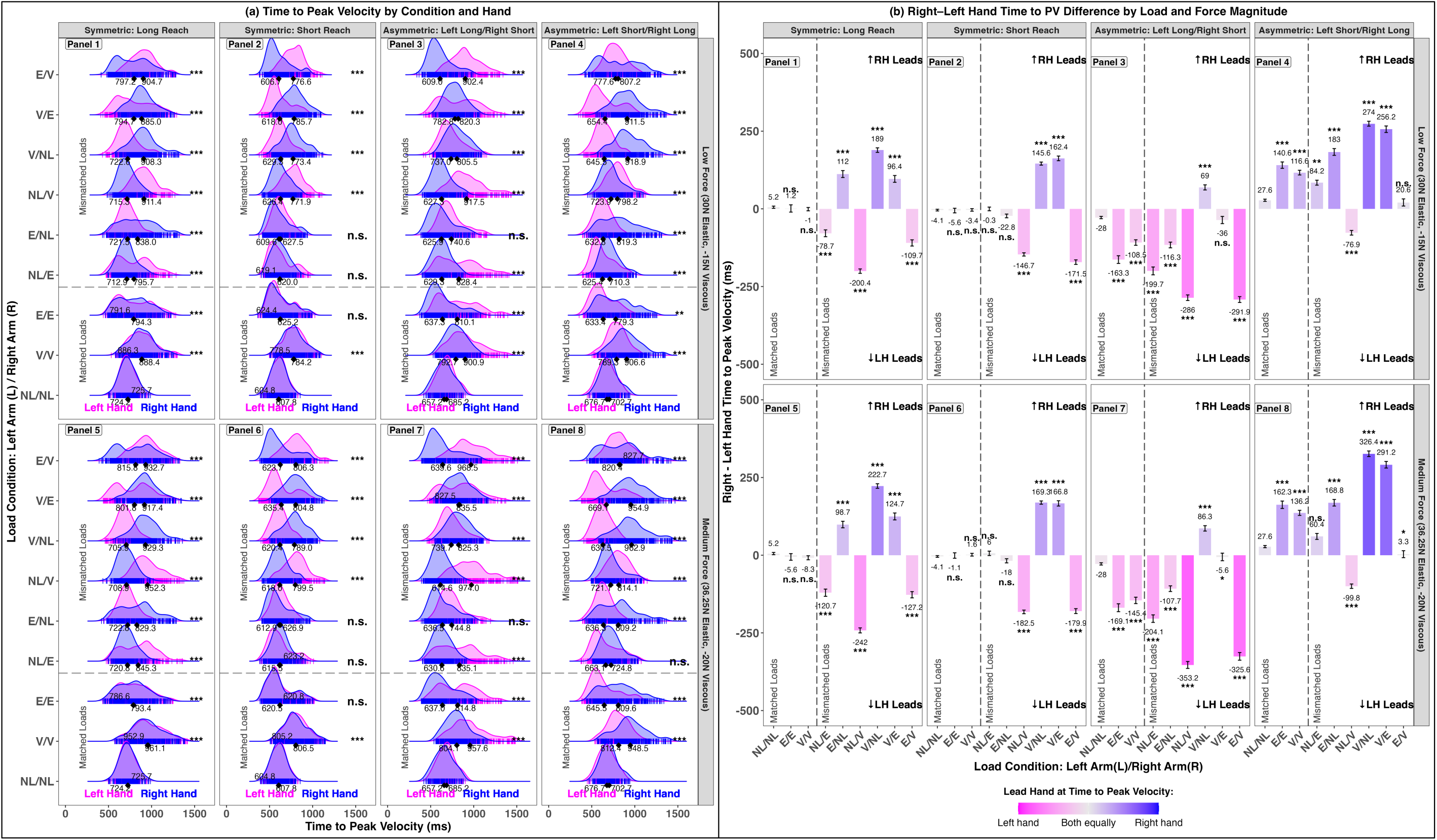
Time at Peak Velocity (TPV) During Reaching Movements. (a) TPV across load conditions and reach types. Panels 1-4 show symmetric long reach, symmetric short reach, asymmetric left-long/right-short reach, and asymmetric left-short/right-long reach under the low force condition (30N Elastic, −15N Viscous). Panels 5-8 show the same reach types under the medium force condition (36.25N Elastic, −20N Viscous). The gray dashed line separates matched and mismatched load conditions. Asterisks indicate statistical significance relative to the no load (NL/NL) baseline. Ridge density plots illustrate the distribution of TPV values, with curves representing the overall shape of the data and vertical bars indicating individual trials. Diamonds mark the mean TPV for each hand. (b) Right-left hand differences in TPV. Panel layout matches (a). Positive values indicate a right-hand lead; negative values indicate a left-hand lead. Statistical significance is shown relative to the NL/NL baseline. The gradient legend reflects TPV synchronization: bright pink indicates greater left-hand contribution, bright blue indicates greater right-hand contribution, and intermediate shades represent more synchronized timing between the hands.

The model—seen in table 2b—revealed that TPV was significantly influenced by load configuration. Compared to the no load/no load baseline, both matched-load conditions (viscous/viscous: 195.21ms, SE = 4.62, p < .001; elastic/elastic: 48.30ms, SE = 4.62, p < .001) and several mismatched load conditions (no load/elastic: 54.69ms, SE = 4.68, p < .001; no load/viscous: 195.47ms, SE = 4.66, p < .001; viscous/elastic: 59.49ms, SE = 4.62, p < .001; elastic/viscous: 194.03ms, SE = 4.62, p < .001) produced significant increases in TPV. Elastic/no load (7.15ms, SE = 4.67, p = .126) and viscous/no load (4.64ms, SE = 4.66, p = .319) did not significantly differ from baseline. Symmetry type exerted a strong influence: symmetric short-reach movements were significantly faster than symmetric long-reach trials (−130.20ms, SE = 1.92, p < .001), and both asymmetric reaches—left-long/right-short (−49.83ms, SE = 1.96, p < .001) and left-short/right-long (−52.02ms, SE = 1.96, p < .001)—also led to faster movements. Additionally, medium force trials introduced a modest but reliable increase in TPV compared to low force (17.06ms, SE = 1.37, p < .001).

While no main effect of hand (right vs. left) was observed (0.68ms, SE = 5.17, p = .896), several significant condition x hand interactions emerged. Notably, the interactions showed that in matched-load conditions (viscous/viscous x hand, elastic/elastic x hand), there were no significant differences relative to baseline. In contrast, all mismatched-load conditions (e.g., no load/elastic, no load/viscous, viscous/elastic, elastic/viscous, elastic/no load, viscous/no load) showed significant hand-dependent effects, indicating that lateralization only emerged under asymmetric force pairings. For example, the right hand was faster under no load/elastic (−61.71ms, SE = 6.57, p < .001) and elastic/viscous (−148.25ms, SE = 6.49, p < .001), but slower in no load/viscous (202.50ms, SE = 6.55, p < .001) and viscous/no load (188.00ms, SE = 6.54, p < .001), demonstrating clear condition-specific shifts.

In summary, time to peak velocity was shaped by load configuration, symmetry type, and their interaction with handedness. Both matched and mismatched loads imposed distinct timing costs, but only the mismatched-load conditions triggered significant hand-dependent asymmetries, revealing the adaptive role of lateralization under load imbalance. Symmetric short reaches yielded the fastest movements, while medium force produced a modest slowing effect. Together, these findings highlight the complex interplay between load demands and sensorimotor control in shaping movement timing.

Lastly, while we report this reduced model for clarity, we also fit a full-factorial model including all Condition x Hand x Symmetry x Force interactions to generate estimated marginal means and visualize interaction effects. Post hoc pairwise comparisons between left and right hands were conducted within each load condition, reach type, and force level (Supplementary Table 16). These comparisons revealed substantial timing asymmetries in multiple mismatched conditions, with temporal differences often exceeding ±300ms, particularly under asymmetric reach configurations. To more directly assess these asymmetries, we computed the difference in time to peak velocity between the right and left hands and fit a separate full-factorial mixed-effects model using Condition x Symmetry x Force as fixed effects. This model treated hand as a differenced variable, capturing the temporal lead of one hand over the other. To determine whether the temporal degree of separation between the hands varied systematically across load conditions, we compared these right-left difference scores to the No Load/No Load (NL/NL) baseline (Supplementary Table 17). Positive or negative values indicated which hand led, while larger absolute values reflected a greater temporal separation. Several mismatched load configurations resulted in significantly greater temporal separation than NL/NL, especially when both reach symmetry and loads applied to each hand were mismatched. In many of these conditions, the right hand consistently reached peak velocity earlier than the left, suggesting a reliable temporal coordination bias. These post-hoc results provide key insight into how load distribution and biomechanical constraints shape bilateral motor timing at the moment of peak movement execution.

### Peak Acceleration (PA)

Figure 6a illustrates the peak acceleration (PA) achieved during reaching movements across all load and reach configurations. PA reflects the maximum rate of change in hand velocity, offering insight into how participants scaled their force production under varying load conditions. To assess differences across conditions, we fit a linear mixed-effects model with fixed effects of load configuration (Condition), hand (left or right), symmetry type (symmetric vs. asymmetric reach), and force magnitude (low vs. medium), including the Condition x Hand interaction (Table 3a). Random intercepts were included for subject and trial. To further explore differences between the hands, we computed right-minus-left PA difference scores on each trial and fit a second model using these values as the outcome. This model included Condition, reach symmetry, and force magnitude as fixed effects. Figure 6b summarizes the results of this difference-based model; full statistical output is provided in Supplementary Table 18a.

**Figure 6.**
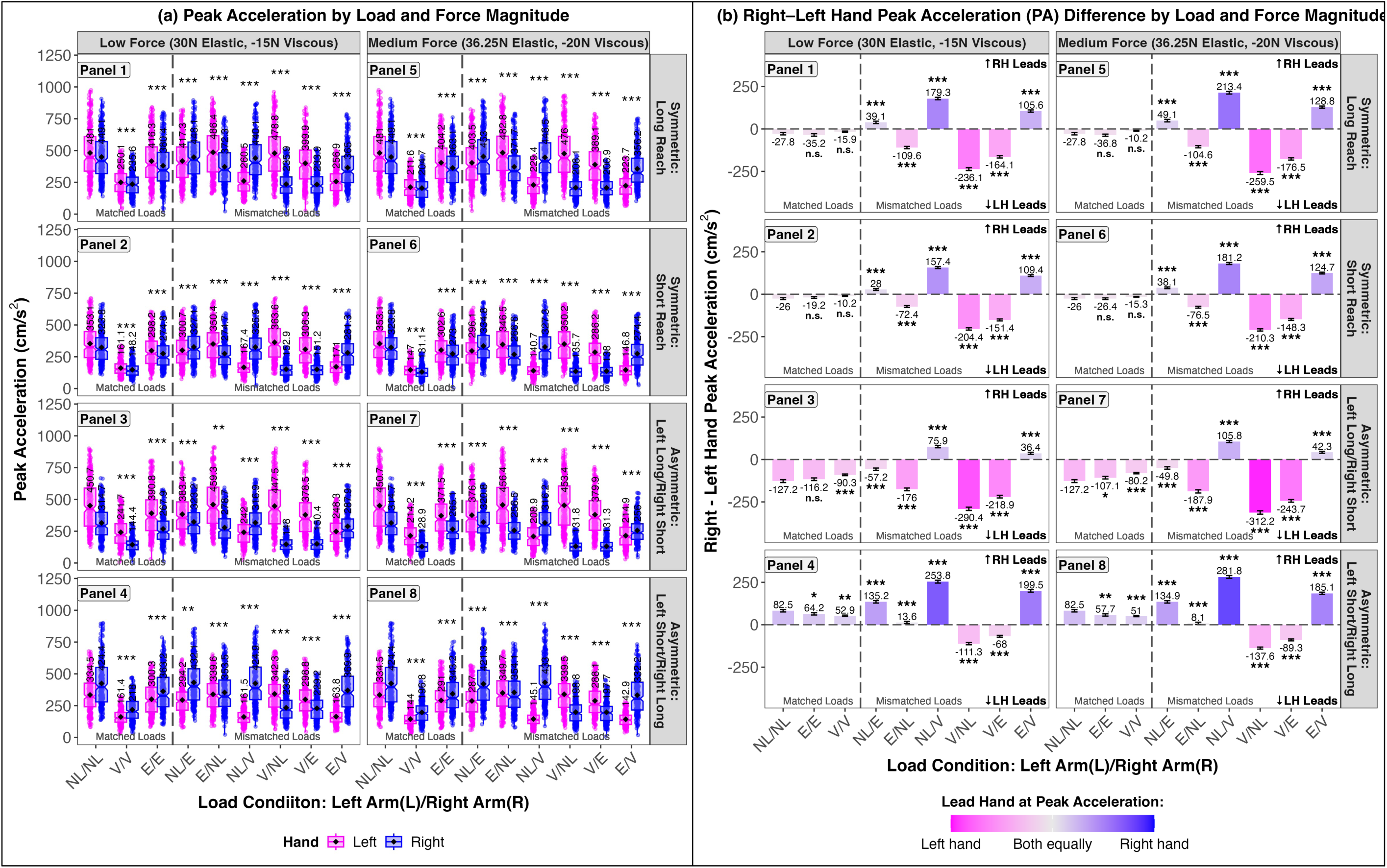
Peak Acceleration (PA) During Reaching Movements. (a) PA across load conditions and reach types. Panels 1-4 show symmetric long reach, symmetric short reach, asymmetric left-long/right-short reach, and asymmetric left-short/right-long reach under the low force condition (30N Elastic, −15N Viscous). Panels 5-8 show the same reach types under the medium force condition (36.25N Elastic, −20N Viscous). The gray dashed line separates matched from mismatched load conditions. Asterisks indicate statistical significance relative to the no load (NL/NL) baseline. Boxplots include notches for medians, solid black dots for means, and whiskers for the full data range. (b) Right-left hand differences in peak acceleration. Panels match the layout in (a). Positive values indicate a right-hand lead; negative values indicate a left-hand lead. Statistical differences are shown relative to the NL/NL baseline. The gradient legend reflects PA synchronization: bright pink indicates stronger left-hand involvement, bright blue indicates stronger right-hand involvement, and intermediate shades reflect more synchronized contributions from both hands.

**Table 3.** Results from the linear mixed-effects analysis of (a) peak acceleration (PA) and (b) timepoint at peak acceleration (TPA) across symmetric and asymmetric bimanual reaching configurations. Fixed effects estimates (β), standard errors (SE), and p-values are reported. The model included random intercepts for participant and trial number and included fixed effects for Load Condition × Hand, Reach Configuration, and Force Magnitude. Significant effects (p < 0.05) are highlighted in bold.

|  | Linear Mixed-Effects Model Results: PA & TPA |  |  |  |  |  |
| --- | --- | --- | --- | --- | --- | --- |
|  | (a) Peak Acceleration (PA) |  |  | (b) Timepoint at Peak Acceleration (TPA) |  |  |
| Fixed effects<br>L=left arm/R=right arm) | $\beta$ | SE | $p(\chi^2)$ | $\beta$ | SE | $p(\chi^2)$ |
| Intercept<br>(No Load (L)/No Load (R)) | 478.25 | 18.99 | <0.001 | 514.02 | 10.48 | <0.001 |
| Viscous (L)/Viscous (R) | -213.51 | 2.79 | <0.001 | 71.66 | 2.85 | <0.001 |
| Elastic (L)/ Elastic (R) | -60.02 | 2.78 | <0.001 | 0.23 | 2.88 | 0.937 |
| No Load (L)/Elastic (R) | -61.74 | 2.78 | <0.001 | -4.64 | 2.88 | 0.107 |
| Elastic (L)/No Load (R) | 0.90 | 2.78 | 0.745 | -3.80 | 2.86 | 0.183 |
| No Load (L)/Viscous (R) | -210.27 | 2.79 | <0.001 | 74.12 | 2.85 | <0.001 |
| Viscous (L)/No Load (R) | -1.73 | 2.78 | 0.533 | -4.40 | 2.86 | 0.123 |
| Viscous (L)/Elastic (R) | -64.55 | 2.78 | <0.001 | 10.54 | 2.88 | <0.001 |
| Elastic (L)/Viscous (R) | -208.98 | 2.79 | <0.001 | 77.13 | 2.85 | <0.001 |
| Hand [Right] | -24.54 | 2.78 | <0.001 | -8.55 | 2.84 | 0.003 |
| Reach Configuration [Symmetric Short Reach] | -100.14 | 1.32 | <0.001 | -59.85 | 1.35 | <0.001 |
| Reach Configuration [Asymmetric Left-Long/Right-Short Reach] | -61.45 | 1.32 | <0.001 | -29.98 | 1.35 | <0.001 |
| Reach Configuration [Asymmetric Left-Short/Right-Long Reach] | -64.09 | 1.32 | <0.001 | -27.24 | 1.35 | <0.001 |
| Force Magnitude [Medium Force: 36.25N Elastic, -20N Viscous] | -11.71 | 0.93 | <0.001 | 9.01 | 0.96 | <0.001 |
| Viscous (L)/Viscous (R) x Hand [Right] | 10.11 | 3.92 | 0.010 | 3.13 | 4.02 | 0.436 |
| Elastic (L)/ Elastic (R) x Hand [Right] | -2.68 | 3.92 | 0.494 | -1.77 | 4.05 | 0.662 |
| No Load (L)/Elastic (R) x Hand [Right] | 63.99 | 3.92 | <0.001 | 1.69 | 4.03 | 0.675 |
| Elastic (L)/No Load (R) x Hand [Right] | -64.06 | 3.92 | <0.001 | -5.04 | 4.04 | 0.212 |
| No Load (L)/Viscous (R) x Hand [Right] | 209.57 | 3.92 | <0.001 | -78.30 | 4.02 | <0.001 |
| Viscous (L)/No Load (R) x Hand [Right] | -197.33 | 3.92 | <0.001 | 91.89 | 4.02 | <0.001 |
| Viscous (L)/Elastic (R) x Hand [Right] | -134.71 | 3.92 | <0.001 | 74.96 | 4.03 | <0.001 |
| Elastic (L)/Viscous (R) x Hand [Right] | 144.87 | 3.92 | <0.001 | -84.22 | 4.04 | <0.001 |
| Random effects | Groups |  | SD | Groups |  | SD |
|  | Trial Number | Intercept | 11.07 | Trial Number | Intercept | 7.66 |
|  | Participant | Intercept | 103.32 | Participant | Intercept | 56.07 |
|  | Residual |  | 93.33 | Residual |  | 96.21 |
|  | Observations: 40925 |  |  | Observations: 40894 |  |  |
| Full Model: PA or TPA ~ (Load Condition x Hand) + Reach Configuration + Force Magnitude + (1 \| Trial Number) + (1 \| Participant) |  |  |  |  |  |  |

The model—shown in table 3a—revealed that PA was significantly influenced by load condition. Compared to the neutral no load/no load baseline, both matched-load conditions (viscous/viscous: −213.51cm/s^2^, SE = 2.79, p < .001; elastic/elastic: −60.02cm/s^2^, SE = 2.78, p < .001) and several mismatched load conditions (no load/elastic: −61.74cm/s^2^, SE = 2.78, p < .001; no load/viscous: −210.27cm/s^2^, SE = 2.79, p < .001; viscous/elastic: −64.55cm/s^2^, SE = 2.78, p < .001; elastic/viscous: −208.98cm/s^2^, SE = 2.79, p < .001) produced substantial reductions in PA. Elastic/no load and viscous/no load did not significantly differ from baseline (both p > .5). Symmetry type also had a strong effect: symmetric short reaches produced significantly greater accelerations (−100.14cm/s^2^, SE = 1.32, p < .001), while asymmetric configurations—left-long/right-short (−61.45cm/s^2^, SE = 1.32, p < .001) and left-short/right-long (−64.09cm/s^2^, SE = 1.32, p < .001)—also led to higher accelerations relative to symmetric long reaches. Additionally, medium-force trials introduced a small but reliable decrease in PA (−11.71cm/s^2^, SE = 0.93, p < .001). A main effect of hand showed that right hand reaches were slightly slower overall (−24.54cm/s^2^, SE = 2.78, p < .001). Several significant condition x hand interactions emerged, indicating that acceleration differences between hands depended on the load configuration. For example, the viscous/viscous symmetric matched-load condition showed a small but significant right-hand advantage (10.11cm/s^2^, SE = 3.92, p = .010), while the elastic/elastic condition showed no significant hand difference (−2.68cm/s^2^, p = .494). In contrast, mismatched load conditions produced large and reliable hand differences: no load/elastic favored the right hand (63.99cm/s^2^, p < .001), whereas elastic/no load favored the left (−64.06cm/s^2^, p < .001). Similar asymmetric effects appeared in no load/viscous (209.57cm/s^2^, p < .001), viscous/no load (−197.33cm/s^2^, p < .001), viscous/elastic (−134.71cm/s^2^, p < .001), and elastic/viscous (144.87 cm/s^2^, p < .001).

In summary, peak acceleration was shaped by load condition, symmetry type, force magnitude, and hand-specific effects that depended on the applied load configuration. While both matched and mismatched loads introduced distinct acceleration costs, the most pronounced and consistent hand dependent asymmetries emerged under mismatched load conditions, highlighting how the system adjusts limb-specific control when managing unequal load conditions. Importantly, force magnitude—particularly medium force—significantly reduced acceleration, reflecting the added difficulty of moving the hand under increased elastic, viscous, or combined loads. Together, these findings demonstrate how task related constraints systematically affect the motor system’s ability to accelerate quickly, with movement initiation slowing as load demands increase.

Lastly, while we report this reduced model for clarity, we also fit a full-factorial model including all Condition x Hand x Symmetry x Force interactions to generate estimated marginal means and visualize interaction effects. Post-hoc pairwise comparisons between left and right hands were conducted within each load condition, reach type, and force level (Supplementary Table 19). These comparisons revealed significant temporal asymmetries in many mismatched conditions, with differences in time to peak acceleration often exceeding ±250cm/s^2^. In many of these cases, the right hand reached peak acceleration earlier than the left. To further assess these asymmetries, we computed right-minus-left difference scores for time to peak acceleration on each trial and fit a second mixed-effects model using Condition x Symmetry x Force as fixed effects. This model allowed us to assess whether the temporal degree of separation between hands systematically varied by condition. We then compared these difference scores to the No Load/No Load (NL/NL) baseline (Supplementary Table 20). Several mismatched configurations produced significantly greater temporal separation than NL/NL, particularly when both reach symmetry and loads were mismatched. Together, these results suggest that the coordination of bilateral force initiation is highly sensitive to load pairing and task asymmetry.

### Timepoint at Peak Acceleration (TPA)

Figure 7a illustrates the timepoint at peak acceleration (TPA) achieved during reaching movements across all load and reach configurations. TPA reflects the time at which the hand reaches its maximum acceleration, providing insight into how quickly the motor system ramps up force under varying load conditions. To assess differences across conditions, we fit a linear mixed-effects model with fixed effects of load configuration (Condition), hand (left or right), symmetry type (symmetric vs. asymmetric reach), and force magnitude (low vs. medium), including the Condition x Hand interaction (Table 3b). Random intercepts were included for participant and trial number. To further characterize interlimb differences, we calculated right-minus-left TPA values on each trial and fit a second model using these difference scores as the dependent variable. This model included condition, reach symmetry, and force magnitude as fixed effects. Figure 7b summarizes the results of this model, with full statistical details provided in Supplementary Table 18b.

**Figure 7.**
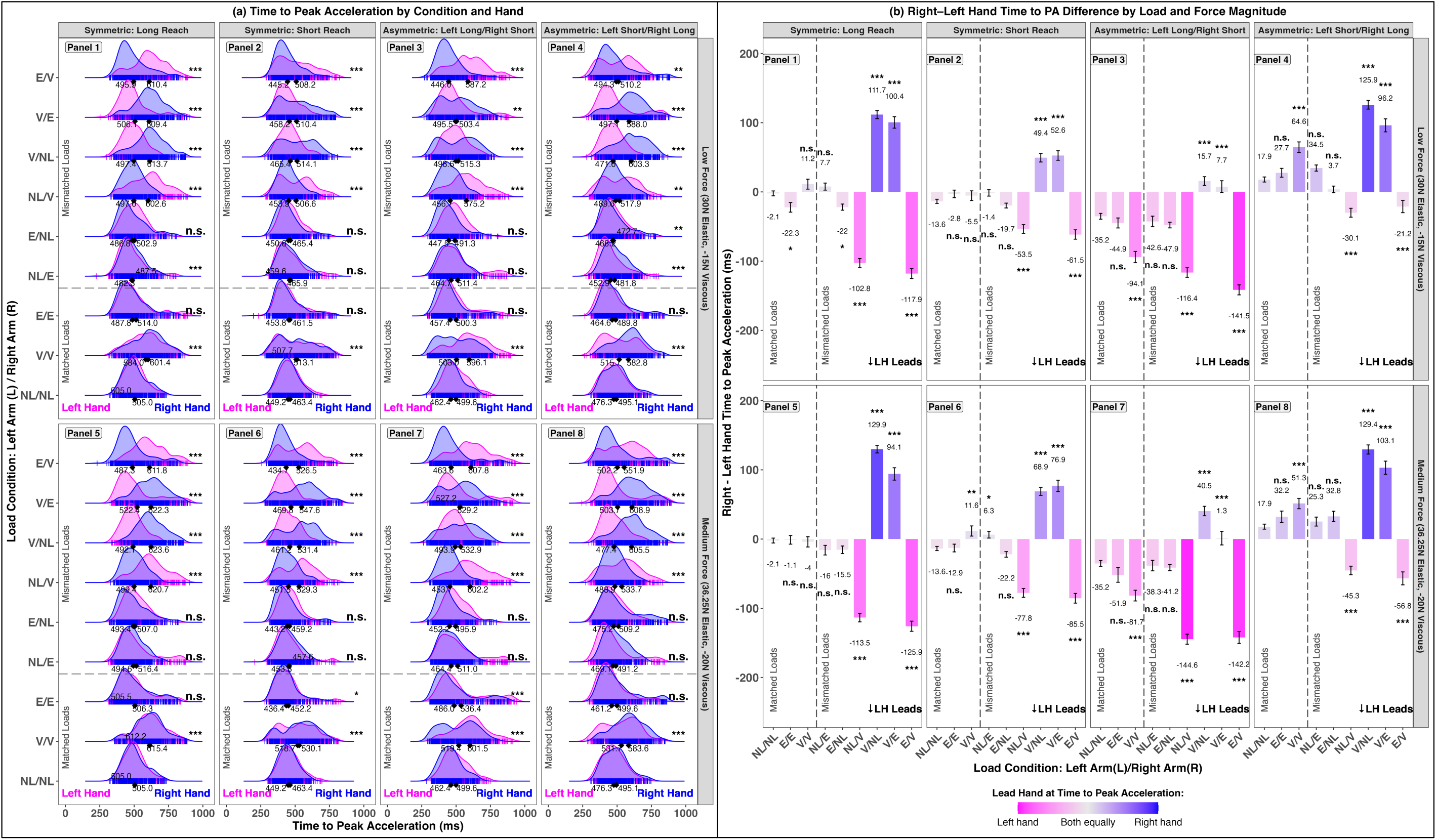
Time at Peak Acceleration (TPA) During Reaching Movements. (a) TPA across load conditions and reach types. Panels 1-4 show symmetric long reach, symmetric short reach, asymmetric left-long/right-short reach, and asymmetric left-short/right-long reach under the low force condition (30N Elastic, −15N Viscous). Panels 5-8 show the same reach types under the medium force condition (36.25N Elastic, −20N Viscous). The gray dashed line separates matched and mismatched load conditions. Asterisks indicate statistical significance relative to the no load (NL/NL) baseline. Ridge density plots illustrate the distribution of TPA values, with curves representing the overall shape of the data and vertical bars indicating individual trials. Diamonds mark the mean TPA for each hand. (b) Right-left hand differences in TPA. Panel layout matches (a). Positive values indicate a right-hand lead; negative values indicate a left-hand lead. Statistical significance is shown relative to the NL/NL baseline. The gradient legend reflects TPA synchronization: bright pink indicates greater left-hand contribution, bright blue indicates greater right-hand contribution, and intermediate shades represent more synchronized timing between the hands.

The model—showed in table 3b—showed that TPA was significantly influenced by load conditions. Compared to the no load/no load condition, matched load conditions produced mixed effects: viscous/viscous significantly increased TPA (71.66ms, SE = 2.85, p < .001), while elastic/elastic showed no reliable difference (0.23ms, SE = 2.88, p = .937). Several mismatched-load configurations, including no load/viscous (74.12ms, SE = 2.85, p < .001), viscous/elastic (10.54ms, SE = 2.88, p < .001), and elastic/viscous (77.13ms, SE = 2.85, p < .001), significantly prolonged TPA, whereas no load/elastic, elastic/no load, and viscous/no load showed no significant change relative to baseline (all p > .10). These findings indicate that the timing of peak acceleration is selectively modulated by specific load conditions, with mismatched and certain matched loads imposing greater temporal demands.

Symmetry type strongly shaped TPA: congruent short-reach trials (−59.85ms, SE = 1.35, p < .001), incongruent left-long/right-short trials (−29.98ms, SE = 1.35, p < .001), and incongruent left-short/right-long trials (−27.24ms, SE = 1.35, p < .001) all led to earlier peak acceleration compared to symmetric long reaches. Medium force magnitude trials also modestly increased TPA (9.01ms, SE = 0.96, p < .001), indicating slower force buildup under greater force magnitude. Although the main effect of hand (right vs. left) was small (−8.55ms, SE = 2.84, p = .003), the condition x hand interactions revealed substantial asymmetries under mismatched loads. For example, no load/viscous x hand (−78.30ms, SE = 4.02, p < .001), viscous/no load x hand (91.89ms, SE = 4.02, p < .001), viscous/elastic x hand (74.96ms, SE = 4.03, p < .001), and elastic/viscous x hand (−84.22ms, SE = 4.04, p < .001) all showed robust and significant hand-specific effects, highlighting the motor system’s adaptive lateralization when handling mismatched loads.

Lastly, while we report this reduced model for clarity, we also fit a full-factorial model including all Condition x Hand x Symmetry x Force interactions to generate estimated marginal means and visualize interaction effects. Post-hoc pairwise comparisons between left and right hands were conducted within each load condition, reach type, and force level (Supplementary Table 21). These comparisons revealed significant timing asymmetries in conditions involving mismatched load pairings and asymmetric load configurations, with differences in time to peak acceleration frequently exceeding ±100ms. The largest discrepancies were observed in mismatched load pairings such as V/NL and NL/V, with left–right differences surpassing ±250ms in several cases. In many of these conditions, the right hand consistently reached peak acceleration earlier than the left.

For a more targeted analysis of these asymmetries, we computed the difference in time to peak acceleration between the right and left hands and fit a separate full-factorial mixed-effects model using Condition x Symmetry x Force as fixed effects (since Hand was already differenced). To assess whether the temporal degree of separation between the hands varied systematically across load conditions, we compared these difference scores to the No Load/No Load (NL/NL) baseline (Supplementary Table 22). Positive or negative values reflect which hand led, while larger values indicate a greater temporal degree of separation. Several mismatched configurations produced significantly greater separations than NL/NL, particularly under conditions involving both asymmetric reach configurations and unequal force requirements. Together, these findings highlight that while moderate asymmetries appear across conditions, the most robust and reliable hand differences in TPA emerge under mismatched-load conditions, reinforcing the idea that lateralization is a key adaptive mechanism when the motor system faces asymmetric load challenges.

### Peak Deceleration (PD)

Figure 8a illustrates the peak deceleration (PD) across all load and reach configurations. To assess differences across conditions, we fit a linear mixed-effects model with fixed effects of load configuration (Condition), hand (left or right), symmetry type (symmetric vs. asymmetric reach), and force magnitude (low vs. medium), including the Condition x Hand interaction. Random intercepts were included for participant and trial number (Table 4a). PD reflects the maximum rate of deceleration during movement, offering insight into how the motor system modulates force output and adjusts movement dynamics in response to varying load conditions and reach configurations. To further assess between-hand differences, we computed right-minus-left PD values for each trial and fit a second model using these difference scores as the dependent variable. This model included Condition, reach symmetry, and force magnitude as fixed effects. Figure 8b presents a summary of these results, with full statistical output available in Supplementary Table 23a.

**Figure 8.**
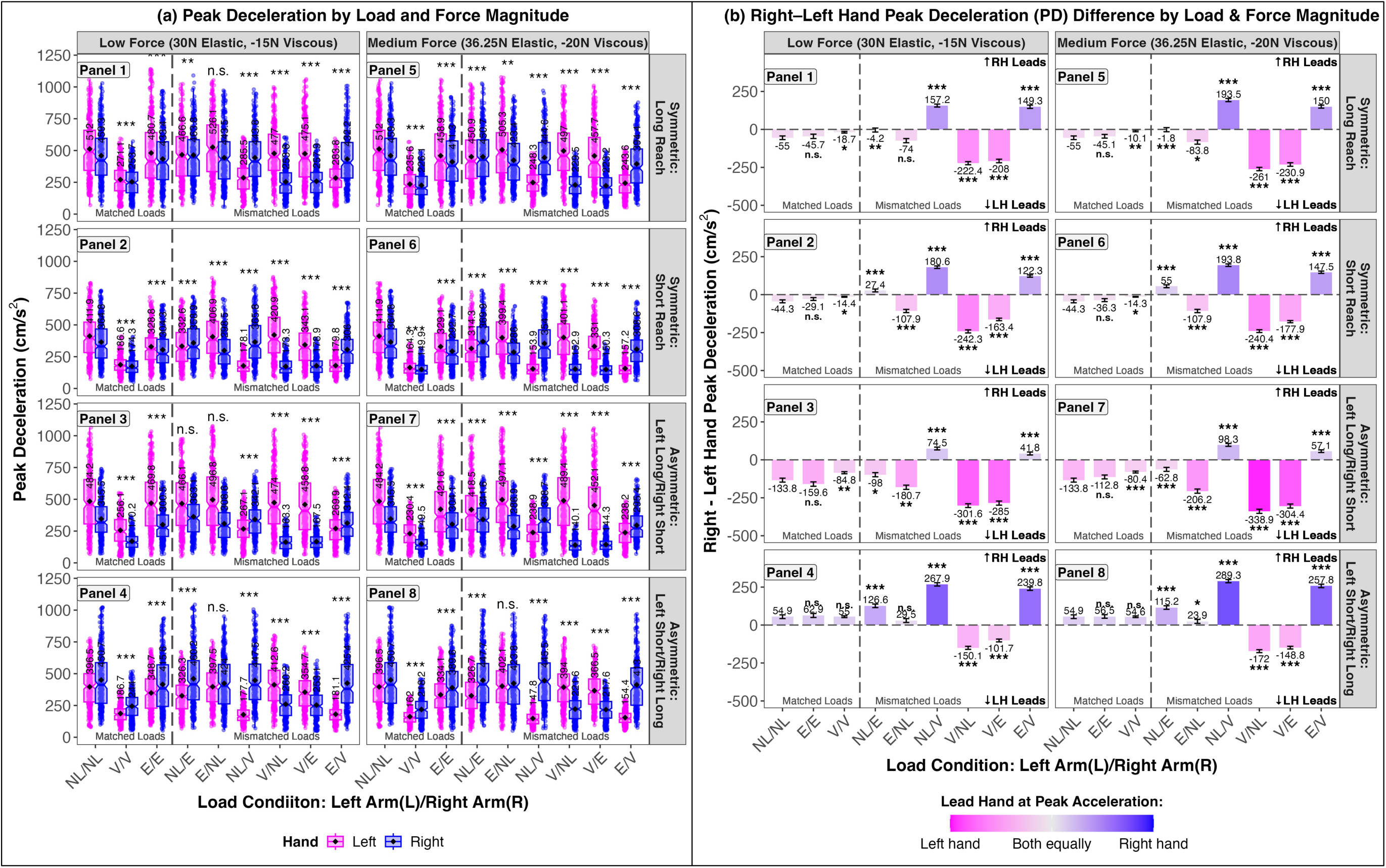
Peak Deceleration (PD) During Reaching Movements. (a) PD across load conditions and reach types. Panels 1-4 show symmetric long reach, symmetric short reach, asymmetric left-long/right-short reach, and asymmetric left-short/right-long reach under the low force condition (30N Elastic, −15N Viscous). Panels 5-8 show the same reach types under the medium force condition (36.25N Elastic, −20N Viscous). The gray dashed line separates matched from mismatched load conditions. Asterisks indicate statistical significance relative to the no load (NL/NL) baseline. Boxplots include notches for medians, solid black dots for means, and whiskers for the full data range. (b) Right-left hand differences in peak deceleration. Panels match the layout in (a). Positive values indicate a right-hand lead; negative values indicate a left-hand lead. Statistical differences are shown relative to the NL/NL baseline. The gradient legend reflects PD synchronization: bright pink indicates stronger left-hand involvement, bright blue indicates stronger right-hand involvement, and intermediate shades reflect more synchronized contributions from both hands.

**Table 4.** Results from the linear mixed-effects analysis of (a) peak deceleration (PD) and (b) timepoint at peak deceleration (TPD) across symmetric and asymmetric bimanual reaching configurations. Fixed effects estimates (β), standard errors (SE), and p-values are reported. The model included random intercepts for participant and trial number and included fixed effects for Load Condition × Hand, Reach Configuration, and Force Magnitude. Significant effects (p < 0.05) are highlighted in bold.

|  | Linear Mixed-Effects Model Results: PD & TPD |  |  |  |  |  |
| --- | --- | --- | --- | --- | --- | --- |
|  | (a) Peak Deceleration (PD) |  |  | (b) Timepoint at Peak Deceleration (TPD) |  |  |
| Fixed effects<br>L=left arm/R=right arm) | $\beta$ | SE | $p(\chi^2)$ | $\beta$ | SE | $p(\chi^2)$ |
| Intercept<br>(No Load (L)/No Load (R)) | 518.47 | 18.03 | <0.001 | 961.89 | 21.09 | <0.001 |
| Viscous (L)/Viscous (R) | -243.81 | 4.13 | <0.001 | 237.08 | 5.21 | <0.001 |
| Elastic (L)/ Elastic (R) | -53.78 | 4.05 | <0.001 | 84.18 | 5.11 | <0.001 |
| No Load (L)/Elastic (R) | -62.30 | 4.05 | <0.001 | 82.40 | 5.11 | <0.001 |
| Elastic (L)/No Load (R) | 2.35 | 4.05 | 0.561 | -1.96 | 5.15 | 0.703 |
| No Load (L)/Viscous (R) | -241.87 | 4.10 | <0.001 | 230.43 | 5.19 | <0.001 |
| Viscous (L)/No Load (R) | -9.93 | 4.09 | 0.015 | 7.77 | 5.18 | 0.133 |
| Viscous (L)/Elastic (R) | -49.14 | 4.09 | <0.001 | 97.48 | 5.16 | <0.001 |
| Elastic (L)/Viscous (R) | -240.44 | 4.09 | <0.001 | 239.87 | 5.19 | <0.001 |
| Hand [Right] | -42.73 | 4.03 | <0.001 | 1.73 | 5.12 | 0.735 |
| Reach Configuration [Symmetric Short Reach] | -98.61 | 1.94 | <0.001 | -166.89 | 2.44 | <0.001 |
| Reach Configuration [Asymmetric Left-Long/Right-Short Reach] | -60.33 | 1.95 | <0.001 | -66.34 | 2.45 | <0.001 |
| Reach Configuration [Asymmetric Left-Short/Right-Long Reach] | -56.77 | 1.95 | <0.001 | -66.79 | 2.45 | <0.001 |
| Force Magnitude [Medium Force: 36.25N Elastic, -20N Viscous] | -16.76 | 1.37 | <0.001 | 20.36 | 1.73 | <0.001 |
| Viscous (L)/Viscous (R) x Hand [Right] | 29.24 | 5.81 | <0.001 | -4.27 | 7.34 | 0.561 |
| Elastic (L)/ Elastic (R) x Hand [Right] | 3.55 | 5.71 | 0.534 | -1.64 | 7.18 | 0.820 |
| No Load (L)/Elastic (R) x Hand [Right] | 63.34 | 5.71 | <0.001 | -93.22 | 7.23 | <0.001 |
| Elastic (L)/No Load (R) x Hand [Right] | -46.51 | 5.71 | <0.001 | 89.94 | 7.22 | <0.001 |
| No Load (L)/Viscous (R) x Hand [Right] | 228.43 | 5.77 | <0.001 | -235.50 | 7.31 | <0.001 |
| Viscous (L)/No Load (R) x Hand [Right] | -200.93 | 5.76 | <0.001 | 223.80 | 7.29 | <0.001 |
| Viscous (L)/Elastic (R) x Hand [Right] | -162.00 | 5.76 | <0.001 | 138.61 | 7.28 | <0.001 |
| Elastic (L)/Viscous (R) x Hand [Right] | 191.20 | 5.77 | <0.001 | -151.00 | 7.28 | <0.001 |
| Random effects | Groups |  | SD | Groups |  | SD |
|  | Trial Number | Intercept | 12.54 | Trial Number | Intercept | 16.80 |
|  | Participant | Intercept | 97.20 | Participant | Intercept | 113.40 |
|  | Residual |  | 136.51 | Residual |  | 172.40 |
|  | Observations: 40109 |  |  | Observations: 40346 |  |  |
| Full Model: PD or TPD ~ (Load Condition x Hand) + Reach Configuration + Force Magnitude + (1 \| Trial Number) + (1 \| Participant) |  |  |  |  |  |  |

The model in Table 4a revealed that matched load conditions (e.g., viscous/viscous: −243.81cm/s^2^, SE = 4.15, p < .001; elastic/elastic: −53.78cm/s^2^, SE = 4.03, p < .001) and mismatched load conditions (e.g., no load/elastic: −62.30cm/s^2^, SE = 4.04, p < .001; no load/viscous: −241.87cm/s^2^, SE = 4.10, p < .001; viscous/no load: −9.93cm/s^2^, SE = 4.06, p = .015) significantly influenced peak deceleration compared to the no load/no load baseline. Notably, mismatched load pairings such as no load/viscous, viscous/no load, and elastic/no load showed distinct deceleration adjustments, suggesting heightened modulation demands under mismatched load conditions. There was a significant main effect of hand, with the right hand showing overall reduced PD (−42.73cm/s^2^, SE = 3.99, p < .001). Asymmetric reach configurations (e.g., incongruent left-long/right-short or left-short/right-long) further decreased PD (e.g., −60.33cm/s^2^, SE = 1.80, p < .001; −56.77cm/s^2^, SE = 1.82, p < .001), indicating task-specific adaptations. Medium force magnitude also reduced PD compared to low force (−16.76cm/s^2^, SE = 1.37, p < .001). These results suggest that differences between the hands and reach configurations systematically shape how the motor system modulates deceleration, reflecting adjustments specifically tuned to the load conditions and coordination demands of the task, rather than general or fixed patterns of control.

Importantly, the condition x hand interaction showed that some load pairings produced significant lateralized differences in peak deceleration. Specifically, the viscous/viscous x hand interaction was significant (29.24cm/s^2^, SE = 5.69, p < .001), indicating that even under symmetric viscous loading, left-right differences in deceleration control emerged. In contrast, the elastic/elastic x hand interaction was not significant (3.55cm/s^2^, SE = 5.66, p = .534), suggesting no meaningful lateralization when both hands operated under matched elastic loads. Furthermore, all mismatched load conditions showed robust and highly significant condition x hand interactions: no load/elastic x hand (63.34cm/s^2^, SE = 5.66, p < .001), elastic/no load x hand (−46.51cm/s^2^, SE = 5.66, p < .001), no load/viscous x hand (228.43cm/s^2^, SE = 5.72, p < .001), viscous/no load x hand (−200.93 m/s^2^, SE = 5.69, p < .001), and elastic/viscous x hand (191.20cm/s^2^, SE = 5.71, p < .001). These findings indicate that deceleration control is especially sensitive to mismatched load conditions, amplifying left-right differences. This pattern suggests that the motor system flexibly adjusts deceleration dynamics in a load-specific and lateralized manner, reflecting the demands of coordinating interlimb differences when faced with asymmetric load conditions.

Lastly, to maintain consistency with previous analyses, we fit a full-factorial model including all Condition x Hand x Symmetry x Force interactions to generate estimated marginal means and visualize interaction effects for peak deceleration (PD). Post-hoc pairwise comparisons between left and right hands were conducted within each load condition, reach type, and force level (Supplementary Table 24). These comparisons revealed significant PD asymmetries in a wide range of mismatched conditions, particularly under incongruent reach configurations and asymmetric force pairings, with differences exceeding ±250cm/s^2^ in several cases. Notably, the right hand frequently exhibited a consistent performance advantage in peak deceleration. This pattern was especially evident when the right hand was loaded, and the left hand was not. Conversely, when the left hand was loaded and the right hand was not, PD responses were often weaker or showed less modulation, reinforcing a broader asymmetry in how the two hands contribute to movement deceleration, suggesting that the left hand may be less effective at stopping movement precisely when it has to work harder—possibly due to reduced force control or coordination under mismatched loading.

To further quantify these asymmetries, we computed right-left difference scores for each trial and fit a separate mixed-effects model using Condition x Symmetry x Force as fixed effects (with Hand already differenced). These scores were referenced to the No Load/No Load (NL/NL) baseline to evaluate systematic deviations across experimental conditions (Supplementary Table 25). Positive or negative values indicate which hand decelerated more forcefully, while larger magnitudes reflect greater bilateral divergence in terminal motor control. Several mismatched configurations produced significantly greater separations than the matched baseline, particularly when both reach symmetry and force magnitude were imbalanced. These results demonstrate that peak deceleration, like other aspects of movement during bimanual reaching, is sensitive to how load and task demands are distributed across the hands.

### Timepoint at Peak Deceleration (TPD)

Figure 9a presents the timepoint at peak deceleration (TPD) across all load and reach configurations. TPD reflects the moment when the hand reaches maximum deceleration, marking the shift from speeding up to slowing down, and indicating how the motor system regulates deceleration forces under varying load conditions. We analyzed these data using a linear mixed-effects model with fixed effects for load configuration (Condition), hand (left or right), symmetry type (symmetric vs. asymmetric reach), force magnitude (low vs. medium), and their interactions, including random intercepts for participant and trial number (Table 4). To further evaluate interlimb differences, we computed right-minus-left TPD times on each trial and fit a second model using these difference scores as the dependent variable. This model included condition, symmetry, and force as fixed effects. Figure 9b summarizes these results, with full statistical output provided in Supplementary Table 23b.

**Figure 9.**
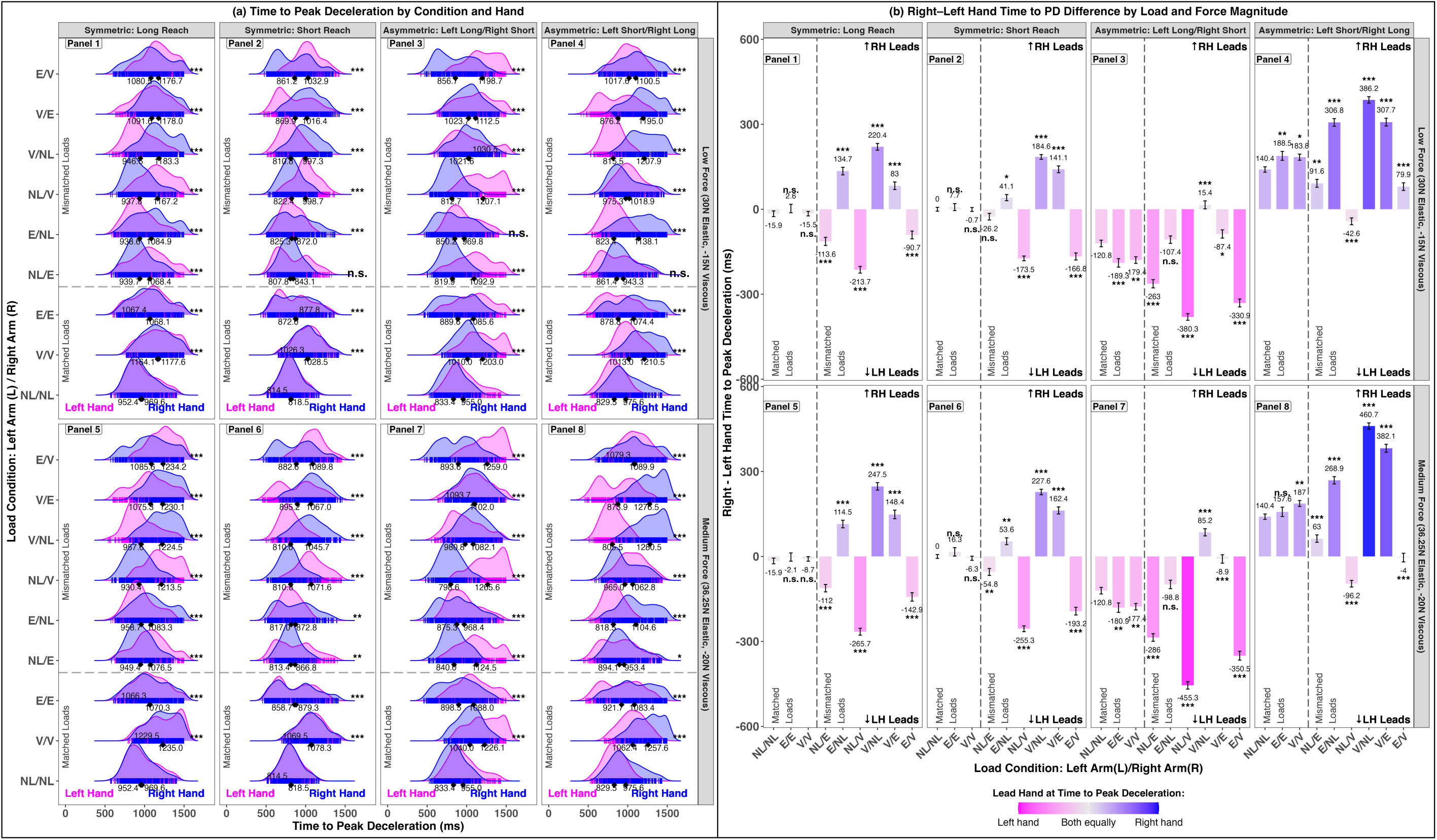
Time at Peak Deceleration (TPD) During Reaching Movements. (a) TPD across load conditions and reach types. Panels 1-4 show symmetric long reach, symmetric short reach, asymmetric left-long/right-short reach, and asymmetric left-short/right-long reach under the low force condition (30N Elastic, −15N Viscous). Panels 5-8 show the same reach types under the medium force condition (36.25N Elastic, −20N Viscous). The gray dashed line separates matched and mismatched load conditions. Asterisks indicate statistical significance relative to the no load (NL/NL) baseline. Ridge density plots illustrate the distribution of TPD values, with curves representing the overall shape of the data and vertical bars indicating individual trials. Diamonds mark the mean TPD for each hand. (b) Right-left hand differences in TPD. Panel layout matches (a). Positive values indicate a right-hand lead; negative values indicate a left-hand lead. Statistical significance is shown relative to the NL/NL baseline. The gradient legend reflects TPD synchronization: bright pink indicates greater left-hand contribution, bright blue indicates greater right-hand contribution, and intermediate shades represent more synchronized timing between the hands.

The model in Table 4b revealed that TPD was strongly modulated by load condition. Compared to the no load/no load baseline, matched load configurations such as viscous/viscous (237.08ms, SE = 5.21, p < .001) and elastic/elastic (84.18ms, SE = 5.11, p < .001) significantly delayed the timing of peak deceleration. Several mismatched load conditions, including no load/elastic (82.41ms, SE = 5.11, p < .001), viscous/elastic (97.48ms, SE = 5.16, p < .001), and elastic/viscous (239.87ms, SE = 5.19, p < .001), produced similarly large delays, whereas others like elastic/no load (−1.96ms, SE = 5.15, p = .703) and viscous/no load (7.77ms, SE = 5.18, p = .133) showed no reliable change, suggesting that not all mismatched-load pairings equally affect deceleration timing, and only certain combinations create enough challenge to alter when the motor system slows the movement. This highlights that the system adjusts deceleration timing mainly when the load asymmetry is strong enough to demand it, helping maintain control under more difficult conditions.

Symmetry type exerted a clear influence: congruent short-reach trials (−166.89ms, SE = 2.44, p < .001), incongruent left-long/right-short (−66.34ms, SE = 2.45, p < .001), and incongruent left-short/right-long (−66.79ms, SE = 2.45, p < .001) all led to earlier peak deceleration compared to symmetric long reaches. Medium force magnitude further increased TPD by ∼20ms (SE = 1.73, p < .001), reflecting the additional time required for movement deceleration under increased load. Interestingly, there was no significant main effect of hand (1.73ms, SE = 5.12, p = .735), but the condition x hand interactions revealed striking lateralized differences under mismatched loads. Specifically, no load/elastic (−93.22ms, SE = 7.23, p < .001), elastic/no load (89.94ms, SE = 7.22, p < .001), no load/viscous (−235.50ms, SE = 7.31, p < .001), viscous/no load (223.80ms, SE = 7.29, p < .001), viscous/elastic (138.61ms, SE = 7.28, p < .001), and elastic/viscous (−151.00ms, SE = 7.28, p < .001) produced robust, consistent hand asymmetries, underscoring the adaptive role of lateralization when the motor system compensates for mismatched load conditions.

Finally, we analyzed the TPD to assess how bilateral load pairing modulates temporal aspects of movement termination. As with prior measures, we first fit a linear mixed-effects model including fixed effects of condition, hand, symmetry, and force, along with their interactions. Post-hoc comparisons were used to evaluate differences in TPD between the right and left hands across conditions (Supplementary Table 26). While no significant lateralization was observed in the matched (NL/NL) baseline, numerous asymmetric configurations—especially those involving mismatched loading or asymmetric reach configurations —exhibited large, statistically significant hand differences. In some cases, differences between hands exceeded ±450ms, with the right hand more frequently reaching peak deceleration earlier under mismatched load conditions. To quantify these effects more directly, we computed right-minus-left TPD difference scores for each trial and fit a second mixed-effects model with Condition, Symmetry, and Force as fixed effects (Supplementary Table 27). Comparisons relative to the NL/NL baseline confirmed that mismatched load conditions—including V/NL, NL/V, and E/NL—produced reliably larger temporal deviations than matched load conditions. These findings indicate that while minor asymmetries exist across conditions, the most pronounced and consistent lateralized differences in timepoint at peak deceleration appear when the motor system must adapt to mismatched load conditions across the two hands.

## Discussion

Bimanual movements pose a fundamental challenge for the motor system: how to coordinate two limbs that may face different demands. While foundational research has focused on temporal (Kelso et al., 1979) and spatial (Kelso et al., 1983) coupling during symmetrical or mirror-symmetric tasks (Marteniuk et al., 1984; Spijkers & Heuer, 1995; Diedrichsen et al., 2004; Heuer & and Klein, 2006), relatively few studies have explored how the motor system responds when asymmetries—such as applying viscous or elastic load—are introduced to each limb (Brunfeldt et al., 2022). In this study, we examined how load conditions (matched vs. mismatched), force magnitude (low vs. medium), and reach configuration (symmetric vs. asymmetric target distances) shape bimanual reaching when participants are instructed to move both hands simultaneously. As predicted and based on prior findings (Kelso et al., 1979; Diedrichsen et al., 2001; Bingham et al., 2008), reaction time (RT) remained stable across conditions, indicating that participants initiated both hands together regardless of load type, reach configurations, or force magnitude. However, during movement execution, we observed clear effects of load condition on movement time (MT) and total response time (ResT). Mismatched load conditions (e.g., no load/viscous, viscous/no load, no load/elastic, elastic/no load, elastic/viscous, viscous/elastic) reliably increased MT and ResT, reflecting the motor system’s need to adjust movement duration to accommodate different load demands on each limb. Notably, these increases in ResT were driven by execution-phase adjustments, not planning delays, suggesting that participants compensated during movement to maintain synchrony at the target despite facing mismatched load conditions.

Beyond RT, MT, and ResT, movement end time (ME) was especially sensitive to load condition and reach configuration. Matched loads (elastic/elastic, viscous/viscous) produced modest but reliable increases in ME compared to no load/no load, with elastic/elastic showing larger delays than viscous/viscous. This suggests that position-dependent elastic loads require more adjustment near the endpoint than velocity-dependent viscous loads. Mismatched load conditions (e.g., no load/viscous, viscous/no load, elastic/viscous) led to even greater increases in ME, highlighting the added coordination challenge when each limb faces different load demands. Notably, participants may have used a hover phase strategy—briefly slowing or pausing the less-loaded, faster limb near the target—to allow the more-loaded, slower limb to catch up, maintaining endpoint synchrony despite different movement dynamics (Riek et al., 2003; Srinivasan & Martin, 2010; Coats & Wann, 2012; Sardar et al., 2023). We also found strong effects of reach configuration: symmetric short reaches and asymmetric reaches (left-long/right-short, left-short/right-long) consistently reduced ME compared to symmetric long reaches, showing that ME is strongly driven by target distance. Finally, force magnitude (low vs. medium) also increased ME slightly across conditions, consistent with the added effort required to manage greater overall resistance.

In addition to these findings, we observed clear effects of load condition, force magnitude, and reach configuration on peak velocity (PV) and the timepoint of peak velocity (TPV). Both matched load conditions (elastic/elastic, viscous/viscous) reduced PV compared to the no load/no load baseline, with viscous/viscous showing larger reductions. This suggests that resisting either position-dependent elastic loads or velocity-dependent viscous loads constrains movement speed, with viscous resistance having a stronger damping effect. Mismatched load conditions also generally reduced PV, but the extent varied depending on the specific pairing, indicating that the motor system must adjust movement speed differently for each limb depending on its load condition. Reach configuration also mattered: asymmetric and symmetric short reaches both showed lower PV than symmetric long reaches, reflecting the shorter distances and reduced demands in those conditions. Force magnitude had a consistent effect as well, with medium force leading to slightly reduced PV across conditions. For TPV, clear differences emerged between matched and mismatched load conditions. Under matched loads, both hands adjusted timing similarly, preserving temporal coupling. In contrast, mismatched loads led to significant timing differences between the limbs, showing that each hand adjusted its peak velocity timing independently to manage its specific load demands. This decoupling of timing under mismatched loads suggests a flexible, limb-specific control strategy that trades strict synchrony for successful task completion. Importantly, the fixed effect of hand showed that the right hand reached peak velocity slightly earlier overall—a result consistent with all participants being right-hand dominant, with the dominant hand typically moving faster (Peters, 1981; Serrien et al., 2006; Wang & Sainburg, 2007; Goble & Brown, 2008; Buckingham & Carey, 2009). Reach configuration and force magnitude also influenced TPV: asymmetric and symmetric short reaches allowed the system to reach peak velocity sooner compared to symmetric long reaches, while higher force magnitude caused small but reliable delays, reflecting the additional time needed to overcome greater resistance. Together, these findings highlight how the motor system balances shared coordination goals with limb-specific adaptations, maintaining overall task performance even under mismatched load conditions.

We also observed clear effects of load condition, force magnitude, and reach configuration on peak acceleration (PA) and the timepoint of peak acceleration (TPA). Both matched load conditions (elastic/elastic, viscous/viscous) reduced PA compared to no load/no load, with viscous/viscous showing larger reductions. This indicates that resisting velocity-dependent viscous loads places a greater constraint on rapid movement changes than position-dependent elastic loads. Mismatched load conditions also generally reduced PA, but the effects varied depending on the specific pairing and which hand carried the load. Notably, applying a load to the dominant (right) hand tended to reduce PA more than loading the non-dominant hand, consistent with the idea that the dominant limb takes on a larger role in driving movement dynamics (Wang & Sainburg, 2007). These results suggest that the motor system flexibly tunes acceleration for each limb to adapt to specific load demands, rather than applying a uniform command to both hands. For TPA, we observed a strong difference between matched and mismatched load conditions. Matched loads preserved temporal synchrony, with both hands reaching peak acceleration at similar times. In contrast, mismatched loads—especially those involving viscous resistance—produced significant timing differences between the limbs. This decoupling of timing highlights the system’s ability to prioritize limb-specific adaptations when coordination demands increase, trading off continuous temporal coupling to ensure each hand can handle its unique load. Additionally, reach configuration affected both PA and TPA: symmetric short and asymmetric reach configurations consistently reduced peak acceleration and allowed the system to reach it sooner compared to symmetric long reaches, reflecting reduced movement demands. Force magnitude also had consistent effects, with medium force reducing PA and slightly delaying TPA, indicating that managing greater resistance requires refined, slower control. Together, these findings extend prior evidence of strong temporal coupling in symmetric tasks (Kelso et al., 1979) by showing that while synchrony is preserved under matched load conditions, it can break down under mismatched load conditions, particularly when viscous resistance is involved. This demonstrates the motor system’s adaptive strategy: initiating movements in synchrony, allowing independent limb-specific adjustments during execution, and working to maintain overall task goals even at the cost of strict temporal synchrony.

Lastly, we found that load condition, force magnitude, and reach configuration all shaped peak deceleration (PD) and the timepoint of peak deceleration (TPD). Both matched load conditions (elastic/elastic, viscous/viscous) reduced PD relative to no load/no load, with viscous/viscous producing larger reductions. This suggests that velocity-dependent viscous loads place greater demands on slowing movements than position-dependent elastic loads. Mismatched load conditions further amplified these effects, with the degree of PD reduction varying by the specific pairing and which limb carried the load. Notably, the dominant (right) hand showed greater sensitivity to viscous resistance, highlighting asymmetries in how each limb adjusts deceleration. These results suggest that under mismatched load conditions, the motor system cannot rely on a uniform deceleration strategy but must adapt each limb’s control to its specific load demands. For TPD, matched load conditions preserved temporal synchrony, with both hands reaching peak deceleration at similar times. In contrast, mismatched load conditions disrupted this synchrony, producing significant timing differences between the limbs. Viscous resistance, especially in mismatched pairings, posed a particular challenge by inducing larger temporal shifts. Reach configuration also influenced PD and TPD: symmetric short and asymmetric reaches consistently reduced both peak deceleration and its timing compared to symmetric long reaches, reflecting the shorter distances and reduced demands. Force magnitude also played a role, with medium force reducing PD and slightly delaying TPD, indicating the need for more gradual, controlled slowing when managing greater resistance. Overall, these findings highlight the motor system’s flexible coordination strategy: while matched load conditions allow for synchronized deceleration, mismatched loads force the system to prioritize limb-specific adaptations over strict bilateral coupling, especially during the deceleration phase.

Looking across all dependent measures, several clear patterns emerged. Matched load conditions— especially elastic/elastic—generally preserved movement profiles similar to the no load/no load baseline, suggesting that position-dependent elastic loads can be more easily integrated into planning when applied symmetrically. In contrast, viscous loads, whether matched or mismatched, consistently reduced peak velocity, acceleration, and deceleration, reflecting their velocity-dependent resistance that directly dampens movement speed and intensity. Timing measures—including time to peak velocity, acceleration, and deceleration—also highlighted differences between matched and mismatched load conditions. Matched loads maintained temporal synchrony across limbs, while mismatched loads often disrupted it, requiring each hand to adjust timing independently. Notably, some elastic mismatched load conditions preserved synchrony better than viscous mismatches, likely because elastic resistance is steadier and more predictable, supporting coordinated planning. For movement end time (ME), elastic/elastic loads produced greater delays than viscous/viscous loads, suggesting that position-dependent resistance affects final movement phases more strongly. Interestingly, some mismatched conditions (e.g., no load/viscous) still showed preserved synchrony at movement end, implying that even when earlier phases become decoupled, the motor system can prioritize endpoint coordination. This temporal decoupling reflects a strategic trade-off by the CNS, prioritizing efficient adaptation to limb-specific constraints over strict temporal symmetry between the limbs. It allows each limb to optimize its trajectory to overcome its specific load condition, even at the cost of synchrony during the movement itself. The observed behavioral pattern suggests the nervous system implements a sequence of coordinated processes that integrates shared motor planning with independent, limb-specific execution:

1. A unified command initiates bilateral movement, aligning with task demands for simultaneous action.
2. Each limb adapts its movement to its specific load, resulting in distinct kinematics despite a shared goal.
3. These load-dependent adjustments can cause the limbs to reach peak velocity, acceleration, and deceleration at different times, disrupting temporal synchrony during the movement.
4. Despite these timing differences, participants often employed a ‘hover’ strategy—briefly slowing the faster limb near the target to allow the slower limb, facing increased resistance, to catch up.
5. Ultimately, both hands arrived at their respective targets near-simultaneously, demonstrating a high-level coordination mechanism that prioritizes endpoint synchrony over continuous temporal symmetry.

This emergent pattern suggests that bimanual reaching under varying load conditions does not rely on rigid interlimb coupling throughout the entire reach. Instead, coordination is achieved through flexible, adaptive control mechanisms that allow each limb to adjust independently while maintaining overall task coordination. In summary, the central nervous system adopts an adaptive coordination strategy, initiating movements in parallel, allowing load-specific adjustments, and realigning at the endpoint to achieve task goals—highlighting a system that prioritizes effective task execution over strict temporal synchrony.

Importantly, the synchronized timing of peak velocity, acceleration, and deceleration under matched load conditions—but the clear decoupling under mismatched loads—points to the critical role of interhemispheric coordination in bimanual control (Geffen et al., 1994; Gerloff & Andres, 2002; Serrien & Brown, 2002; Morishita et al., 2022). Prior research shows that the corpus callosum is essential for coordinating timing and force between the hands during symmetric tasks, with studies in callosotomy patients demonstrating that without callosal communication, the hemispheres struggle to synchronize continuous bimanual movements (Kennerley et al., 2002; Gooijers & Swinnen, 2014). Functional imaging studies also reveal that the supplementary motor area (SMA) plays a central role in planning and synchronizing bimanual actions, integrating input from both hemispheres to generate coordinated outputs, particularly under symmetric conditions (Jäncke et al., 2000). Our findings suggest that when loads are matched, these interhemispheric pathways efficiently align timing and force profiles, enabling smooth, synchronous movement. However, when loads are mismatched—especially when one limb encounters viscous or elastic resistance—the shared timing signals may no longer suffice, prompting the system to shift toward more independent, limb-specific control, even at the cost of strict temporal synchrony.

These insights have important implications for rehabilitation, particularly for therapies targeting asymmetric limb control, such as in stroke or neurological injury. Our findings show that elastic and viscous loads differently affect bimanual coordination and timing, suggesting that targeted interventions could help retrain impaired systems. For example, robotic devices could apply controlled viscous resistance to the less-affected limb, forcing the affected limb to better engage, or apply elastic resistance to challenge positional coordination. Rather than focusing solely on strengthening or range of motion, rehabilitation protocols could integrate load-based coordination training, progressively adjusting the type and magnitude of loads to restore synchrony and functional bimanual performance. Prior work supports this approach: Patton & Mussa-Ivaldi, 2004 showed that robot-applied adaptive force fields can guide motor learning; Whitall et al., 2000 showed that repetitive bilateral arm training improves motor function in chronic stroke; and Lo et al., 2010 demonstrated that robot-assisted therapy using controlled environments can significantly enhance long-term upper-limb recovery. Building on these advances, our study provides new insights into how specific load types and whether they are matched or mismatched across limbs affect bimanual timing, coordination, and real-time control. By distinguishing the distinct challenges posed by viscous and elastic loads, our findings can help guide the design of precise rehabilitation protocols that deliberately manipulate the type and distribution of load across limbs to retrain interlimb timing, deceleration, and synchronization, ultimately improving real-world bimanual function.

## Supporting information

SupplementalMaterials_Force_Field_Perturbations_Bimanual_Reaching

## Data Availability

Datasets generated and analyzed during the current study are available from the corresponding author upon reasonable request.

## Conflict of Interest Statement

On behalf of all authors, the corresponding author states that there is no conflict of interest.

## Funding

No funding was received.

## Notes

### Competing Interest Statement

The authors have declared no competing interest.

