## SupplementalMaterials_Force_Field_Perturbations_Bimanual_Reaching for "The Influence of Force Field Perturbations on Symmetric and Asymmetric Bimanual Reaching": SupplementalMaterials_Force_Field_Perturbations_Bimanual_Reaching.pdf

#### Supplementary Figure 1.

##### Viscous (L)/Viscous (R) — Low Force Condition (Elastic = 30 N, Viscous = –15 N)

**(a) Bimanual Symmetric : Long Reach**

**(b) Bimanual Symmetric : Short Reach**

**(c) Bimanual Asymmetric : Left Long, Right Short Reach**

**(d) Bimanual Asymmetric : Left Short, Right Long Reach**

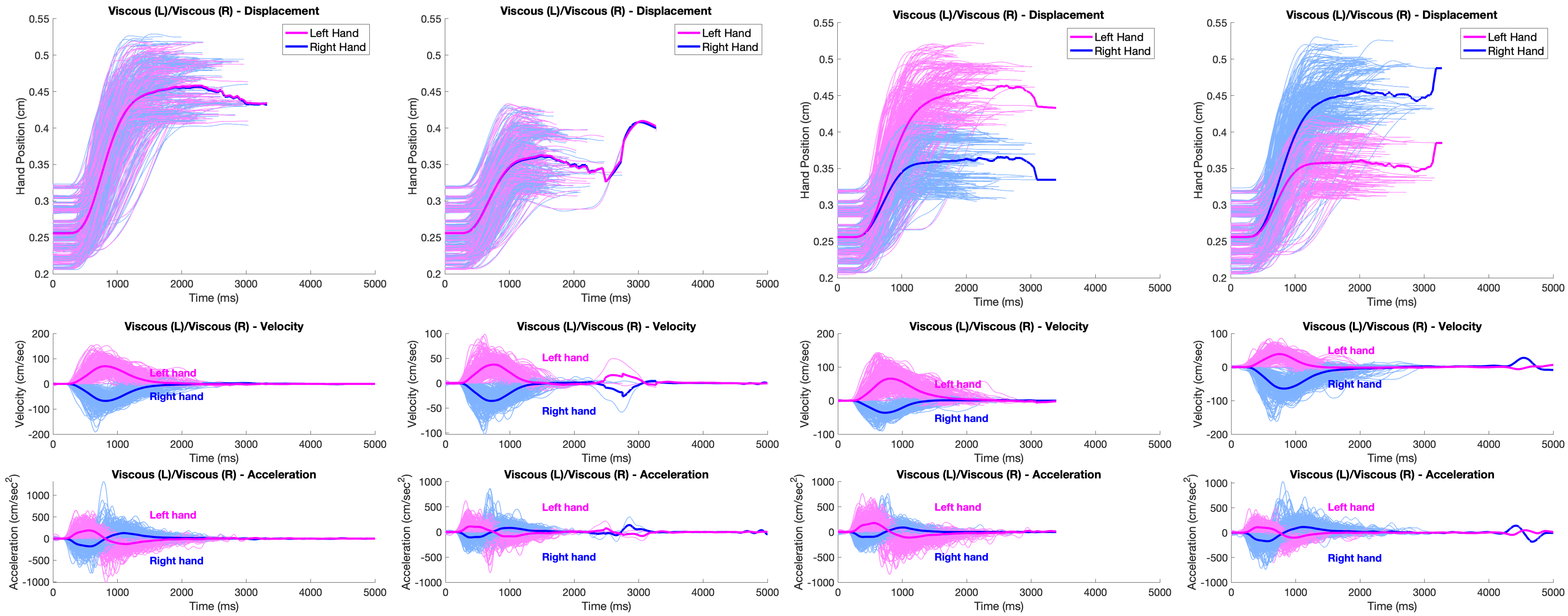


**S1. Kinematic Profile of Low Force Viscous (L)/Viscous (R) Load.** Displacement, velocity, and acceleration over time are shown for the Viscous (L)/Viscous (R) condition under low force, across all four reaching configurations. Individual trajectories from all 30 participants are plotted in light magenta (left hand) and light blue (right hand). Dark magenta and dark blue lines represent the average trajectories for the left and right hands, respectively.

**Supplementary Figure 2.**

#### Elastic (L)/Elastic (R) — Low Force Condition (Elastic = 30 N, Viscous = –15 N)

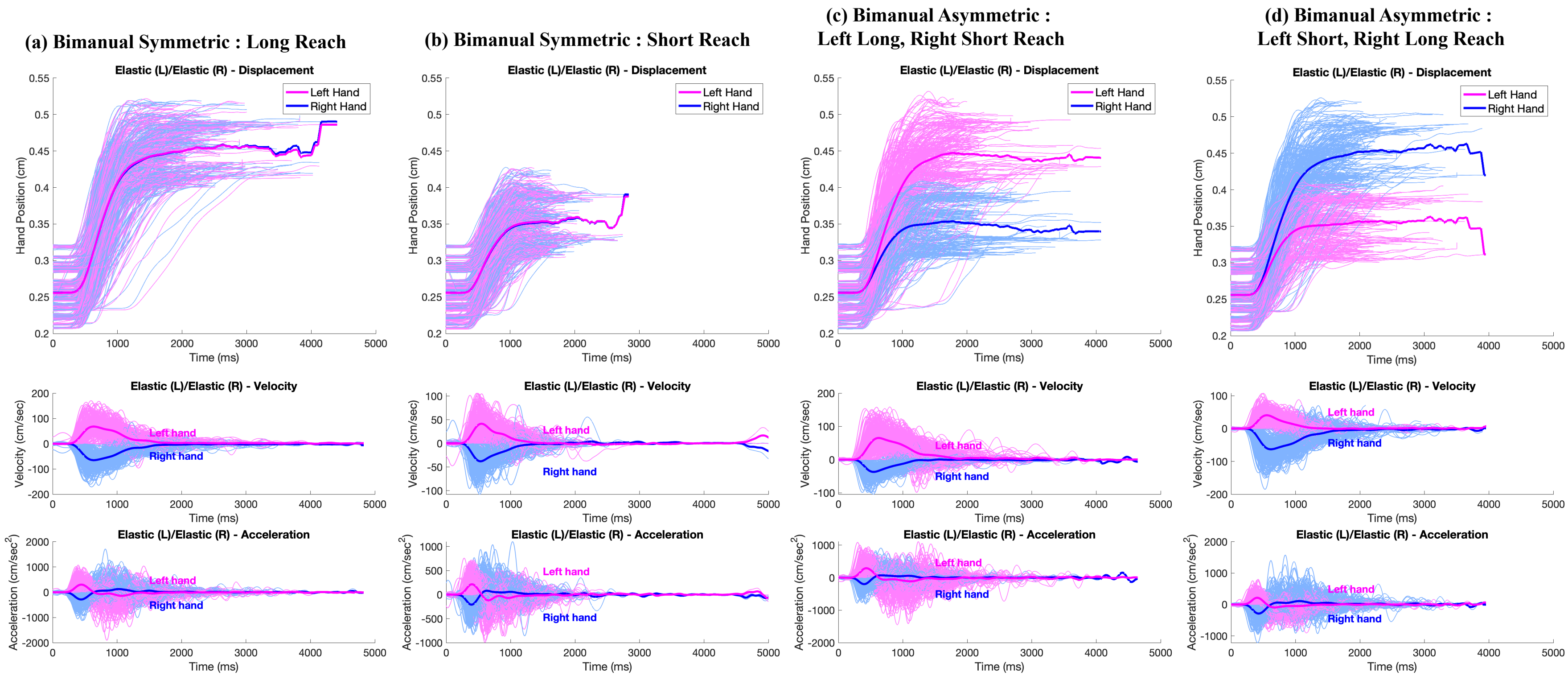


**S2. Kinematic Profile of Low Force Elastic (L)/Elastic (R) Load.** Displacement, velocity, and acceleration over time are shown for the Elastic (L)/Elastic (R) condition under low force, across all four reaching configurations. Individual trajectories from all 30 participants are plotted in light magenta (left hand) and light blue (right hand). Dark magenta and dark blue lines represent the average trajectories for the left and right hands, respectively.

**Supplementary Figure 3.**

#### No Load (L)/Elastic (R) — Low Force Condition (Elastic = 30 N, Viscous = –15 N)

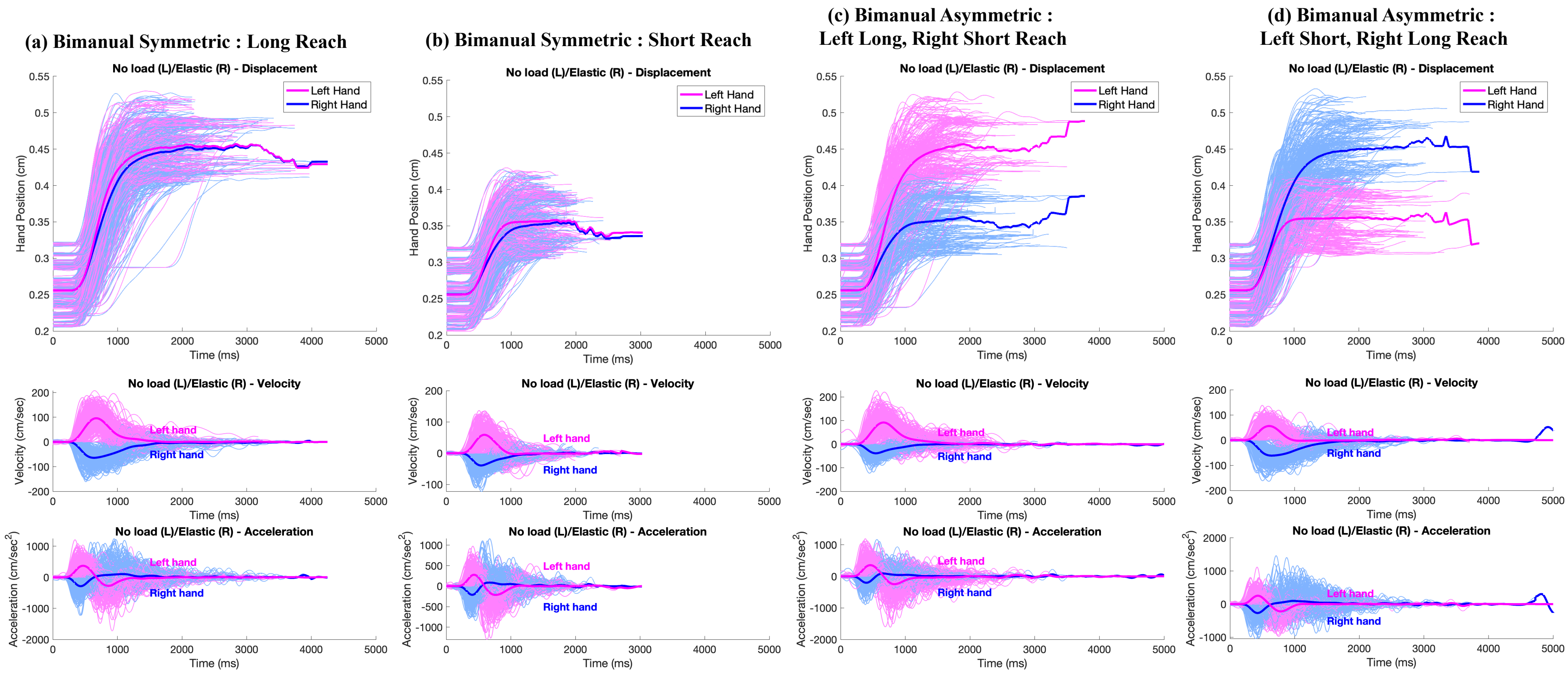


**S3. Kinematic Profile of Low Force No Load (L)/Elastic (R) Load.** Displacement, velocity, and acceleration over time are shown for the No Load (L)/Elastic (R) condition under low force, across all four reaching configurations. Individual trajectories from all 30 participants are plotted in light magenta (left hand) and light blue (right hand). Dark magenta and dark blue lines represent the average trajectories for the left and right hands, respectively.

#### Supplementary Figure 4.

### Elastic (L)/ No Load (R) — Low Force Condition (Elastic = 30 N, Viscous = –15 N)

**(a) Bimanual Symmetric : Long Reach**

**(b) Bimanual Symmetric : Short Reach**

**(c) Bimanual Asymmetric : Left Long, Right Short Reach**

**(d) Bimanual Asymmetric : Left Short, Right Long Reach**

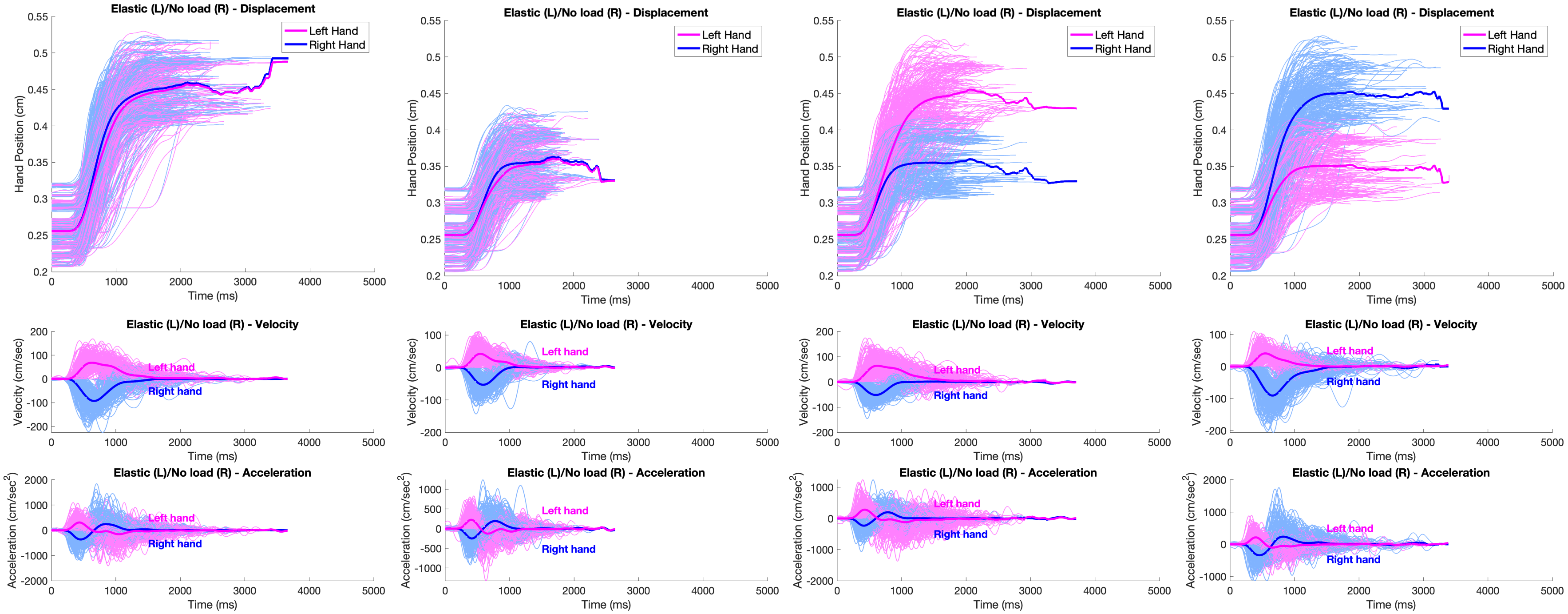


**S4. Kinematic Profile of Low Force Elastic (L)/No Load (R) Load.** Displacement, velocity, and acceleration over time are shown for the Elastic (L)/No Load (R) condition under low force, across all four reaching configurations. Individual trajectories from all 30 participants are plotted in light magenta (left hand) and light blue (right hand). Dark magenta and dark blue lines represent the average trajectories for the left and right hands, respectively.

**Supplementary Figure 5.**

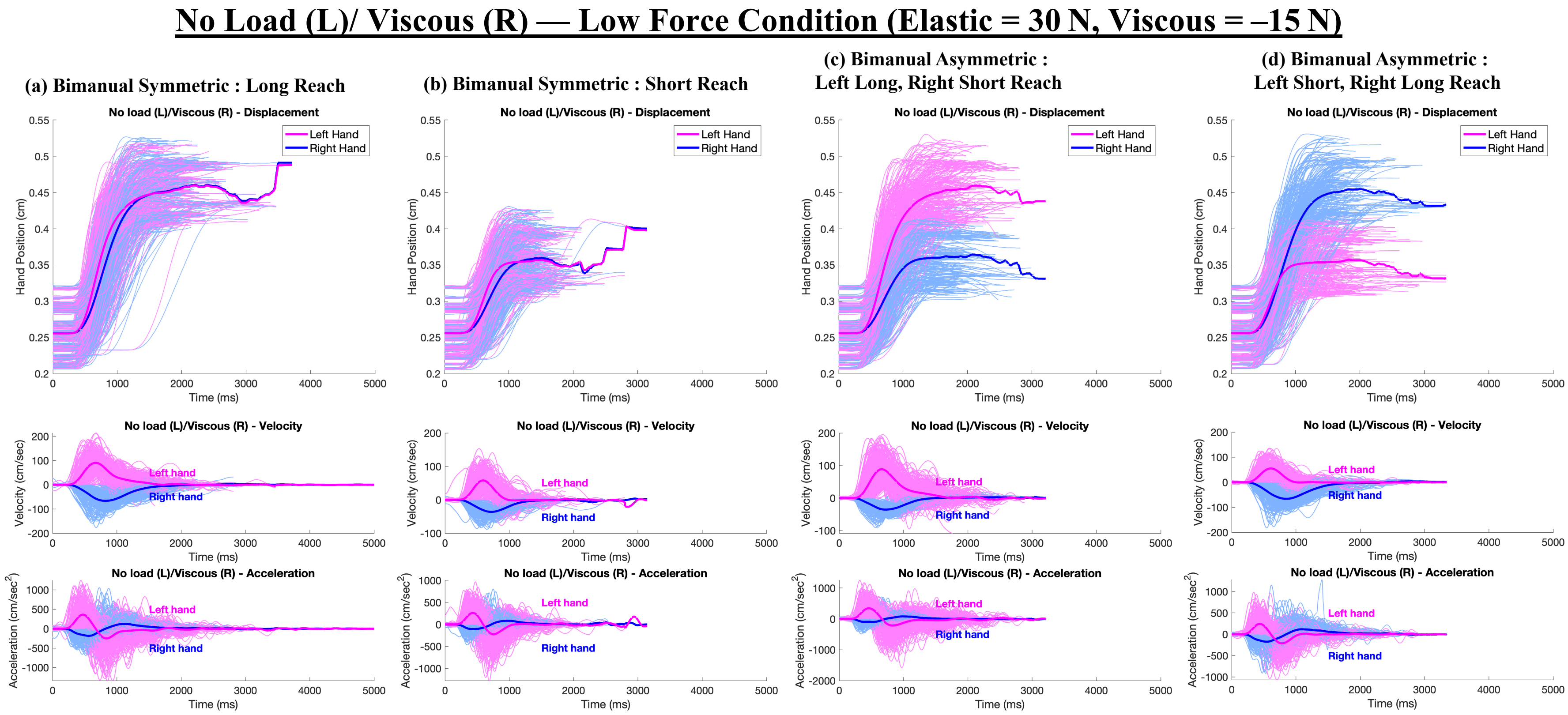


**S5. Kinematic Profile of Low Force No Load (L)/Viscous (R) Load.** Displacement, velocity, and acceleration over time are shown for the No Load (L)/Viscous (R) condition under low force, across all four reaching configurations. Individual trajectories from all 30 participants are plotted in light magenta (left hand) and light blue (right hand). Dark magenta and dark blue lines represent the average trajectories for the left and right hands, respectively.

#### Supplementary Figure 6.

### Viscous (L)/ No Load (R) — Low Force Condition (Elastic = 30 N, Viscous = –15 N)

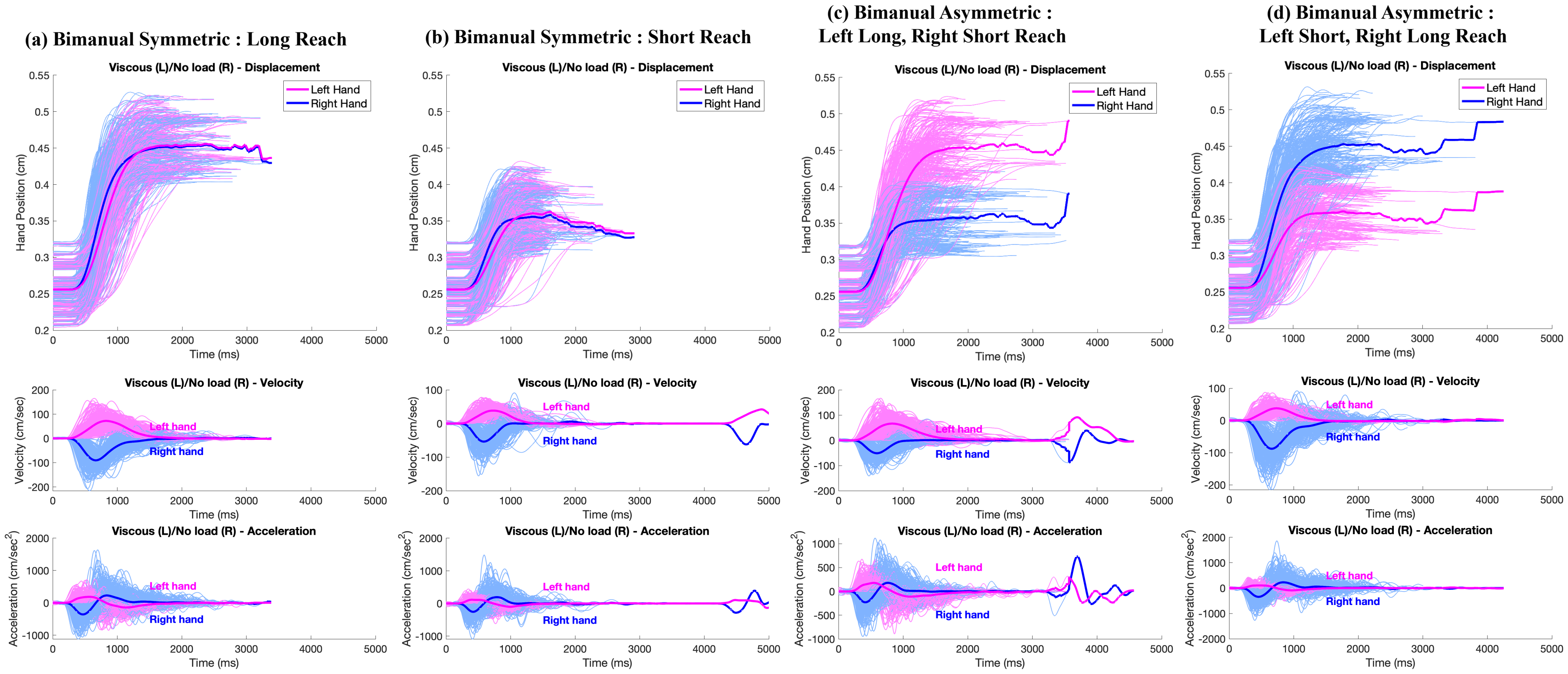


**S6. Kinematic Profile of Low Force Viscous (L)/No Load (R) Load.** Displacement, velocity, and acceleration over time are shown for the Viscous (L)/No Load (R) condition under low force, across all four reaching configurations. Individual trajectories from all 30 participants are plotted in light magenta (left hand) and light blue (right hand). Dark magenta and dark blue lines represent the average trajectories for the left and right hands, respectively.

**Supplementary Figure 7.**

#### Viscous (L)/ Elastic (R) — Low Force Condition (Elastic = 30 N, Viscous = –15 N)

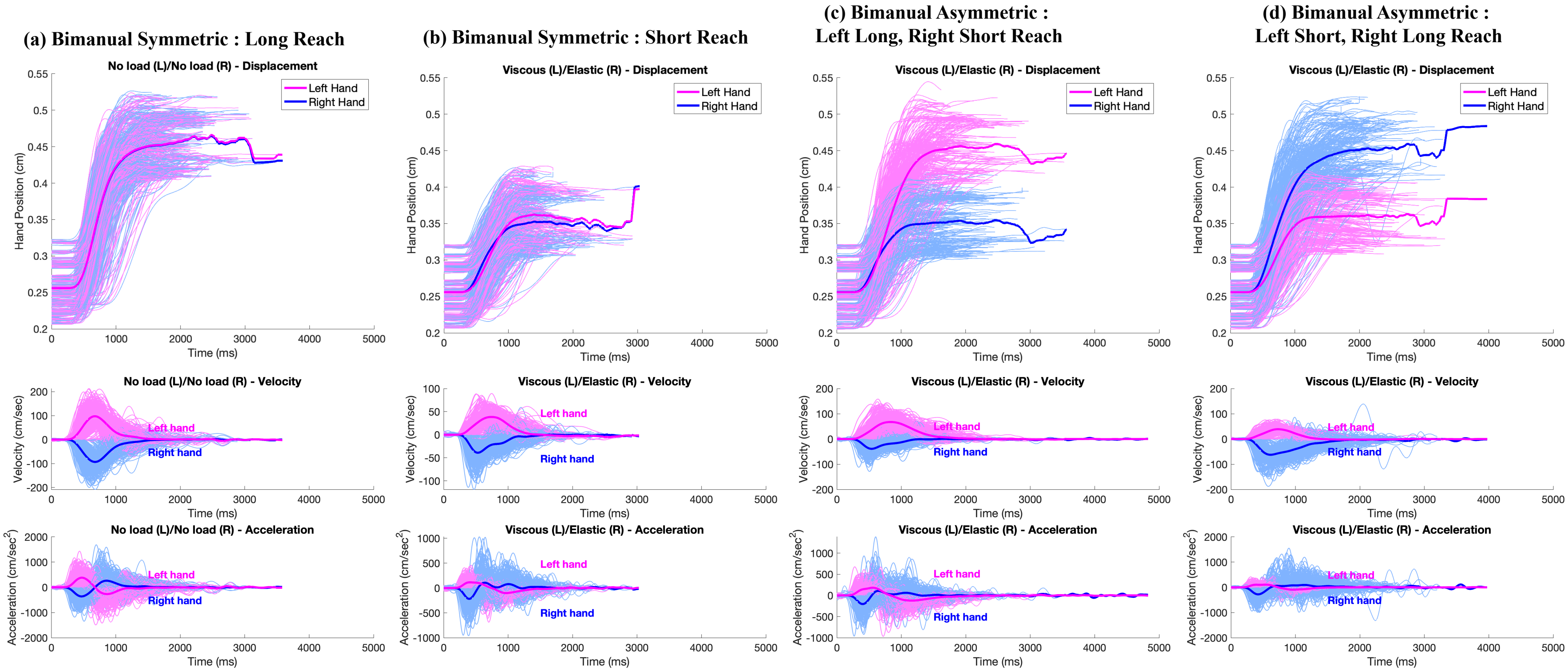


**S7. Kinematic Profile of Low Force Viscous (L)/Elastic (R) Load.** Displacement, velocity, and acceleration over time are shown for the Viscous (L)/Elastic (R) condition under low force, across all four reaching configurations. Individual trajectories from all 30 participants are plotted in light magenta (left hand) and light blue (right hand). Dark magenta and dark blue lines represent the average trajectories for the left and right hands, respectively.

**Supplementary Figure 8.**

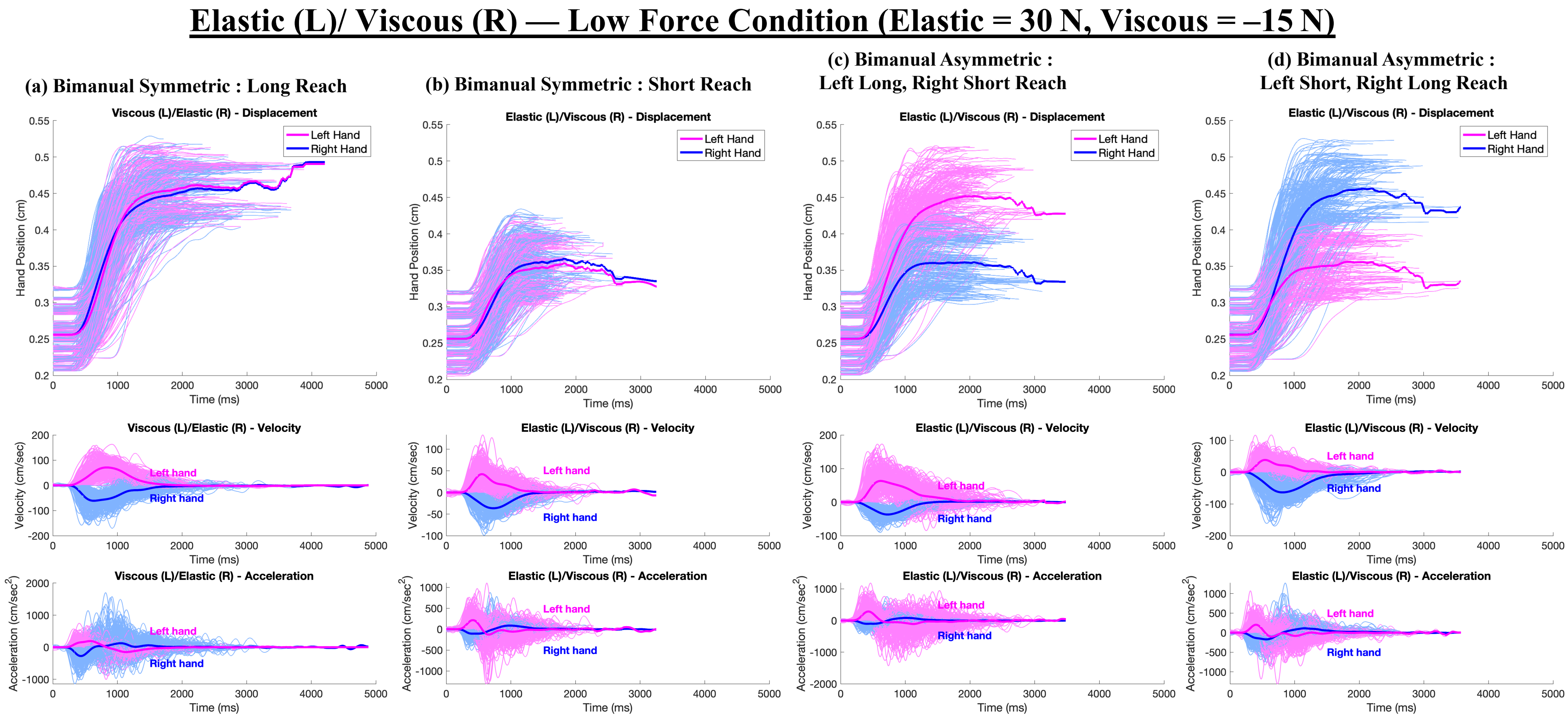


**S8. Kinematic Profile of Low Force Elastic (L)/Viscous (R) Load.** Displacement, velocity, and acceleration over time are shown for the Elastic (L)/Viscous (R) condition under low force, across all four reaching configurations. Individual trajectories from all 30 participants are plotted in light magenta (left hand) and light blue (right hand). Dark magenta and dark blue lines represent the average trajectories for the left and right hands, respectively.

**Supplementary Figure 9.**

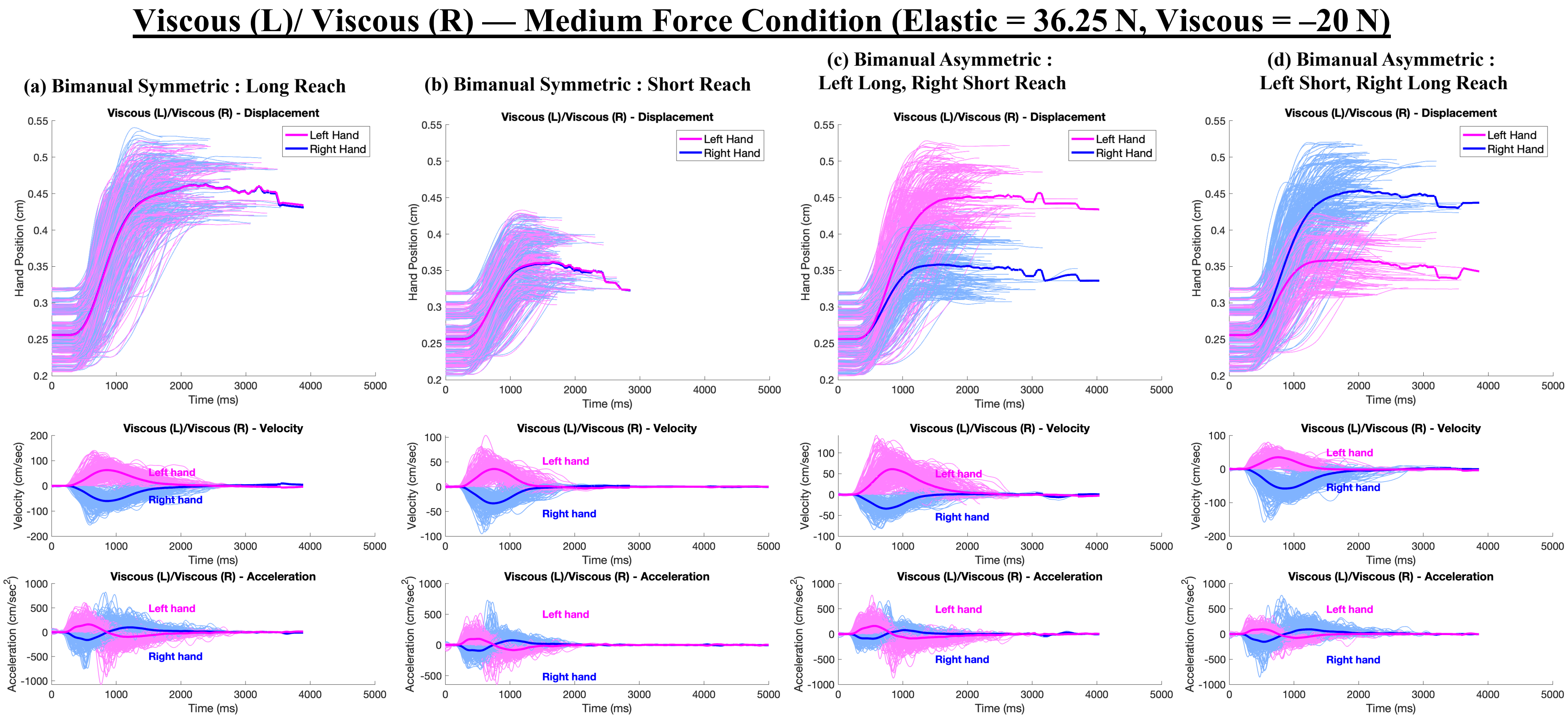


**S9. Kinematic Profile of Medium Force Viscous (L)/Viscous (R) Load.** Displacement, velocity, and acceleration over time are shown for the Viscous (L)/Viscous (R) condition under medium force, across all four reaching configurations. Individual trajectories from all 30 participants are plotted in light magenta (left hand) and light blue (right hand). Dark magenta and dark blue lines represent the average trajectories for the left and right hands, respectively.

#### Supplementary Figure 10.

### Elastic (L)/ Elastic (R) — Medium Force Condition (Elastic = 36.25 N, Viscous = –20 N)

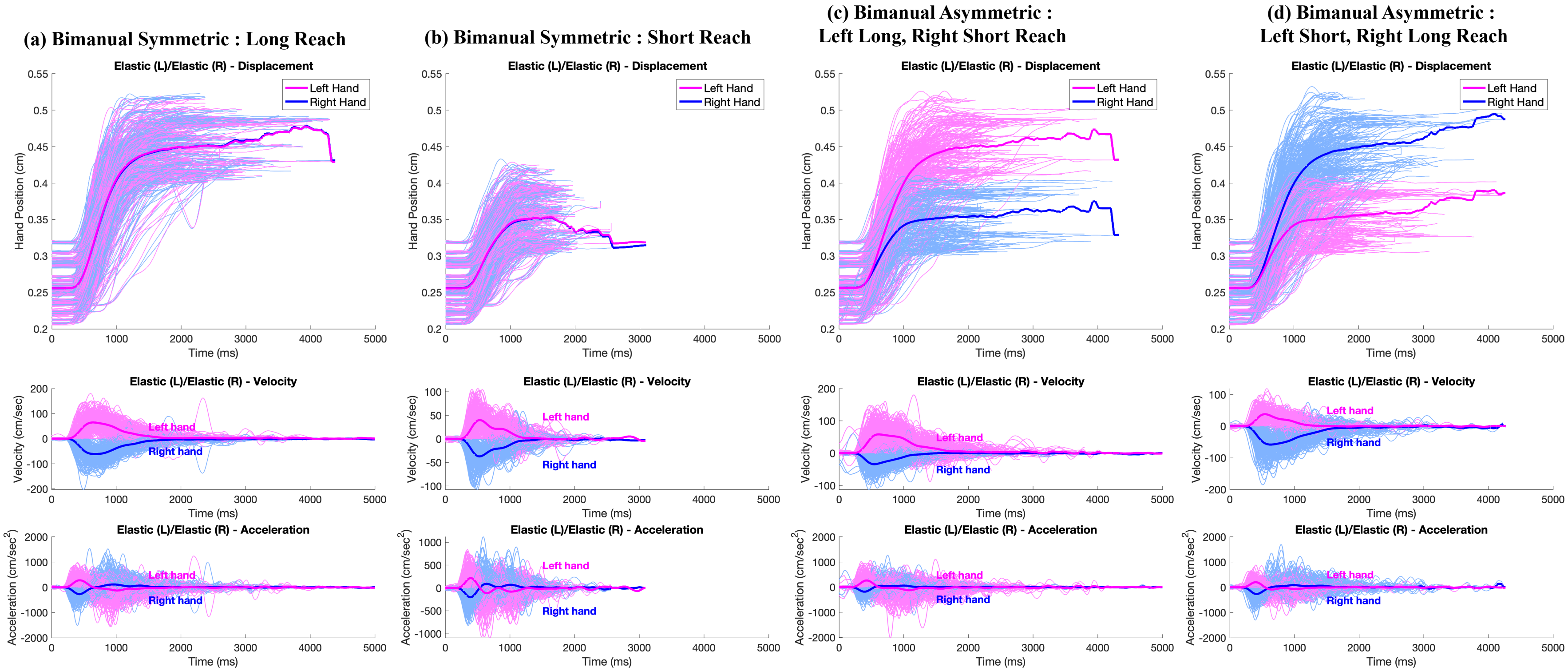


**S10. Kinematic Profile of Medium Force Elastic (L)/Elastic (R) Load.** Displacement, velocity, and acceleration over time are shown for the Elastic (L)/Elastic (R) condition under medium force, across all four reaching configurations. Individual trajectories from all 30 participants are plotted in light magenta (left hand) and light blue (right hand). Dark magenta and dark blue lines represent the average trajectories for the left and right hands, respectively.

#### Supplementary Figure 11.

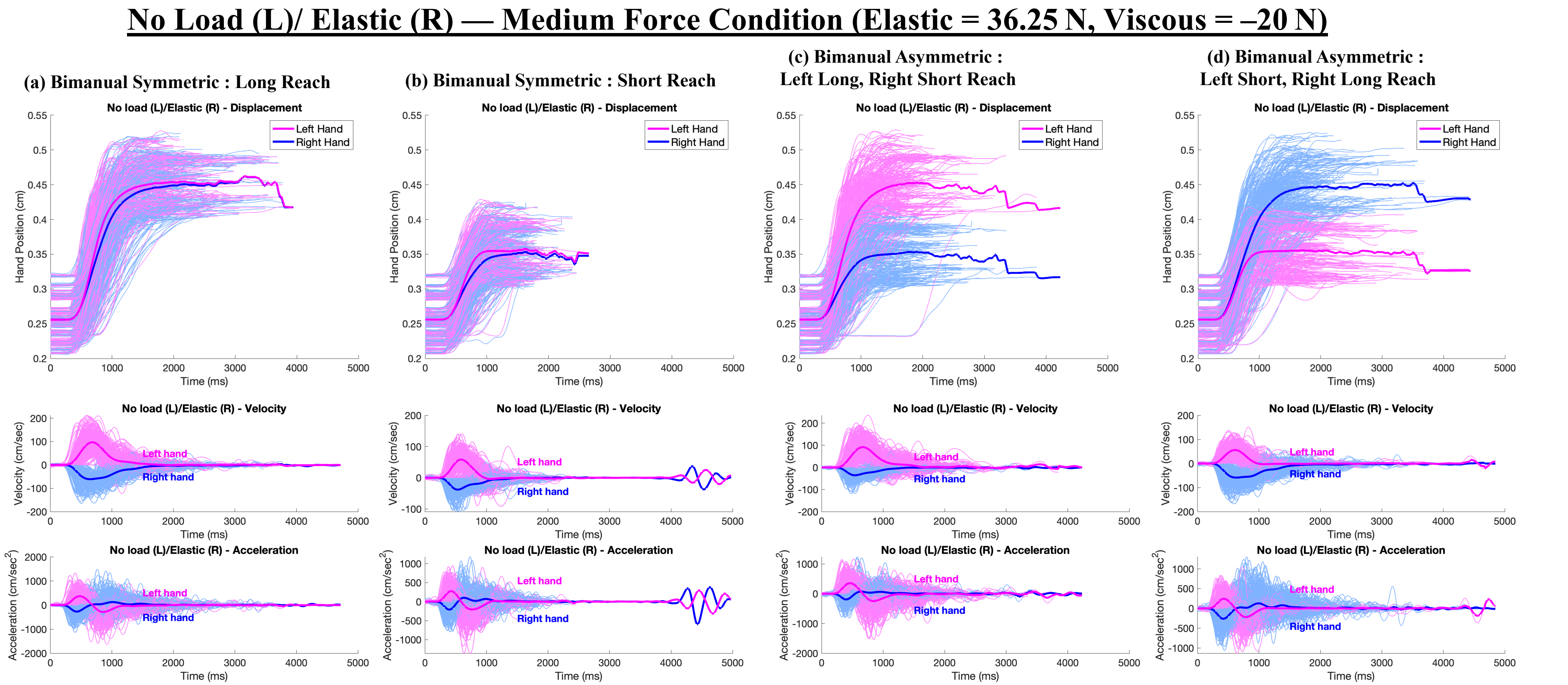


**S11. Kinematic Profile of Medium Force No Load (L)/Elastic (R) Load.** Displacement, velocity, and acceleration over time are shown for the No Load (L)/Elastic (R) condition under medium force, across all four reaching configurations. Individual trajectories from all 30 participants are plotted in light magenta (left hand) and light blue (right hand). Dark magenta and dark blue lines represent the average trajectories for the left and right hands, respectively.

#### Supplementary Figure 12.

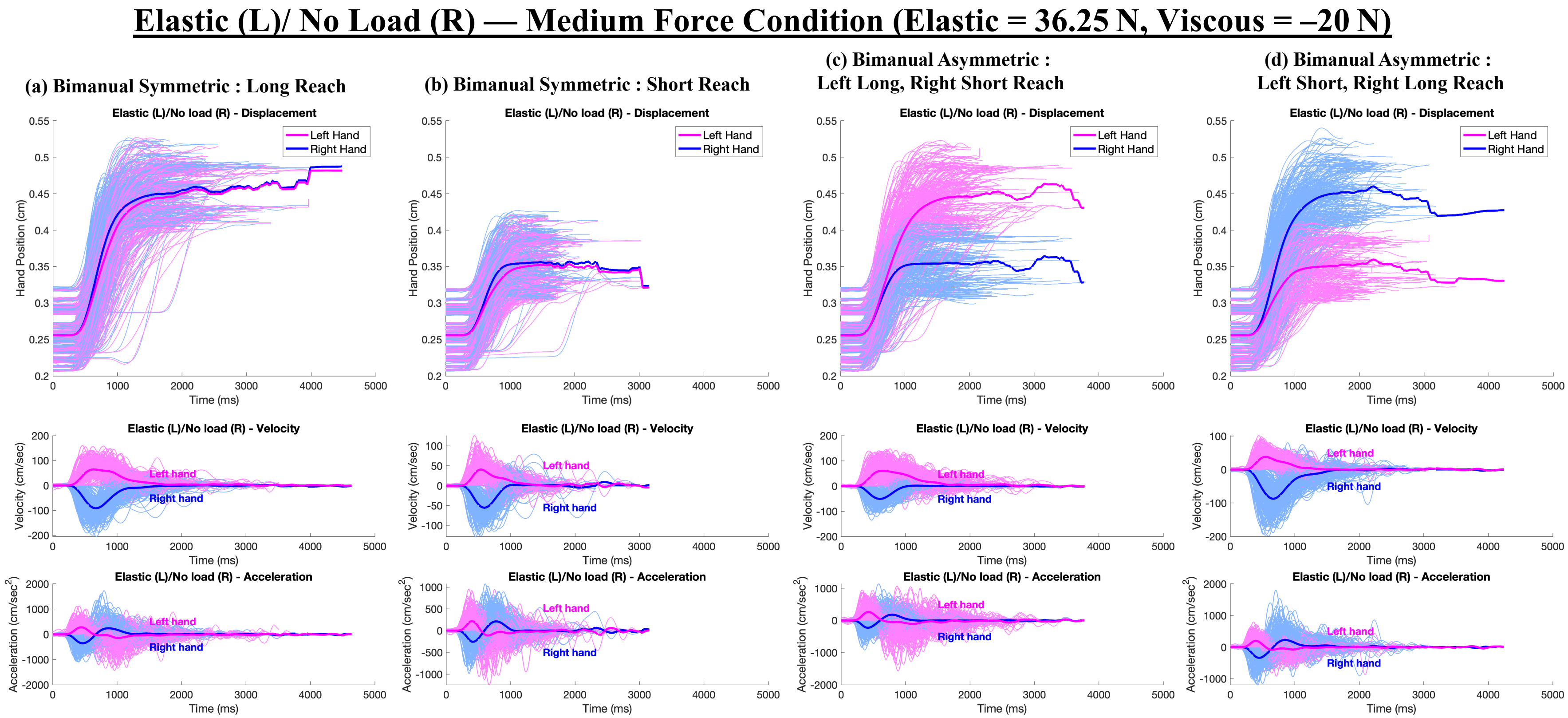


**S12. Kinematic Profile of Medium Force Elastic (L)/No Load (R) Load.** Displacement, velocity, and acceleration over time are shown for the Elastic (L)/No Load (R) condition under medium force, across all four reaching configurations. Individual trajectories from all 30 participants are plotted in light magenta (left hand) and light blue (right hand). Dark magenta and dark blue lines represent the average trajectories for the left and right hands, respectively.

**Supplementary Figure 13.**

#### No Load (L)/ Viscous (R) — Medium Force Condition (Elastic = 36.25 N, Viscous = –20 N)

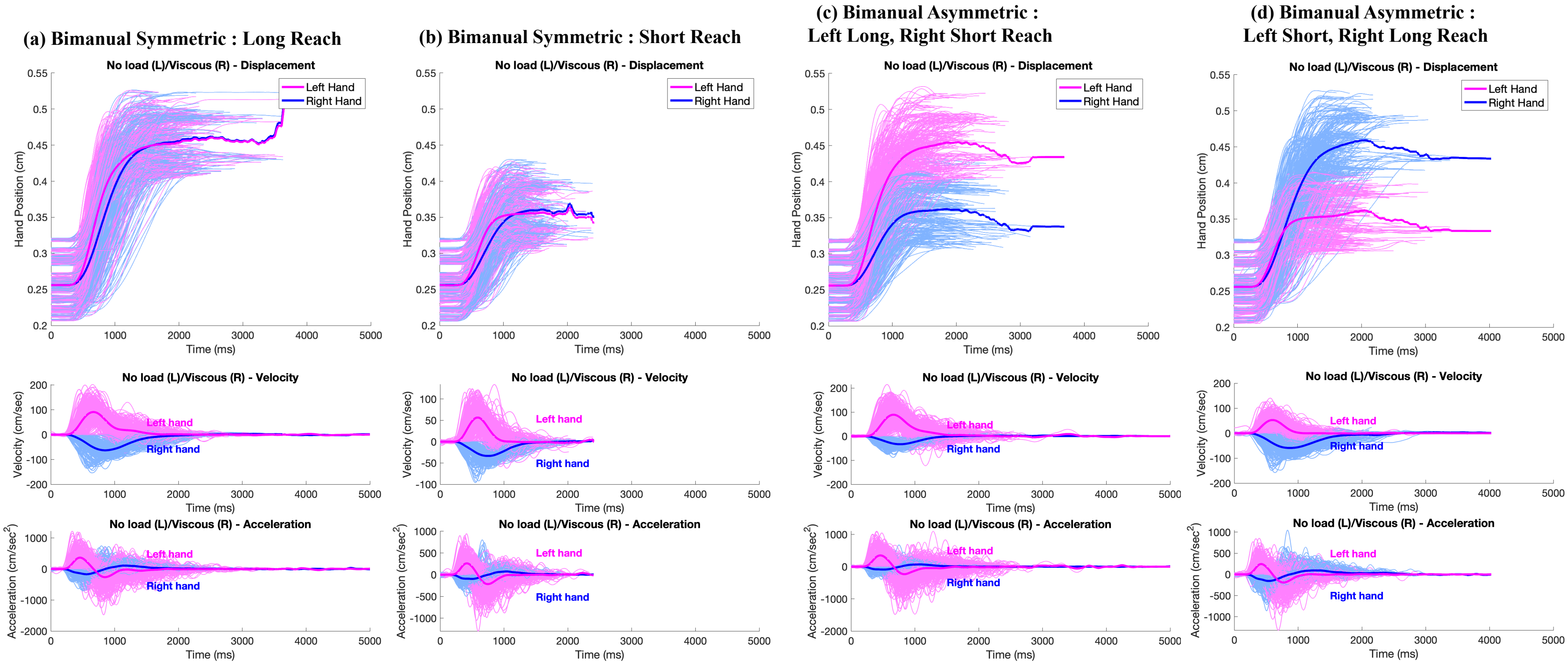


**S13. Kinematic Profile of Medium Force No Load (L)/Viscous (R) Load.** Displacement, velocity, and acceleration over time are shown for the No Load (L)/Viscous (R) condition under medium force, across all four reaching configurations. Individual trajectories from all 30 participants are plotted in light magenta (left hand) and light blue (right hand). Dark magenta and dark blue lines represent the average trajectories for the left and right hands, respectively.

#### Supplementary Figure 14.

### Viscous (L)/ No Load (R) — Medium Force Condition (Elastic = 36.25 N, Viscous = –20 N)

(a) Bimanual Symmetric : Long Reach

(b) Bimanual Symmetric : Short Reach

(c) Bimanual Asymmetric : Left Long, Right Short Reach

(d) Bimanual Asymmetric : Left Short, Right Long Reach

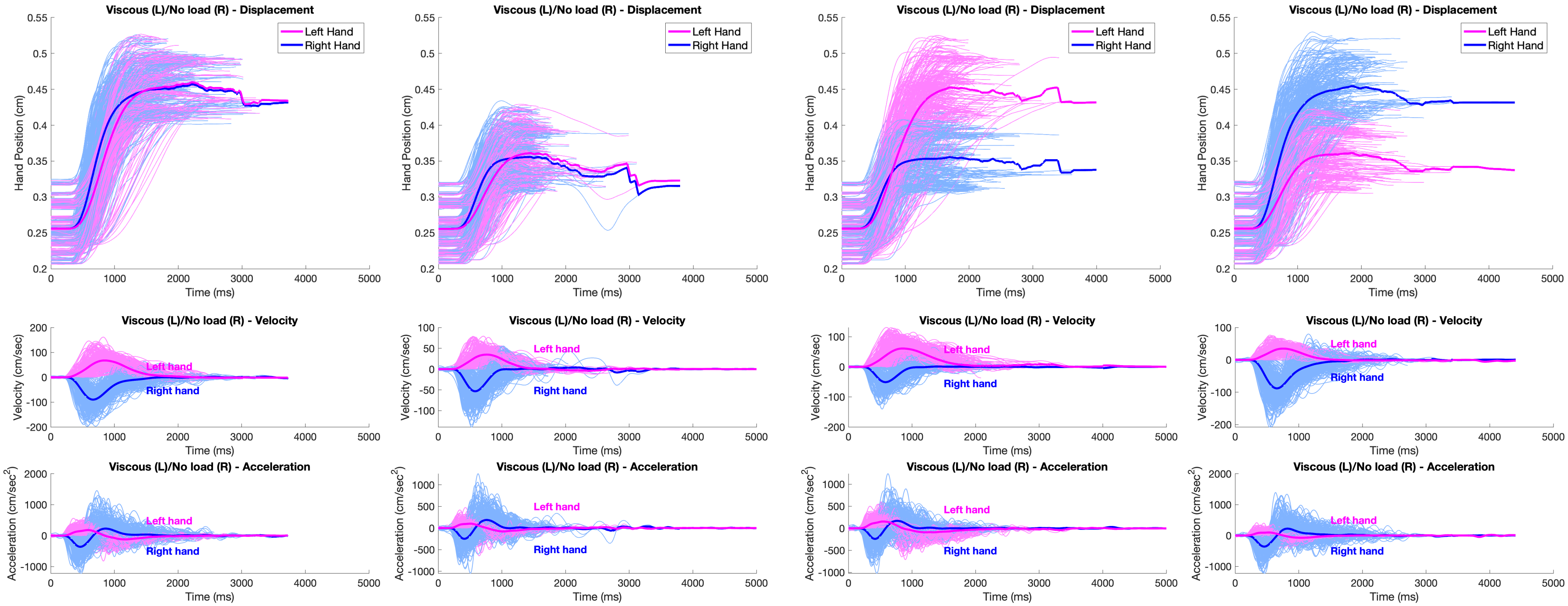


**S14. Kinematic Profile of Medium Force Viscous (L)/No Load (R) Load.** Displacement, velocity, and acceleration over time are shown for the Viscous (L)/No Load (R) condition under medium force, across all four reaching configurations. Individual trajectories from all 30 participants are plotted in light magenta (left hand) and light blue (right hand). Dark magenta and dark blue lines represent the average trajectories for the left and right hands, respectively.

#### Supplementary Figure 15.

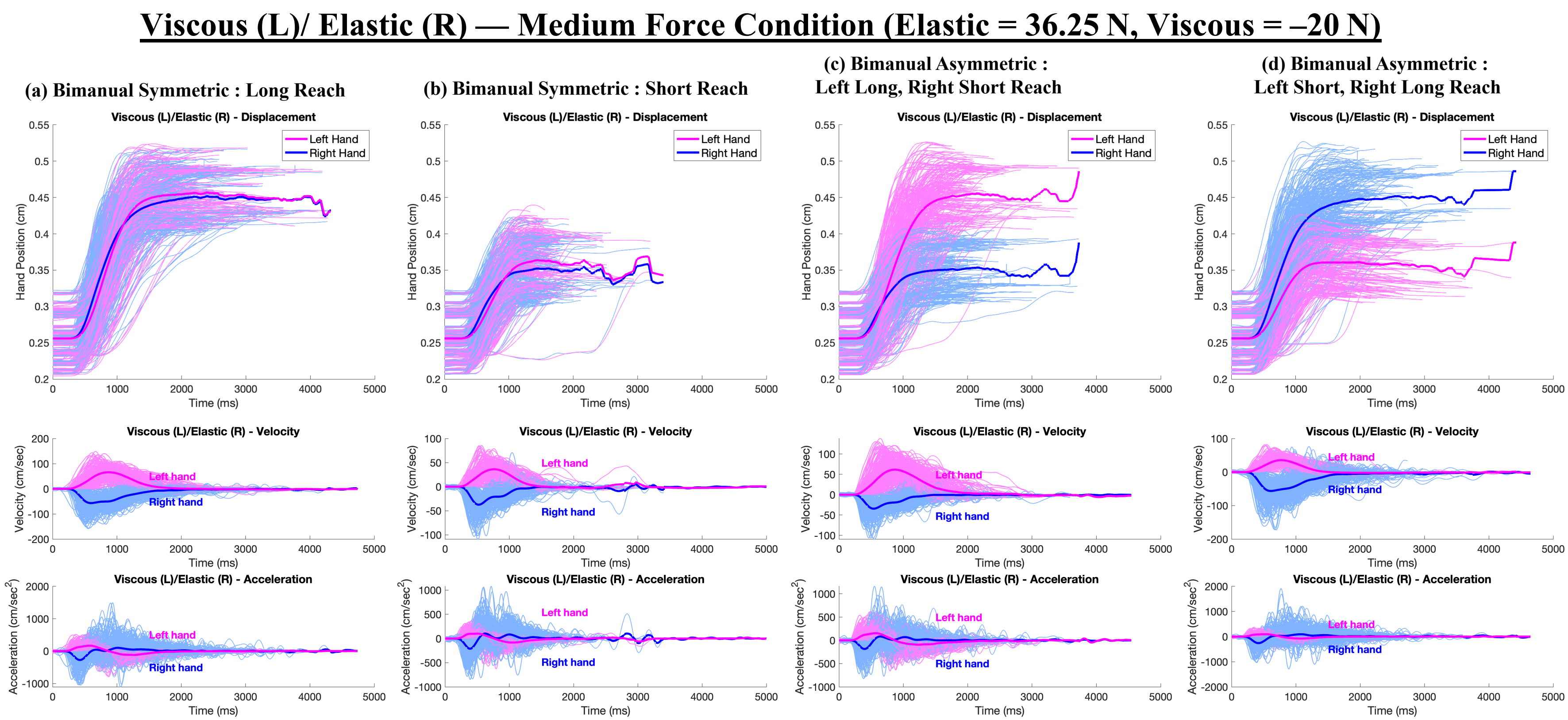


**S15. Kinematic Profile of Medium Force Viscous (L)/Elastic (R) Load.** Displacement, velocity, and acceleration over time are shown for the Viscous (L)/Elastic (R) condition under medium force, across all four reaching configurations. Individual trajectories from all 30 participants are plotted in light magenta (left hand) and light blue (right hand). Dark magenta and dark blue lines represent the average trajectories for the left and right hands, respectively.

#### Supplementary Figure 16.

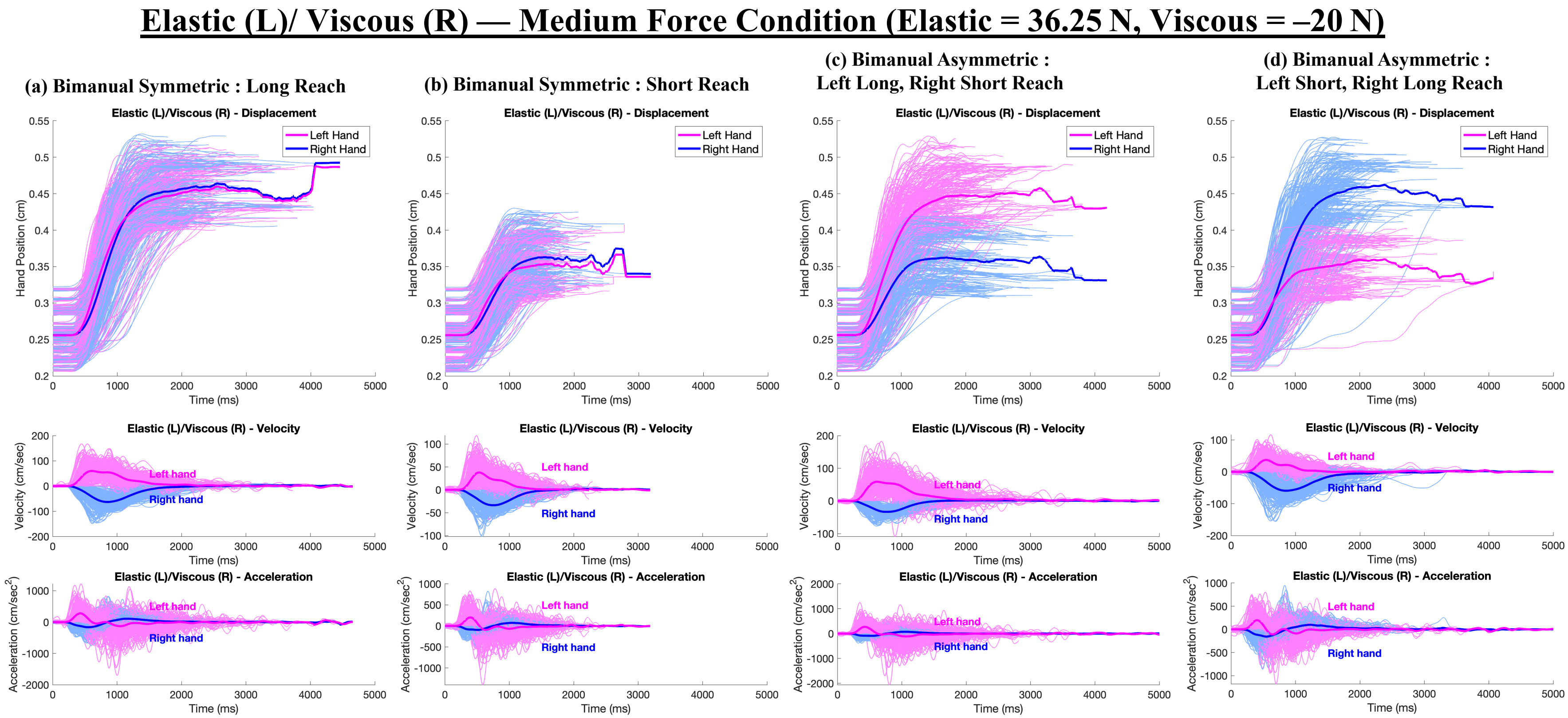


**S16. Kinematic Profile of Medium Force Elastic (L)/Viscous (R) Load.** Displacement, velocity, and acceleration over time are shown for the Elastic (L)/Viscous (R) condition under medium force, across all four reaching configurations. Individual trajectories from all 30 participants are plotted in light magenta (left hand) and light blue (right hand). Dark magenta and dark blue lines represent the average trajectories for the left and right hands, respectively.

### Supplementary Figure 17. Reaction Time (RT) During Reaching Movement.

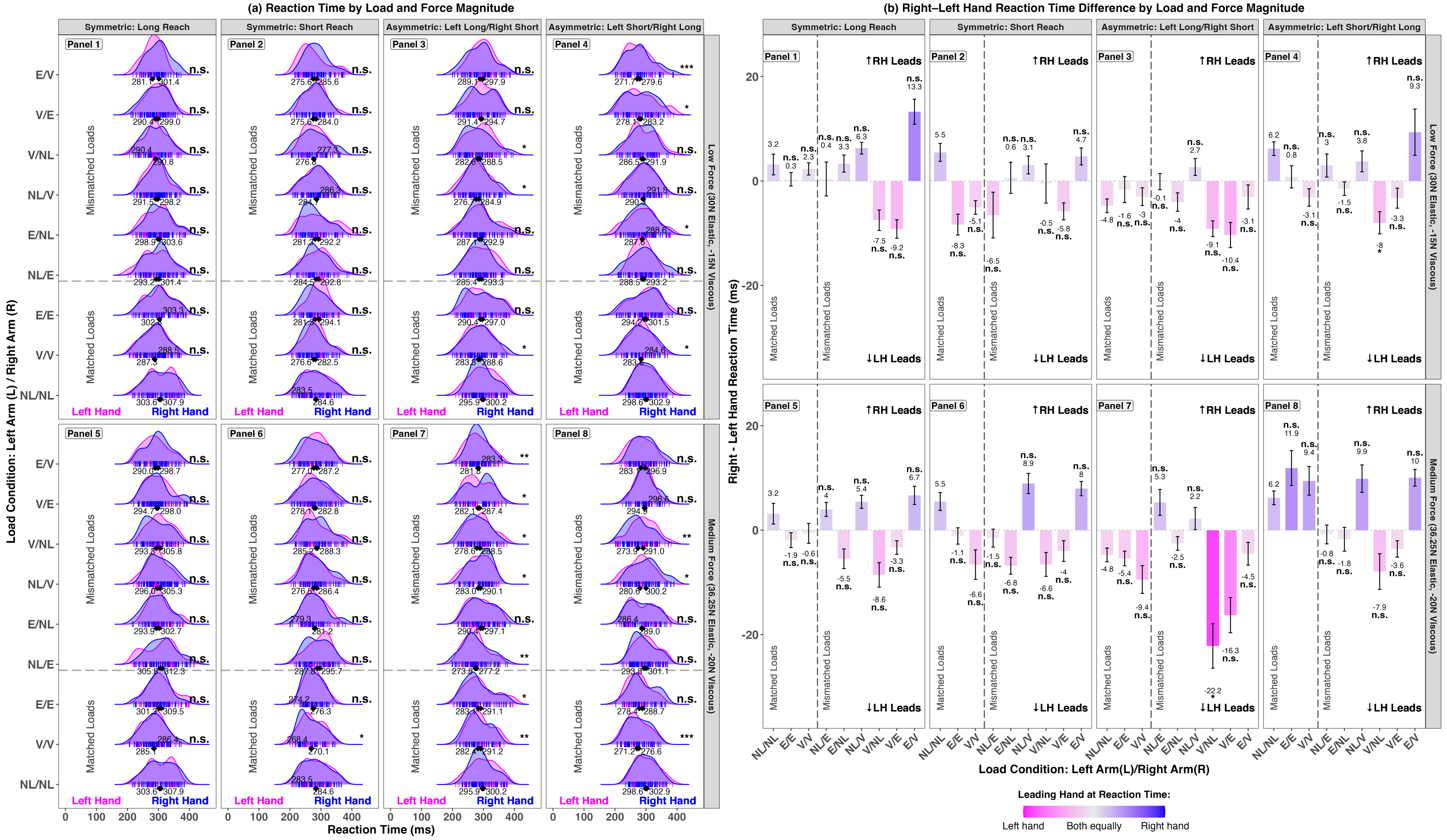


**S17. Reaction Time (RT) During Reaching Movements.** (a) RT across load conditions and reach types. Panels 1-4 show symmetric long, symmetric short, asymmetric left-long/right-short, and asymmetric left-short/right-long reaches under low force (30N Elastic, -15N Viscous); Panels 5-8 show the same under medium force (36.25N Elastic, -20N Viscous). The gray dashed line separates matched from mismatched loads. Asterisks indicate significance relative to the no-load (NL/NL) baseline. Ridge plots depict RT distributions, with vertical bars for individual trials and diamonds for mean RTs. (b) Right-left hand RT differences (layout as in a). Positive values reflect right-hand leads; negative values reflect left-hand leads. Significance is shown relative to NL/NL. The color gradient reflects timing asymmetry: pink = left-hand lead, blue = right-hand lead, gray = synchronized.

#### Supplementary Table 1. Right-Left Hand Difference, Reaction Time (RT)

**T1. Results from Linear Mixed-Effects Models Examining Right-Left Hand Differences in reaction time (RT).** Fixed effects estimates (β), standard errors (SE), and p-values are reported for each model, which included random intercepts for participant and trial number. Fixed effects included Load Condition, Reach Configuration, and Force Magnitude. Significant effects ($p < 0.05$) are shown in bold.

| | **Linear Mixed-Effects Model Results: Right–Left Hand Differences in Reaction Time (RT)** | | |
|---|---|---|---|
| **Fixed effects L=left arm/R=right arm)** | ***β*** | **SE** | ***p*($\chi^2$)** |
| Intercept (No Load (L)/No Load (R)) | 3.03 | 2.09 | 0.147 |
| Elastic (L)/Elastic (R) | -2.91 | 1.92 | 0.131 |
| Viscous (L)/Viscous (R) | -3.43 | 1.98 | 0.084 |
| No Load (L)/Elastic (R) | -1.09 | 1.93 | 0.572 |
| Elastic (L)/No Load (R) | -4.05 | 1.90 | **0.033** |
| No Load (L)/Viscous (R) | 3.21 | 1.90 | 0.091 |
| Viscous (L)/No Load (R) | -10.71 | 1.96 | **<0.001** |
| Viscous (L)/Elastic (R) | -8.74 | 2.04 | **<0.001** |
| Elastic (L)/Viscous (R) | 3.92 | 1.94 | **0.043** |
| Reach Configuration [Symmetric Short Reach] | -1.21 | 1.33 | 0.363 |
| Reach Configuration [Asymmetric Left-Long/Right-Short Reach] | -5.63 | 1.29 | **<0.001** |
| Reach Configuration [Asymmetric Left-Short/Right-Long Reach] | 2.01 | 1.30 | 0.122 |
| Force Magnitude [Medium Force: 36.25N Elastic, -20N Viscous] | -0.47 | 0.94 | 0.619 |
| **Random effects** | **Groups** | | **SD** |
| | Trial Number | Intercept | 3.25 |
| | Participant | Intercept | 6.66 |
| | Residual | | 20.68 |
| | Observations: 2017 | | |
| **Full Model:** Right-Left Difference in RT ~ Load Condition + Reach Configuration + Force Magnitude + (1 \| Trial Number) + (1 \| Participant) | | | |

#### Supplementary Table 2. Left/Right Pairwise Comparison, Reaction Time (RT)

**T2. Post-Hoc Comparisons of Reaction Time (RT) between hands.** Tukey-adjusted pairwise comparisons of left- and right-hand RT within each load condition, across all reach types and force levels.

| Reaction Time (RT) | | | | | | | | | | | | | | | | |
|---|---|---|---|---|---|---|---|---|---|---|---|---|---|---|---|---|
| Symmetric Long Reach<br><br>Low Force:<br>30N Elastic, -15N Viscous | | | | | Symmetric Short Reach<br><br>Low Force:<br>30N Elastic, -15N Viscous | | | | Asymmetric Left Long. Right Short Reach<br><br>Low Force:<br>30N Elastic, -15N Viscous | | | | Asymmetric Left Short, Right Long Reach<br><br>Low Force:<br>30N Elastic, -15N Viscous | | | |
| **Loads Applied: Left/Right Arm** | ***β*** | **SE** | **Z-Ratio** | **P Value** | ***β*** | **SE** | **Z-Ratio** | **P Value** | ***β*** | **SE** | **Z-Ratio** | **P Value** | **Estimate** | **SE** | **Z-Ratio** | **P Value** |
| NL/NL | -3.73 | 6.94 | -0.54 | 0.591 | 1.07 | 7.37 | 0.15 | 0.884 | 3.37 | 6.73 | 0.50 | 0.617 | 2.67 | 7.45 | 0.36 | 0.720 |
| V/V | -0.51 | 6.21 | -0.08 | 0.935 | 6.09 | 7.39 | 0.82 | 0.410 | -2.12 | 7.28 | -0.29 | 0.771 | 3.54 | 8.05 | 0.44 | 0.660 |
| E/E | -1.38 | 7.12 | -0.19 | 0.846 | 9.04 | 8.10 | 1.12 | 0.264 | 6.00 | 7.14 | 0.84 | 0.401 | 6.75 | 7.39 | 0.91 | 0.361 |
| NL/E | -6.31 | 6.78 | -0.93 | 0.352 | 6.65 | 8.93 | 0.74 | 0.457 | 8.44 | 7.49 | 1.13 | 0.260 | -5.41 | 7.24 | -0.75 | 0.455 |
| E/NL | -6.38 | 6.53 | -0.98 | 0.329 | 5.98 | 7.84 | 0.76 | 0.446 | 4.43 | 7.04 | 0.63 | 0.529 | -0.53 | 7.07 | -0.08 | 0.940 |
| NL/V | -11.47 | 7.03 | -1.63 | 0.103 | -1.77 | 7.14 | -0.25 | 0.804 | -8.04 | 7.00 | -1.15 | 0.251 | -2.96 | 7.23 | -0.41 | 0.682 |
| V/NL | 1.34 | 7.92 | 0.17 | 0.865 | -0.74 | 8.18 | -0.09 | 0.928 | 5.76 | 7.32 | 0.79 | 0.431 | 3.73 | 7.55 | 0.49 | 0.621 |
| V/E | 13.21 | 7.36 | 1.79 | 0.073 | 4.02 | 8.93 | 0.45 | 0.653 | 3.97 | 7.34 | 0.54 | 0.588 | 6.20 | 8.21 | 0.76 | 0.450 |
| E/V | -19.22 | 8.20 | -2.34 | **0.019** | -8.65 | 6.67 | -1.30 | 0.195 | 9.89 | 7.06 | 1.40 | 0.161 | -11.06 | 8.10 | -1.37 | 0.172 |
| **Medium Force:<br>36.25N Elastic, -20N Viscous** | | | | | **Medium Force:<br>36.25N Elastic, -20N Viscous** | | | | **Medium Force:<br>36.25N Elastic, -20N Viscous** | | | | **Medium Force:<br>36.25N Elastic, -20N Viscous** | | | |
| NL/NL | -3.73 | 6.94 | -0.54 | 0.591 | 1.07 | 7.37 | 0.15 | 0.884 | 3.37 | 6.73 | 0.50 | 0.617 | 2.67 | 7.45 | 0.36 | 0.720 |
| V/V | 0.33 | 6.75 | 0.05 | 0.961 | 4.24 | 8.25 | 0.51 | 0.607 | 3.90 | 8.02 | 0.49 | 0.627 | -7.99 | 8.15 | -0.98 | 0.327 |
| E/E | 4.36 | 6.61 | 0.66 | 0.509 | 3.26 | 7.90 | 0.41 | 0.680 | 6.07 | 7.34 | 0.83 | 0.408 | -12.61 | 8.22 | -1.53 | 0.125 |
| NL/E | -6.68 | 6.75 | -0.99 | 0.322 | 6.34 | 8.09 | 0.78 | 0.433 | -1.13 | 7.84 | -0.14 | 0.886 | 4.96 | 7.02 | 0.71 | 0.480 |
| E/NL | 9.59 | 7.20 | 1.33 | 0.183 | 1.32 | 7.89 | 0.17 | 0.867 | -6.63 | 6.78 | -0.98 | 0.329 | 2.61 | 7.41 | 0.35 | 0.725 |
| NL/V | -10.26 | 6.39 | -1.60 | 0.109 | -13.53 | 8.03 | -1.69 | 0.092 | -5.11 | 8.21 | -0.62 | 0.534 | -22.74 | 7.30 | -3.12 | **0.002** |
| V/NL | 14.03 | 7.09 | 1.98 | **0.048** | 4.39 | 7.41 | 0.59 | 0.553 | 10.48 | 7.34 | 1.43 | 0.154 | 15.71 | 7.32 | 2.15 | **0.032** |
| V/E | -2.20 | 6.89 | -0.32 | 0.749 | -5.58 | 7.66 | -0.73 | 0.466 | -4.23 | 8.30 | -0.51 | 0.610 | -5.79 | 8.34 | -0.69 | 0.488 |
| E/V | -8.81 | 7.22 | -1.22 | 0.223 | 5.33 | 7.38 | 0.72 | 0.470 | 0.19 | 7.34 | 0.03 | 0.979 | -10.37 | 7.33 | -1.41 | 0.157 |

#### Supplementary Table 3. Between-Hand Reaction Time (RT) Difference Across Loads (Relative to NL/NL)

**T3. Post-Hoc Comparisons of Reaction Time (RT) Differences between Hands by Load Condition and Reach Type.** Pairwise contrasts were conducted to assess whether the difference in RT between the right and left hands varied across load conditions, using the No Load/No Load (NL/NL) condition as the reference. As is standard in treatment-versus-control comparisons, NL/NL serves as the baseline and is not shown in the table; all other estimates reflect differences relative to this reference condition.

| Reaction Time (RT) | | | | | | | | | | | | | | | | |
|---|---|---|---|---|---|---|---|---|---|---|---|---|---|---|---|---|
| **Symmetric Long Reach<br><br>Low Force:<br>30N Elastic, -15N Viscous** | | | | | **Symmetric Short Reach<br><br>Low Force:<br>30N Elastic, -15N Viscous** | | | | **Asymmetric Left Long. Right Short Reach<br><br>Low Force:<br>30N Elastic, -15N Viscous** | | | | **Asymmetric Left Short, Right Long Reach<br><br>Low Force:<br>30N Elastic, -15N Viscous** | | | |
| **Loads Applied: Left/Right Arm** | ***β*** | **SE** | **Z-Ratio** | **P Value** | ***β*** | **SE** | **Z-Ratio** | **P Value** | ***β*** | **SE** | **Z-Ratio** | **P Value** | **Estimate** | **SE** | **Z-Ratio** | **P Value** |
| V/V | -0.41 | 4.92 | -0.08 | 0.934 | -9.27 | 6.25 | -1.48 | 0.369 | 2.40 | 5.37 | 0.45 | 0.862 | -9.57 | 5.47 | -1.75 | 0.321 |
| E/E | -0.90 | 4.70 | -0.19 | 0.934 | -10.40 | 5.72 | -1.82 | 0.369 | 0.95 | 5.47 | 0.17 | 0.862 | -6.64 | 6.03 | -1.10 | 0.434 |
| NL/E | -1.69 | 4.86 | -0.35 | 0.934 | -9.71 | 6.34 | -1.53 | 0.369 | 3.73 | 5.71 | 0.65 | 0.862 | -2.12 | 5.48 | -0.39 | 0.799 |
| E/NL | 0.46 | 4.90 | 0.09 | 0.934 | -4.03 | 5.72 | -0.70 | 0.642 | -0.97 | 5.33 | -0.18 | 0.862 | -5.77 | 5.21 | -1.11 | 0.434 |
| NL/V | 3.12 | 4.94 | 0.63 | 0.934 | 0.15 | 5.55 | 0.03 | 0.979 | 8.09 | 5.33 | 1.52 | 0.862 | -0.75 | 5.24 | -0.14 | 0.887 |
| V/NL | -9.76 | 5.32 | -1.84 | 0.177 | -5.16 | 5.78 | -0.89 | 0.595 | -4.75 | 5.54 | -0.86 | 0.862 | -15.43 | 5.49 | -2.81 | 0.040 |
| V/E | -11.56 | 5.15 | -2.24 | 0.121 | -8.45 | 6.68 | -1.26 | 0.413 | -4.80 | 5.48 | -0.88 | 0.862 | -9.86 | 6.34 | -1.55 | 0.321 |
| E/V | 12.83 | 5.92 | 2.17 | 0.121 | 1.85 | 5.41 | 0.34 | 0.837 | 2.65 | 5.38 | 0.49 | 0.862 | 4.88 | 6.02 | 0.81 | 0.557 |
| **Medium Force:<br>36.25N Elastic, -20N Viscous** | | | | | **Medium Force:<br>36.25N Elastic, -20N Viscous** | | | | **Medium Force:<br>36.25N Elastic, -20N Viscous** | | | | **Medium Force:<br>36.25N Elastic, -20N Viscous** | | | |
| V/V | -2.75 | 4.84 | -0.57 | 0.651 | -12.77 | 5.76 | -2.22 | 0.208 | -2.84 | 5.33 | -0.53 | 0.950 | 4.03 | 6.03 | 0.67 | 0.576 |
| E/E | -3.94 | 5.12 | -0.77 | 0.589 | -6.09 | 6.80 | -0.90 | 0.576 | -1.04 | 6.05 | -0.17 | 0.980 | 5.40 | 6.22 | 0.87 | 0.576 |
| NL/E | 1.59 | 4.90 | 0.32 | 0.746 | -4.48 | 5.88 | -0.76 | 0.576 | 9.54 | 5.87 | 1.63 | 0.277 | -5.83 | 5.24 | -1.11 | 0.532 |
| E/NL | -6.05 | 5.02 | -1.21 | 0.589 | -10.87 | 6.16 | -1.76 | 0.208 | -0.14 | 5.38 | -0.03 | 0.980 | -8.07 | 5.70 | -1.41 | 0.419 |
| NL/V | 3.69 | 4.72 | 0.78 | 0.589 | 3.14 | 6.45 | 0.49 | 0.626 | 7.58 | 6.24 | 1.21 | 0.450 | 2.47 | 5.38 | 0.46 | 0.646 |
| V/NL | -9.92 | 5.31 | -1.87 | 0.495 | -10.39 | 5.62 | -1.85 | 0.208 | -17.64 | 5.78 | -3.05 | **0.018** | -14.90 | 5.58 | -2.67 | 0.061 |
| V/E | -7.51 | 4.98 | -1.51 | 0.528 | -6.79 | 5.94 | -1.14 | 0.508 | -11.59 | 6.24 | -1.86 | 0.253 | -9.63 | 6.34 | -1.52 | 0.419 |
| E/V | 4.18 | 5.06 | 0.83 | 0.589 | 3.85 | 5.76 | 0.67 | 0.576 | 0.68 | 5.32 | 0.13 | 0.980 | 3.66 | 5.33 | 0.69 | 0.576 |

#### Supplementary Figure 18. Movement Time (MT) During Reaching Movements.

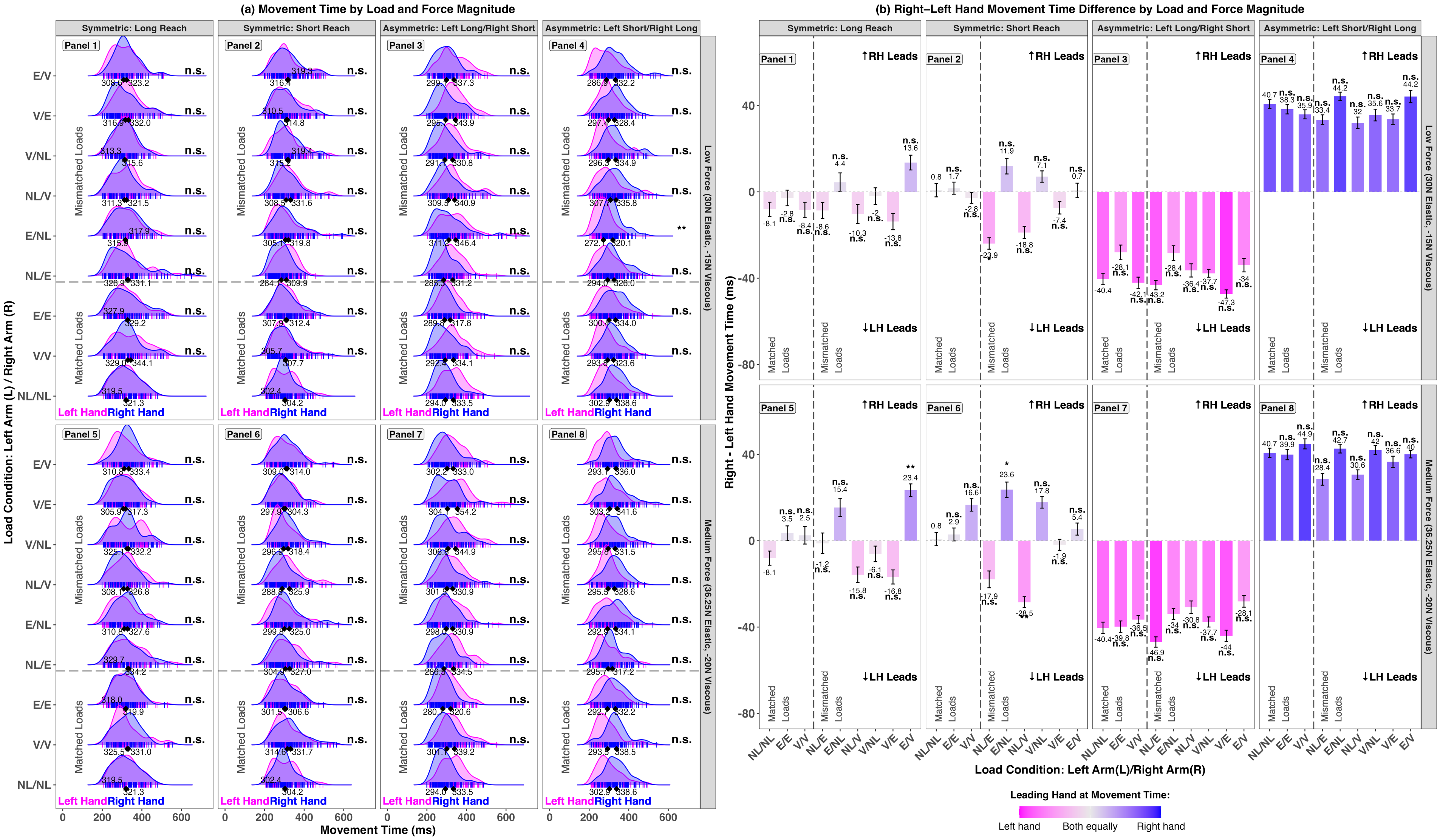


**S18. Movement Time (MT) During Reaching Movements.** (a) MT across load conditions and reach types. Panels 1-4 show symmetric long, symmetric short, asymmetric left-long/right-short, and asymmetric left-short/right-long reaches under low force (30N Elastic, -15N Viscous); Panels 5-8 show the same under medium force (36.25N Elastic, -20N Viscous). The gray dashed line separates matched from mismatched loads. Asterisks indicate significance relative to the no-load (NL/NL) baseline. Ridge plots depict MT distributions, with vertical bars for individual trials and diamonds for mean MTs. (b) Right-left hand MT differences (layout as in a). Positive values reflect right-hand leads; negative values reflect left-hand leads. Significance is shown relative to NL/NL. The color gradient reflects timing asymmetry: pink = left-hand lead, blue = right-hand lead, gray = synchronized.

#### Supplementary Table 4. Right-Left Hand Difference, Movement Time (MT)

**T4. Results from Linear Mixed-Effects Models Examining Right-Left Hand Differences in movement time (MT).** Fixed effects estimates (β), standard errors (SE), and p-values are reported for each model, which included random intercepts for participant and trial number. Fixed effects included Load Condition, Reach Configuration, and Force Magnitude. Significant effects ($p < 0.05$) are shown in bold.

| | **Linear Mixed-Effects Model Results: Right–Left Hand Differences in Movement Time (MT)** | | |
|---|---|---|---|
| **Fixed effects L=left arm/R=right arm)** | ***β*** | **SE** | ***p*($\chi^2$)** |
| Intercept (No Load (L)/No Load (R)) | -3.92 | 3.01 | 0.193 |
| Elastic (L)/Elastic (R) | 3.33 | 2.89 | 0.249 |
| Viscous (L)/Viscous (R) | 2.66 | 2.91 | 0.360 |
| No Load (L)/Elastic (R) | -8.84 | 2.97 | **0.003** |
| Elastic (L)/No Load (R) | 11.40 | 2.95 | **<0.001** |
| No Load (L)/Viscous (R) | -8.70 | 2.96 | **0.003** |
| Viscous (L)/No Load (R) | 4.51 | 2.96 | 0.127 |
| Viscous (L)/Elastic (R) | -5.98 | 2.93 | **0.041** |
| Elastic (L)/Viscous (R) | 9.69 | 2.93 | **0.001** |
| Reach Configuration [Symmetric Short Reach] | 1.53 | 1.96 | 0.434 |
| Reach Configuration [Asymmetric Left-Long/Right-Short Reach] | -35.57 | 1.98 | **<0.001** |
| Reach Configuration [Asymmetric Left-Short/Right-Long Reach] | 40.59 | 1.97 | **<0.001** |
| Force Magnitude [Medium Force: 36.25N Elastic, -20N Viscous] | 2.59 | 1.36 | 0.057 |
| **Random effects** | **Groups** | | **SD** |
| | Trial Number | Intercept | 6.35 |
| | Participant | Intercept | 8.38 |
| | Residual | | 52.07 |
| | Observations: 5969 | | |
| **Full Model:** Right-Left Difference in MT ~ Load Condition + Reach Configuration + Force Magnitude + (1 \| Trial Number) + (1 \| Participant) | | | |

#### Supplementary Table 5. Left/Right Pairwise Comparison, Movement Time (MT)

**T5. Post-Hoc Comparisons of Movement Time (MT) between hands.** Tukey-adjusted pairwise comparisons of left- and right-hand MT within each load condition, across all reach types and force levels.

| Movement Time (MT) | | | | | | | | | | | | | | | | |
|---|---|---|---|---|---|---|---|---|---|---|---|---|---|---|---|---|
| **Symmetric Long Reach**<br><br>**Low Force:**<br>**30N Elastic, -15N Viscous** | | | | | **Symmetric Short Reach**<br><br>**Low Force:**<br>**30N Elastic, -15N Viscous** | | | | **Asymmetric Left Long. Right Short Reach**<br><br>**Low Force:**<br>**30N Elastic, -15N Viscous** | | | | **Asymmetric Left Short, Right Long Reach**<br><br>**Low Force:**<br>**30N Elastic, -15N Viscous** | | | |
| **Loads Applied: Left/Right Arm** | **β** | **SE** | **Z-Ratio** | **P Value** | **β** | **SE** | **Z-Ratio** | **P Value** | **β** | **SE** | **Z-Ratio** | **P Value** | **Estimate** | **SE** | **Z-Ratio** | **P Value** |
| NL/NL | 3.47 | 9.03 | 0.38 | 0.701 | 0.36 | 8.33 | 0.04 | 0.965 | 38.35 | 8.11 | 4.73 | **<0.001** | -37.57 | 8.79 | -4.28 | **<0.001** |
| V/V | 11.71 | 9.13 | 1.28 | 0.200 | 1.69 | 7.79 | 0.22 | 0.828 | 43.42 | 8.11 | 5.35 | **<0.001** | -31.63 | 7.89 | -4.01 | **<0.001** |
| E/E | 2.56 | 8.00 | 0.32 | 0.749 | -2.12 | 7.77 | -0.27 | 0.785 | 28.05 | 8.56 | 3.28 | 0.001 | -33.99 | 7.69 | -4.42 | **<0.001** |
| NL/E | 5.88 | 8.82 | 0.67 | 0.504 | 26.98 | 7.85 | 3.44 | **<0.001** | 45.16 | 8.41 | 5.37 | **<0.001** | -32.50 | 7.71 | -4.22 | **<0.001** |
| E/NL | -1.88 | 9.44 | -0.20 | 0.842 | -13.41 | 7.91 | -1.70 | 0.090 | 32.88 | 8.56 | 3.84 | **<0.001** | -48.22 | 8.21 | -5.88 | **<0.001** |
| NL/V | 11.97 | 8.91 | 1.34 | 0.179 | 22.21 | 8.16 | 2.72 | **0.006** | 31.37 | 8.54 | 3.68 | **<0.001** | -28.45 | 8.14 | -3.50 | **<0.001** |
| V/NL | 1.81 | 8.79 | 0.21 | 0.837 | -5.31 | 8.16 | -0.65 | 0.515 | 38.51 | 7.91 | 4.87 | **<0.001** | -36.87 | 8.82 | -4.18 | **<0.001** |
| V/E | 16.88 | 8.56 | 1.97 | **0.049** | 6.10 | 7.81 | 0.78 | 0.435 | 45.85 | 8.46 | 5.42 | **<0.001** | -32.98 | 8.00 | -4.12 | **<0.001** |
| E/V | -16.68 | 8.88 | -1.88 | 0.060 | -2.50 | 8.48 | -0.30 | 0.768 | 37.56 | 8.38 | 4.48 | **<0.001** | -43.76 | 8.11 | -5.39 | **<0.001** |
| **Medium Force:**<br>**36.25N Elastic, -20N Viscous** | | | | | **Medium Force:**<br>**36.25N Elastic, -20N Viscous** | | | | **Medium Force:**<br>**36.25N Elastic, -20N Viscous** | | | | **Medium Force:**<br>**36.25N Elastic, -20N Viscous** | | | |
| NL/NL | 3.47 | 9.03 | 0.38 | 0.701 | 0.36 | 8.33 | 0.04 | 0.965 | 38.35 | 8.11 | 4.73 | **<0.001** | -37.57 | 8.79 | -4.28 | **<0.001** |
| V/V | -3.96 | 8.70 | -0.45 | 0.649 | -17.00 | 7.91 | -2.15 | **0.032** | 37.78 | 7.69 | 4.92 | **<0.001** | -45.11 | 8.00 | -5.64 | **<0.001** |
| E/E | 0.30 | 8.38 | 0.04 | 0.972 | -3.78 | 7.96 | -0.48 | 0.635 | 38.53 | 8.09 | 4.76 | **<0.001** | -41.73 | 7.73 | -5.40 | **<0.001** |
| NL/E | 2.96 | 9.62 | 0.31 | 0.759 | 22.16 | 8.43 | 2.63 | **0.009** | 49.90 | 8.73 | 5.72 | **<0.001** | -26.72 | 8.41 | -3.18 | **0.001** |
| E/NL | -16.98 | 8.51 | -2.00 | **0.046** | -24.87 | 7.89 | -3.15 | **0.002** | 33.82 | 8.18 | 4.13 | **<0.001** | -42.47 | 8.28 | -5.13 | **<0.001** |
| NL/V | 17.01 | 9.55 | 1.78 | 0.075 | 31.97 | 7.85 | 4.07 | **<0.001** | 30.62 | 7.83 | 3.91 | **<0.001** | -32.58 | 8.56 | -3.81 | **<0.001** |
| V/NL | 6.91 | 8.70 | 0.79 | 0.427 | -19.79 | 8.02 | -2.47 | **0.014** | 37.29 | 8.54 | 4.37 | **<0.001** | -37.69 | 8.56 | -4.40 | **<0.001** |
| V/E | 11.86 | 8.62 | 1.38 | 0.169 | 5.29 | 8.21 | 0.64 | 0.519 | 48.97 | 7.93 | 6.17 | **<0.001** | -37.83 | 8.07 | -4.69 | **<0.001** |
| E/V | -23.86 | 8.18 | -2.92 | **0.004** | -4.73 | 8.04 | -0.59 | 0.556 | 29.77 | 7.89 | 3.77 | **<0.001** | -40.72 | 7.89 | -5.16 | **<0.001** |

#### Supplementary Table 6. Between-Hand Movement Time (MT) Difference Across Loads (Relative to NL/NL)

**T6. Post-Hoc Comparisons of Movement Time (MT) Differences between Hands by Load Condition and Reach Type.** Pairwise contrasts were conducted to assess whether the difference in MT between the right and left hands varied across load conditions, using the No Load/No Load (NL/NL) condition as the reference. As is standard in treatment-versus-control comparisons, NL/NL serves as the baseline and is not shown in the table; all other estimates reflect differences relative to this reference condition.

| Movement Time (MT) | | | | | | | | | | | | | | | | |
|---|---|---|---|---|---|---|---|---|---|---|---|---|---|---|---|---|
| Symmetric Long Reach<br><br>Low Force:<br>30N Elastic, -15N Viscous | | | | | Symmetric Short Reach<br><br>Low Force:<br>30N Elastic, -15N Viscous | | | | Asymmetric Left Long. Right Short Reach<br><br>Low Force:<br>.30N Elastic, -15N Viscous | | | | Asymmetric Left Short, Right Long Reach<br><br>Low Force:<br>30N Elastic, -15N Viscous | | | |
| **Loads Applied: Left/Right Arm** | ***β*** | **SE** | **Z-Ratio** | **P Value** | ***β*** | **SE** | **Z-Ratio** | **P Value** | ***β*** | **SE** | **Z-Ratio** | **P Value** | **Estimate** | **SE** | **Z-Ratio** | **P Value** |
| V/V | -1.51 | 8.89 | -0.17 | 0.912 | -4.06 | 7.85 | -0.52 | 0.807 | -2.00 | 7.90 | -0.25 | 0.800 | -5.19 | 8.15 | -0.64 | 0.709 |
| E/E | 3.99 | 8.33 | 0.48 | 0.912 | 1.07 | 7.89 | 0.14 | 0.966 | 13.14 | 8.09 | 1.62 | 0.591 | -3.02 | 8.10 | -0.37 | 0.709 |
| NL/E | -0.97 | 8.72 | -0.11 | 0.912 | -24.33 | 7.91 | -3.08 | **0.017** | -3.08 | 8.07 | -0.38 | 0.800 | -7.24 | 8.06 | -0.90 | 0.709 |
| E/NL | 12.39 | 9.04 | 1.37 | 0.681 | 10.26 | 7.92 | 1.30 | 0.447 | 11.80 | 8.15 | 1.45 | 0.591 | 3.48 | 8.30 | 0.42 | 0.709 |
| NL/V | -2.93 | 8.77 | -0.33 | 0.912 | -19.38 | 8.03 | -2.41 | 0.063 | 3.31 | 8.12 | 0.41 | 0.800 | -9.33 | 8.26 | -1.13 | 0.709 |
| V/NL | 6.25 | 8.72 | 0.72 | 0.912 | 6.67 | 8.03 | 0.83 | 0.651 | 2.74 | 7.78 | 0.35 | 0.800 | -4.73 | 8.59 | -0.55 | 0.709 |
| V/E | -6.40 | 8.63 | -0.74 | 0.912 | -9.60 | 7.89 | -1.22 | 0.447 | -6.78 | 8.09 | -0.84 | 0.800 | -6.05 | 8.20 | -0.74 | 0.709 |
| E/V | 21.68 | 8.81 | 2.46 | 0.111 | -0.35 | 8.18 | -0.04 | 0.966 | 5.36 | 8.05 | 0.67 | 0.800 | 4.25 | 8.24 | 0.52 | 0.709 |
| **Medium Force:<br>36.25N Elastic, -20N Viscous** | | | | | **Medium Force:<br>36.25N Elastic, -20N Viscous** | | | | **Medium Force:<br>36.25N Elastic, -20N Viscous** | | | | **Medium Force:<br>36.25N Elastic, -20N Viscous** | | | |
| V/V | 9.04 | 8.66 | 1.04 | 0.464 | 15.89 | 7.90 | 2.01 | 0.071 | 3.75 | 7.69 | 0.49 | 0.828 | 4.15 | 8.19 | 0.51 | 0.975 |
| E/E | 9.92 | 8.53 | 1.16 | 0.464 | 2.37 | 7.95 | 0.30 | 0.766 | 0.94 | 7.87 | 0.12 | 0.905 | -0.25 | 8.06 | -0.03 | 0.975 |
| NL/E | 6.31 | 9.11 | 0.69 | 0.558 | -18.44 | 8.19 | -2.25 | 0.061 | -7.69 | 8.21 | -0.94 | 0.828 | -11.63 | 8.43 | -1.38 | 0.954 |
| E/NL | 21.30 | 8.60 | 2.48 | 0.053 | 22.92 | 7.91 | 2.90 | **0.015** | 6.49 | 7.94 | 0.82 | 0.828 | 1.57 | 8.31 | 0.19 | 0.975 |
| NL/V | -8.51 | 9.07 | -0.94 | 0.464 | -30.13 | 7.92 | -3.80 | **0.001** | 7.78 | 7.75 | 1.00 | 0.828 | -9.96 | 8.45 | -1.18 | 0.954 |
| V/NL | 2.03 | 8.65 | 0.24 | 0.814 | 17.22 | 7.97 | 2.16 | 0.061 | 2.86 | 8.12 | 0.35 | 0.828 | 0.89 | 8.48 | 0.11 | 0.975 |
| V/E | -8.39 | 8.66 | -0.97 | 0.464 | -3.87 | 8.08 | -0.48 | 0.722 | -3.03 | 7.83 | -0.39 | 0.828 | -4.80 | 8.22 | -0.58 | 0.975 |
| E/V | 31.23 | 8.42 | 3.71 | **0.002** | 5.12 | 7.97 | 0.64 | 0.695 | 12.70 | 7.79 | 1.63 | 0.824 | -0.44 | 8.16 | -0.05 | 0.975 |

#### Supplementary Figure 19. Total Response Time (ResT) During Reaching Movements.

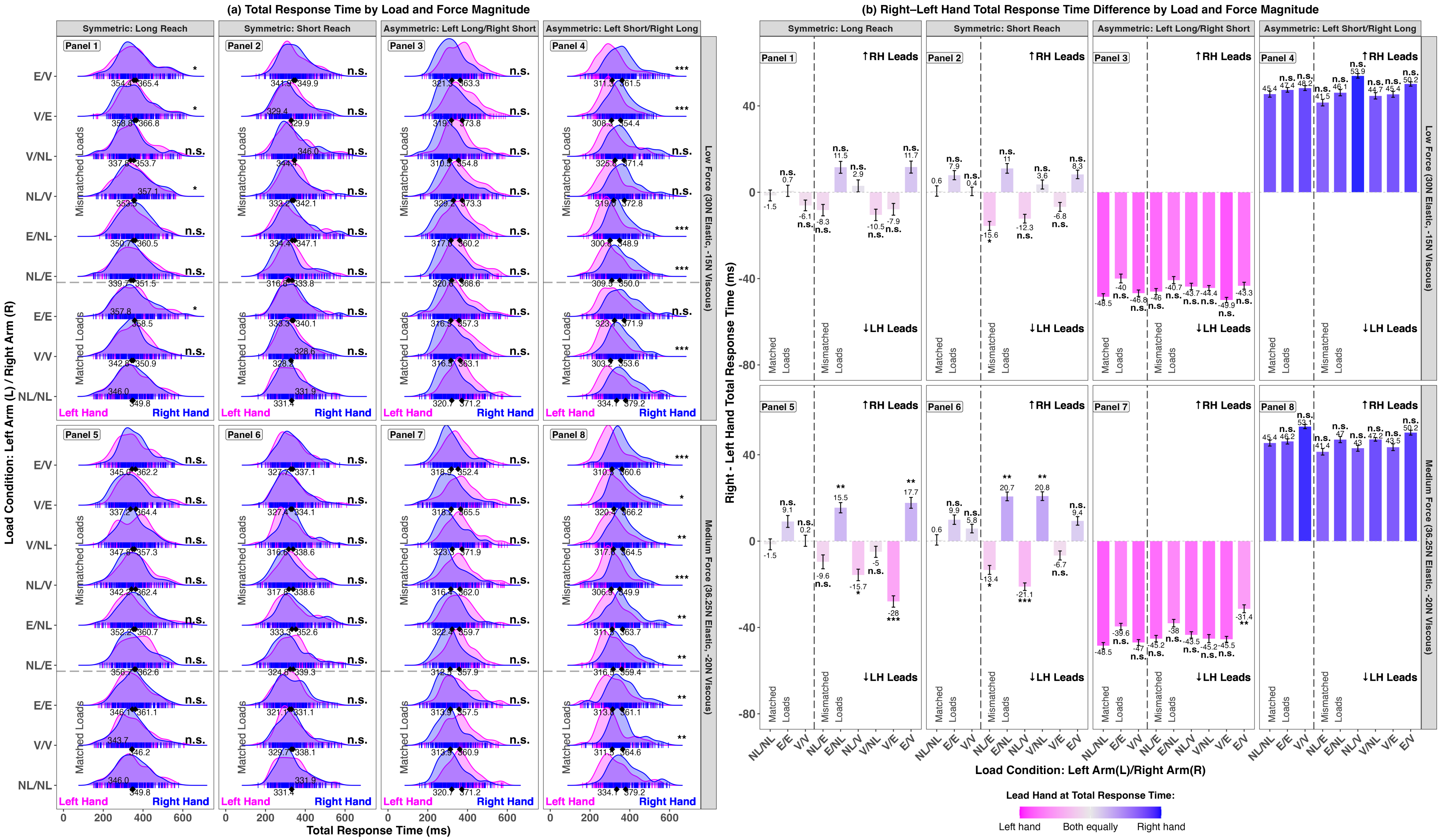


**S19. Total Response Time (ResT) During Reaching Movements.** (a) ResT across load conditions and reach types. Panels 1-4 show symmetric long, symmetric short, asymmetric left-long/right-short, and asymmetric left-short/right-long reaches under low force (30N Elastic, -15N Viscous); Panels 5-8 show the same under medium force (36.25N Elastic, -20N Viscous). The gray dashed line separates matched from mismatched loads. Asterisks indicate significance relative to the no-load (NL/NL) baseline. Ridge plots depict ResT distributions, with vertical bars for individual trials and diamonds for mean ResTs. (b) Right-left hand ResT differences (layout as in a). Positive values reflect right-hand leads; negative values reflect left-hand leads. Significance is shown relative to NL/NL. The color gradient reflects timing asymmetry: pink = left-hand lead, blue = right-hand lead, gray = synchronized.

#### Supplementary Table 7. Right-Left Hand Difference, Total Response Time (ResT)

**T7. Results from Linear Mixed-Effects Models Examining Right-Left Hand Differences in total response time (ResT).** Fixed effects estimates (β), standard errors (SE), and p-values are reported for each model, which included random intercepts for participant and trial number. Fixed effects included Load Condition, Reach Configuration, and Force Magnitude. Significant effects ($p < 0.05$) are shown in bold.

| | **Linear Mixed-Effects Model Results: Right–Left Hand Differences in Total Response Time (ResT)** | | |
|---|---|---|---|
| **Fixed effects L=left arm/R=right arm)** | ***β*** | **SE** | ***p(χ²)*** |
| Intercept (No Load (L)/No Load (R)) | -2.83 | 2.27 | 0.212 |
| Elastic (L)/Elastic (R) | 5.93 | 1.93 | **0.002** |
| Viscous (L)/Viscous (R) | 1.75 | 1.94 | 0.366 |
| No Load (L)/Elastic (R) | -5.94 | 1.96 | **0.002** |
| Elastic (L)/No Load (R) | 9.90 | 1.95 | **<0.001** |
| No Load (L)/Viscous (R) | -4.09 | 1.94 | **0.035** |
| Viscous (L)/No Load (R) | 2.49 | 1.95 | 0.203 |
| Viscous (L)/Elastic (R) | -6.08 | 1.96 | **0.002** |
| Elastic (L)/Viscous (R) | 10.11 | 1.94 | **<0.001** |
| Reach Configuration [Symmetric Short Reach] | 2.52 | 1.30 | 0.052 |
| Reach Configuration [Asymmetric Left-Long/Right-Short Reach] | -42.55 | 1.28 | **<0.001** |
| Reach Configuration [Asymmetric Left-Short/Right-Long Reach] | 47.93 | 1.29 | **<0.001** |
| Force Magnitude [Medium Force: 36.25N Elastic, -20N Viscous] | 0.94 | 0.91 | 0.302 |
| **Random effects** | **Groups** | | **SD** |
| | Trial Number | Intercept | 3.60 |
| | Participant | Intercept | 8.34 |
| | Residual | | 48.64 |
| | Observations: 11486 | | |
| **Full Model:** Right-Left Difference in ResT ~ Load Condition + Reach Configuration + Force Magnitude + (1 \| Trial Number) + (1 \| Participant) | | | |

#### Supplementary Table 8. Left/Right Pairwise Comparison, Total Response Time (ResT)

**T8. Post-Hoc Comparisons of Total Response Time (ResT) between hands.** Tukey-adjusted pairwise comparisons of left- and right-hand ResT within each load condition, across all reach types and force levels.

| Total Response Time (ResT) | | | | | | | | | | | | | | | | |
|---|---|---|---|---|---|---|---|---|---|---|---|---|---|---|---|---|
| **Symmetric Long Reach<br>Low Force:<br>30N Elastic, -15N Viscous** | | | | | **Symmetric Short Reach<br>Low Force:<br>30N Elastic, -15N Viscous** | | | | **Asymmetric Left Long. Right Short Reach<br>Low Force:<br>.30N Elastic, -15N Viscous** | | | | **Asymmetric Left Short, Right Long Reach<br>Low Force:<br>30N Elastic, -15N Viscous** | | | |
| **Loads Applied: Left/Right Arm** | **β** | **SE** | **Z-Ratio** | **P Value** | **β** | **SE** | **Z-Ratio** | **P Value** | **β** | **SE** | **Z-Ratio** | **P Value** | **Estimate** | **SE** | **Z-Ratio** | **P Value** |
| NL/NL | 3.20 | 7.56 | 0.42 | 0.671 | -0.70 | 7.74 | -0.09 | 0.928 | 50.28 | 6.96 | 7.23 | **<0.001** | -45.59 | 7.48 | -6.09 | **<0.001** |
| V/V | 7.70 | 7.23 | 1.07 | 0.287 | -0.40 | 7.27 | -0.05 | 0.956 | 46.66 | 7.31 | 6.39 | **<0.001** | -50.12 | 7.24 | -6.92 | **<0.001** |
| E/E | 0.21 | 7.24 | 0.03 | 0.976 | -7.00 | 7.15 | -0.98 | 0.328 | 40.99 | 7.17 | 5.71 | **<0.001** | -48.63 | 7.09 | -6.86 | **<0.001** |
| NL/E | 10.38 | 7.22 | 1.44 | 0.150 | 16.77 | 7.57 | 2.22 | **0.027** | 47.02 | 7.25 | 6.49 | **<0.001** | -40.66 | 6.97 | -5.84 | **<0.001** |
| E/NL | -9.97 | 7.43 | -1.34 | 0.180 | -12.37 | 7.40 | -1.67 | 0.095 | 43.08 | 7.35 | 5.86 | **<0.001** | -47.97 | 7.36 | -6.52 | **<0.001** |
| NL/V | -2.49 | 7.28 | -0.34 | 0.732 | 10.00 | 7.34 | 1.36 | 0.173 | 43.95 | 7.25 | 6.06 | **<0.001** | -53.39 | 7.55 | -7.07 | **<0.001** |
| V/NL | 14.67 | 7.58 | 1.94 | 0.053 | -1.42 | 7.57 | -0.19 | 0.851 | 44.37 | 7.06 | 6.29 | **<0.001** | -45.37 | 7.42 | -6.11 | **<0.001** |
| V/E | 6.77 | 7.43 | 0.91 | 0.362 | 2.40 | 7.52 | 0.32 | 0.750 | 52.72 | 7.33 | 7.19 | **<0.001** | -46.66 | 7.35 | -6.35 | **<0.001** |
| E/V | -10.31 | 7.41 | -1.39 | 0.164 | -8.34 | 7.29 | -1.14 | 0.253 | 41.73 | 7.13 | 5.85 | **<0.001** | -49.36 | 7.36 | -6.71 | **<0.001** |
| **Medium Force:<br>36.25N Elastic, -20N Viscous** | | | | | **Medium Force:<br>36.25N Elastic, -20N Viscous** | | | | **Medium Force:<br>36.25N Elastic, -20N Viscous** | | | | **Medium Force:<br>36.25N Elastic, -20N Viscous** | | | |
| NL/NL | 3.20 | 7.56 | 0.42 | 0.671 | -0.70 | 7.74 | -0.09 | 0.928 | 50.28 | 6.96 | 7.23 | **<0.001** | -45.59 | 7.48 | -6.09 | **<0.001** |
| V/V | 1.50 | 7.23 | 0.21 | 0.835 | -6.43 | 7.32 | -0.88 | 0.380 | 46.98 | 7.10 | 6.62 | **<0.001** | -53.10 | 7.38 | -7.19 | **<0.001** |
| E/E | -14.42 | 7.13 | -2.02 | **0.043** | -9.59 | 7.51 | -1.28 | 0.201 | 42.58 | 7.22 | 5.90 | **<0.001** | -47.31 | 7.17 | -6.60 | **<0.001** |
| NL/E | 6.52 | 7.64 | 0.85 | 0.394 | 14.60 | 7.54 | 1.94 | 0.053 | 45.52 | 7.66 | 5.95 | **<0.001** | -42.61 | 7.34 | -5.81 | **<0.001** |
| E/NL | -9.60 | 7.25 | -1.32 | 0.186 | -19.94 | 7.26 | -2.75 | **0.006** | 37.40 | 7.18 | 5.21 | **<0.001** | -50.75 | 7.36 | -6.89 | **<0.001** |
| NL/V | 19.31 | 7.23 | 2.67 | **0.008** | 20.41 | 7.08 | 2.88 | **0.004** | 45.32 | 7.06 | 6.42 | **<0.001** | -43.00 | 7.32 | -5.88 | **<0.001** |
| V/NL | 9.32 | 7.26 | 1.28 | 0.199 | -21.65 | 7.40 | -2.93 | **0.003** | 47.67 | 7.25 | 6.58 | **<0.001** | -47.21 | 7.29 | -6.47 | **<0.001** |
| V/E | 26.56 | 7.46 | 3.56 | **<0.001** | 6.58 | 7.46 | 0.88 | 0.378 | 47.02 | 7.32 | 6.43 | **<0.001** | -45.51 | 7.41 | -6.14 | **<0.001** |
| E/V | -18.11 | 7.23 | -2.51 | **0.012** | -9.37 | 7.10 | -1.32 | 0.187 | 32.87 | 7.19 | 4.57 | **<0.001** | -50.46 | 7.23 | -6.98 | **<0.001** |

#### Supplementary Table 9. Between-Hand Total Response Time (ResT) Difference Across Loads (Relative to NL/NL)

**T9. Post-Hoc Comparisons of Total Response Time (ResT) Differences between Hands by Load Condition and Reach Type.** Pairwise contrasts were conducted to assess whether the difference in ResT between the right and left hands varied across load conditions, using the No Load/No Load (NL/NL) condition as the reference. As is standard in treatment-versus-control comparisons, NL/NL serves as the baseline and is not shown in the table; all other estimates reflect differences relative to this reference condition.

| Total Response Time (ResT) | | | | | | | | | | | | | | | | |
|---|---|---|---|---|---|---|---|---|---|---|---|---|---|---|---|---|
| | Symmetric Long Reach<br><br>Low Force:<br>30N Elastic, -15N Viscous | | | | Symmetric Short Reach<br><br>Low Force:<br>30N Elastic, -15N Viscous | | | | Asymmetric Left Long. Right Short Reach<br><br>Low Force:<br>.30N Elastic, -15N Viscous | | | | Asymmetric Left Short, Right Long Reach<br><br>Low Force:<br>30N Elastic, -15N Viscous | | | |
| **Loads Applied: Left/Right Arm** | ***β*** | **SE** | **Z-Ratio** | **P Value** | ***β*** | **SE** | **Z-Ratio** | **P Value** | ***β*** | **SE** | **Z-Ratio** | **P Value** | **Estimate** | **SE** | **Z-Ratio** | **P Value** |
| V/V | -5.35 | 5.49 | -0.97 | 0.441 | -1.05 | 5.55 | -0.19 | 0.850 | 1.30 | 5.28 | 0.25 | 0.805 | 2.72 | 5.45 | 0.50 | 0.970 |
| E/E | 0.83 | 5.49 | 0.15 | 0.880 | 6.53 | 5.51 | 1.19 | 0.314 | 8.49 | 5.23 | 1.62 | 0.714 | 1.91 | 5.39 | 0.35 | 0.970 |
| NL/E | -7.65 | 5.49 | -1.39 | 0.326 | -16.42 | 5.66 | -2.90 | **0.030** | 2.53 | 5.25 | 0.48 | 0.805 | -4.04 | 5.34 | -0.76 | 0.970 |
| E/NL | 13.32 | 5.58 | 2.39 | 0.087 | 9.04 | 5.60 | 1.61 | 0.269 | 7.14 | 5.31 | 1.35 | 0.714 | 0.70 | 5.50 | 0.13 | 0.970 |
| NL/V | 3.52 | 5.52 | 0.64 | 0.599 | -13.20 | 5.58 | -2.37 | 0.072 | 4.17 | 5.25 | 0.79 | 0.725 | 8.32 | 5.56 | 1.50 | 0.970 |
| V/NL | -8.90 | 5.63 | -1.58 | 0.303 | 3.10 | 5.66 | 0.55 | 0.668 | 3.88 | 5.17 | 0.75 | 0.725 | -0.73 | 5.51 | -0.13 | 0.970 |
| V/E | -6.47 | 5.57 | -1.16 | 0.393 | -8.44 | 5.67 | -1.49 | 0.269 | -2.01 | 5.30 | −0.38 | 0.805 | 0.21 | 5.50 | 0.04 | 0.970 |
| E/V | 12.77 | 5.56 | 2.30 | 0.087 | 7.67 | 5.56 | 1.38 | 0.269 | 5.24 | 5.21 | 1.01 | 0.725 | 5.18 | 5.49 | 0.94 | 0.970 |
| **Medium Force:<br>36.25N Elastic, -20N Viscous** | | | | | **Medium Force:<br>36.25N Elastic, -20N Viscous** | | | | **Medium Force:<br>36.25N Elastic, -20N Viscous** | | | | **Medium Force:<br>36.25N Elastic, -20N Viscous** | | | |
| V/V | 1.43 | 5.49 | 0.26 | 0.795 | 5.09 | 5.58 | 0.91 | 0.362 | 1.27 | 5.19 | 0.24 | 0.807 | 7.73 | 5.49 | 1.41 | 0.857 |
| E/E | 10.48 | 5.47 | 1.92 | 0.089 | 8.80 | 5.64 | 1.56 | 0.154 | 8.65 | 5.25 | 1.65 | 0.266 | 0.79 | 5.42 | 0.15 | 0.884 |
| NL/E | -7.98 | 5.65 | -1.41 | 0.211 | -14.54 | 5.66 | -2.57 | **0.020** | 2.13 | 5.41 | 0.39 | 0.793 | -4.02 | 5.48 | -0.73 | 0.857 |
| E/NL | 16.28 | 5.51 | 2.95 | **0.008** | 19.33 | 5.56 | 3.47 | **0.001** | 10.22 | 5.23 | 1.96 | 0.202 | 1.75 | 5.50 | 0.32 | 0.857 |
| NL/V | -14.65 | 5.51 | -2.66 | 0.016 | -22.12 | 5.48 | -4.03 | **<0.001** | 4.12 | 5.18 | 0.80 | 0.746 | -2.64 | 5.46 | -0.48 | 0.857 |
| V/NL | -3.69 | 5.51 | -0.67 | 0.575 | 19.44 | 5.59 | 3.47 | **0.001** | 3.07 | 5.26 | 0.58 | 0.746 | 2.31 | 5.47 | 0.42 | 0.857 |
| V/E | -26.47 | 5.59 | -4.74 | **<0.001** | -8.49 | 5.62 | -1.51 | 0.154 | 3.19 | 5.29 | 0.60 | 0.746 | -2.19 | 5.51 | -0.40 | 0.857 |
| E/V | 19.22 | 5.50 | 3.49 | **0.002** | 8.20 | 5.48 | 1.49 | 0.154 | 17.34 | 5.24 | 3.31 | **0.007** | 4.46 | 5.44 | 0.82 | 0.857 |

#### Supplementary Table 10. Left-Right Hand Difference LME Model, Movement End Time (ME)

**T10. Results from Linear Mixed-Effects Models Examining Right-Left Hand Differences in Movement End Time (ME).** Fixed effects estimates (β), standard errors (SE), and p-values are reported for each model, which included random intercepts for participant and trial number. Fixed effects included Load Condition, Reach Configuration, and Force Magnitude. Significant effects ($p < 0.05$) are shown in bold.

| | **Linear Mixed-Effects Model Results: Right–Left Hand Differences in Movement End Time (ME)** | | |
|---|---|---|---|
| **Fixed effects L=left arm/R=right arm)** | ***β*** | **SE** | ***p(χ²)*** |
| Intercept (No Load (L)/No Load (R)) | 19.67 | 12.22 | 0.107 |
| Elastic (L)/Elastic (R) | 33.57 | 8.45 | **<0.001** |
| Viscous (L)/Viscous (R) | 11.49 | 8.42 | 0.173 |
| No Load (L)/Elastic (R) | -65.14 | 8.44 | **<0.001** |
| Elastic (L)/No Load (R) | 102.90 | 8.45 | **<0.001** |
| No Load (L)/Viscous (R) | 1.99 | 8.46 | 0.814 |
| Viscous (L)/No Load (R) | 10.13 | 8.38 | 0.227 |
| Viscous (L)/Elastic (R) | -73.07 | 8.43 | **<0.001** |
| Elastic (L)/Viscous (R) | 138.45 | 8.46 | **<0.001** |
| Reach Configuration [Symmetric Short Reach] | -30.81 | 5.62 | **<0.001** |
| Reach Configuration [Asymmetric Left-Long/Right-Short Reach] | -431.96 | 5.64 | **<0.001** |
| Reach Configuration [Asymmetric Left-Short/Right-Long Reach] | 368.97 | 5.64 | **<0.001** |
| Force Magnitude [Medium Force: 36.25N Elastic, -20N Viscous] | 1.41 | 3.98 | 0.724 |
| **Random effects** | **Groups** | | **SD** |
| | Trial Number | Intercept | 22.22 |
| | Participant | Intercept | 54.01 |
| | Residual | | 280.87 |
| | Observations: 19997 | | |
| **Full Model:** ME ~ (Load Condition x Hand) + Reach Configuration + Force Magnitude + (1 \| Trial Number) + (1 \| Participant) | | | |

#### Supplementary Table 11. Left/Right Pairwise Comparison, Movement End Time (ME)

**T11. Post-Hoc Comparisons of Movement End Time (ME) between hands.** Tukey-adjusted pairwise comparisons of left- and right-hand ME within each load condition, across all reach types and force levels.

| Movement End Time (ME) | | | | | | | | | | | | | | | | |
|---|---|---|---|---|---|---|---|---|---|---|---|---|---|---|---|---|
| **Symmetric Long Reach**<br><br>**Low Force:**<br>**30N Elastic, -15N Viscous** | | | | | **Symmetric Short Reach**<br><br>**Low Force:**<br>**30N Elastic, -15N Viscous** | | | | **Asymmetric Left Long. Right Short Reach**<br><br>**Low Force:**<br>**.30N Elastic, -15N Viscous** | | | | **Asymmetric Left Short, Right Long Reach**<br><br>**Low Force:**<br>**30N Elastic, -15N Viscous** | | | |
| **Loads Applied: Left/Right Arm** | ***β*** | **SE** | **Z-Ratio** | **P Value** | ***β*** | **SE** | **Z-Ratio** | **P Value** | ***β*** | **SE** | **Z-Ratio** | **P Value** | **Estimate** | **SE** | **Z-Ratio** | **P Value** |
| NL/NL | -4.52 | 17.83 | -0.25 | 0.800 | 11.15 | 17.93 | 0.62 | 0.534 | 396.32 | 17.80 | 22.26 | **<0.001** | -382.69 | 17.79 | -21.51 | **<0.001** |
| V/V | 2.09 | 17.64 | 0.12 | 0.906 | -6.28 | 17.93 | -0.35 | 0.726 | 409.98 | 17.79 | 23.05 | **<0.001** | -421.25 | 17.95 | -23.47 | **<0.001** |
| E/E | -32.49 | 17.88 | -1.82 | 0.069 | -28.32 | 17.91 | -1.58 | 0.114 | 392.81 | 18.25 | 21.52 | **<0.001** | -444.53 | 17.99 | -24.71 | **<0.001** |
| NL/E | 23.30 | 17.96 | 1.30 | 0.195 | 52.75 | 17.73 | 2.98 | **0.003** | 427.52 | 17.87 | 23.93 | **<0.001** | -287.32 | 18.13 | -15.85 | **<0.001** |
| E/NL | -96.79 | 17.91 | -5.40 | **<0.001** | -72.60 | 17.76 | -4.09 | **<0.001** | 308.58 | 17.98 | 17.17 | **<0.001** | -467.63 | 17.83 | -26.22 | **<0.001** |
| NL/V | -74.67 | 18.02 | -4.14 | **<0.001** | 14.41 | 18.03 | 0.80 | 0.424 | 349.38 | 17.94 | 19.47 | **<0.001** | -389.35 | 17.94 | -21.70 | **<0.001** |
| V/NL | 32.96 | 17.73 | 1.86 | 0.063 | -31.53 | 17.80 | -1.77 | 0.077 | 415.59 | 17.70 | 23.48 | **<0.001** | -373.39 | 17.94 | -20.81 | **<0.001** |
| V/E | 68.66 | 18.01 | 3.81 | **<0.001** | 59.46 | 17.83 | 3.33 | **<0.001** | 508.40 | 17.76 | 28.63 | **<0.001** | -371.11 | 17.87 | -20.77 | **<0.001** |
| E/V | -197.27 | 17.98 | -10.97 | **<0.001** | -82.25 | 18.10 | -4.54 | **<0.001** | 322.87 | 17.96 | 17.98 | **<0.001** | -544.67 | 17.84 | -30.54 | **<0.001** |
| **Medium Force:**<br>**36.25N Elastic, -20N Viscous** | | | | | **Medium Force:**<br>**36.25N Elastic, -20N Viscous** | | | | **Medium Force:**<br>**36.25N Elastic, -20N Viscous** | | | | **Medium Force:**<br>**36.25N Elastic, -20N Viscous** | | | |
| NL/NL | -4.52 | 17.83 | -0.25 | 0.800 | 11.15 | 17.93 | 0.62 | 0.534 | 396.32 | 17.80 | 22.26 | **<0.001** | -382.69 | 17.79 | -21.51 | **<0.001** |
| V/V | -43.68 | 18.02 | -2.42 | **0.015** | -16.73 | 17.93 | -0.93 | 0.351 | 427.68 | 17.89 | 23.91 | **<0.001** | -421.30 | 18.02 | -23.38 | **<0.001** |
| E/E | -105.55 | 17.90 | -5.90 | **<0.001** | -21.37 | 17.76 | -1.20 | 0.229 | 449.28 | 18.07 | 24.86 | **<0.001** | -467.33 | 17.84 | -26.20 | **<0.001** |
| NL/E | 70.14 | 17.98 | 3.90 | **<0.001** | 93.43 | 17.77 | 5.26 | **<0.001** | 489.99 | 18.02 | 27.18 | **<0.001** | -318.18 | 17.85 | -17.82 | **<0.001** |
| E/NL | -134.05 | 17.99 | -7.45 | **<0.001** | -111.38 | 17.93 | -6.21 | **<0.001** | 264.43 | 17.99 | 14.70 | **<0.001** | -496.56 | 17.98 | -27.62 | **<0.001** |
| NL/V | -45.99 | 18.06 | -2.55 | **<0.001** | 78.72 | 17.94 | 4.39 | **<0.001** | 383.21 | 18.01 | 21.28 | **<0.001** | -335.72 | 17.90 | -18.76 | **<0.001** |
| V/NL | 18.53 | 17.59 | 1.05 | 0.292 | -69.14 | 17.80 | -3.88 | **<0.001** | 360.06 | 17.68 | 20.36 | **<0.001** | -403.86 | 17.96 | -22.48 | **<0.001** |
| V/E | 127.66 | 18.01 | 7.09 | **<0.001** | 42.60 | 17.83 | 2.39 | **0.017** | 529.83 | 17.79 | 29.79 | **<0.001** | -326.96 | 18.06 | -18.11 | **<0.001** |
| E/V | -286.13 | 18.01 | -15.89 | **<0.001** | -61.93 | 17.99 | -3.44 | **<0.001** | 282.94 | 18.03 | 15.70 | **<0.001** | -574.52 | 17.87 | -32.16 | **<0.001** |

#### Supplementary Table 12. Between-Hand Movement End Time (ME) Difference Across Loads (Relative to NL/NL)

**T12. Post-Hoc Comparisons of Movement End Time (ME) Differences between Hands by Load Condition and Reach Type.** Pairwise comparisons assess whether hand differences in ME significantly vary across load conditions, using the No Load/No Load condition as the baseline.

| **Movement End Time (ME)** | | | | | | | | | | | | | | | | |
|---|---|---|---|---|---|---|---|---|---|---|---|---|---|---|---|---|
| **Symmetric Long Reach<br><br>Low Force:<br>30N Elastic, -15N Viscous** | | | | | **Symmetric Short Reach<br><br>Low Force:<br>30N Elastic, -15N Viscous** | | | | **Asymmetric Left Long. Right Short Reach<br><br>Low Force:<br>.30N Elastic, -15N Viscous** | | | | **Asymmetric Left Short, Right Long Reach<br><br>Low Force:<br>30N Elastic, -15N Viscous** | | | |
| **Loads Applied: Left/Right Arm** | ***β*** | **SE** | **Z-Ratio** | **P Value** | ***β*** | **SE** | **Z-Ratio** | **P Value** | ***β*** | **SE** | **Z-Ratio** | **P Value** | **Estimate** | **SE** | **Z-Ratio** | **P Value** |
| V/V | -4.13 | 23.28 | -0.18 | 0.859 | 23.31 | 23.68 | 0.98 | 0.371 | -23.46 | 23.44 | -1.00 | 0.362 | 32.52 | 23.65 | 1.37 | 0.271 |
| E/E | 20.70 | 23.52 | 0.88 | 0.433 | 46.65 | 23.61 | 1.98 | 0.096 | 1.97 | 24.04 | 0.08 | 0.935 | 56.53 | 23.72 | 2.38 | **0.034** |
| NL/E | -33.26 | 23.68 | -1.40 | 0.214 | -40.82 | 23.47 | -1.74 | 0.131 | -33.11 | 23.63 | -1.40 | 0.258 | -97.09 | 23.92 | -4.06 | **<0.001** |
| E/NL | 82.65 | 23.63 | 3.50 | **0.002** | 95.39 | 23.53 | 4.05 | **<0.001** | 81.67 | 23.75 | 3.44 | **0.002** | 80.58 | 23.55 | 3.42 | **0.002** |
| NL/V | 58.33 | 23.70 | 2.46 | **0.028** | -3.18 | 23.79 | -0.13 | 0.894 | 43.54 | 23.72 | 1.84 | 0.133 | 4.92 | 23.61 | 0.21 | 0.835 |
| V/NL | -39.56 | 23.38 | -1.69 | 0.145 | 48.38 | 23.55 | 2.05 | 0.096 | -25.13 | 23.38 | -1.07 | 0.362 | -5.95 | 23.63 | -0.25 | 0.835 |
| V/E | -71.50 | 23.72 | -3.01 | **0.007** | -38.89 | 23.57 | -1.65 | 0.132 | -118.80 | 23.46 | -5.06 | **<0.001** | -10.76 | 23.58 | -0.46 | 0.835 |
| E/V | 180.24 | 23.65 | 7.62 | **<0.001** | 95.74 | 23.81 | 4.02 | **<0.001** | 61.27 | 23.72 | 2.58 | **0.026** | 147.86 | 23.52 | 6.29 | **<0.001** |
| **Medium Force:<br>36.25N Elastic, -20N Viscous** | | | | | **Medium Force:<br>36.25N Elastic, -20N Viscous** | | | | **Medium Force:<br>36.25N Elastic, -20N Viscous** | | | | **Medium Force:<br>36.25N Elastic, -20N Viscous** | | | |
| V/V | 29.02 | 23.76 | 1.22 | 0.254 | 31.89 | 23.67 | 1.35 | 0.203 | -30.96 | 23.61 | -1.31 | 0.253 | 35.73 | 23.68 | 1.51 | 0.150 |
| E/E | 87.35 | 23.63 | 3.70 | **<0.001** | 39.36 | 23.43 | 1.68 | 0.124 | -68.98 | 23.85 | -2.89 | **0.006** | 81.71 | 23.57 | 3.47 | **0.001** |
| NL/E | -76.89 | 23.65 | -3.25 | **0.002** | -81.24 | 23.46 | -3.46 | **0.001** | -99.40 | 23.79 | -4.18 | **<0.001** | -61.49 | 23.53 | -2.61 | **0.018** |
| E/NL | 124.64 | 23.74 | 5.25 | **<0.001** | 128.36 | 23.67 | 5.42 | **<0.001** | 120.18 | 23.77 | 5.06 | **<0.001** | 111.31 | 23.73 | 4.69 | **<0.001** |
| NL/V | 18.98 | 23.75 | 0.80 | 0.424 | -63.92 | 23.70 | -2.70 | **0.011** | 5.38 | 23.69 | 0.23 | 0.820 | -46.37 | 23.57 | -1.97 | 0.065 |
| V/NL | -29.69 | 23.22 | -1.28 | 0.254 | 87.70 | 23.59 | 3.72 | **<0.001** | 26.99 | 23.36 | 1.16 | 0.284 | 20.58 | 23.70 | 0.87 | 0.385 |
| V/E | -134.75 | 23.70 | -5.69 | **<0.001** | -21.36 | 23.59 | -0.91 | 0.365 | -134.52 | 23.49 | -5.73 | **<0.001** | -55.07 | 23.77 | -2.32 | **0.033** |
| E/V | 260.35 | 23.74 | 10.97 | **<0.001** | 76.59 | 23.72 | 3.23 | **0.002** | 103.25 | 23.79 | 4.34 | **<0.001** | 182.02 | 23.59 | 7.71 | **<0.001** |

#### Supplementary Table 13. Left-Right Hand Difference LME Model, Peak Velocity & Time at Peak Velocity

**T13. Results from Linear Mixed-Effects Models Examining Right-Left Hand Differences in (a) Peak Velocity (PV) and (b) Time to Peak Velocity (TPV).** Fixed effects estimates (β), standard errors (SE), and p-values are reported for each model, which included random intercepts for participant and trial number. Fixed effects included Load Condition, Reach Configuration, and Force Magnitude. Significant effects ($p < 0.05$) are shown in bold.

| | **Linear Mixed-Effects Model Results: Right–Left Hand Differences in Peak Velocity (PV) and Time to Peak Velocity (TPV)** | | | | | |
|---|---|---|---|---|---|---|
| | **(a) PV** | | | **(b) TPV** | | |
| **Fixed effects L=left arm/R=right arm)** | ***β*** | **SE** | ***p(χ²)*** | ***β*** | **SE** | ***p(χ²)*** |
| Intercept (No Load (L)/No Load (R)) | -5.82 | 0.94 | **<0.001** | 1.90 | 9.10 | 0.834 |
| Elastic (L)/Elastic (R) | 0.27 | 0.52 | 0.606 | -1.17 | 5.05 | 0.817 |
| Viscous (L)/Viscous (R) | 1.75 | 0.51 | **0.001** | 2.48 | 5.05 | 0.623 |
| No Load (L)/Elastic (R) | 20.43 | 0.52 | **<0.001** | -50.33 | 5.11 | **<0.001** |
| Elastic (L)/No Load (R) | -20.23 | 0.52 | **<0.001** | 39.95 | 5.10 | **<0.001** |
| No Load (L)/Viscous (R) | 25.39 | 0.52 | **<0.001** | -193.54 | 5.10 | **<0.001** |
| Viscous (L)/No Load (R) | -22.87 | 0.52 | **<0.001** | 187.53 | 5.09 | **<0.001** |
| Viscous (L)/Elastic (R) | -3.50 | 0.52 | **<0.001** | 135.65 | 5.06 | **<0.001** |
| Elastic (L)/Viscous (R) | 5.59 | 0.52 | **<0.001** | -144.35 | 5.05 | **<0.001** |
| Reach Configuration [Symmetric Short Reach] | 1.45 | 0.34 | **<0.001** | -2.61 | 2.98 | 0.381 |
| Reach Configuration [Asymmetric Left-Long/Right-Short Reach] | -31.69 | 0.34 | **<0.001** | -138.95 | 3.04 | **<0.001** |
| Reach Configuration [Asymmetric Left-Short/Right-Long Reach] | 33.67 | 0.34 | **<0.001** | 124.99 | 3.03 | **<0.001** |
| Force Magnitude [Medium Force: 36.25N Elastic, -20N Viscous] | 0.17 | 0.24 | 0.491 | -2.77 | 2.12 | 0.191 |
| **Random effects** | **Groups** | | **SD** | **Groups** | | **SD** |
| | Trial Number | Intercept | 1.46 | Trial Number | Intercept | 3.29 |
| | Participant | Intercept | 4.52 | Participant | Intercept | 43.17 |
| | Residual | | 16.65 | Residual | | 144.49 |
| | Observations: 19,295 | | | Observations: 18,583 | | |
| **Full Model:** Right-Left Difference in PV or TPV ~ Load Condition + Reach Configuration + Force Magnitude + (1 \| Trial Number) + (1 \| Participant) | | | | | | |

#### Supplementary Table 14. Left/Right Pairwise Comparison, Peak Velocity (PV)

**T14. Post-Hoc Comparisons of Peak Velocity (PV) between hands.** Tukey-adjusted pairwise comparisons of left- and right-hand PV within each load condition, across all reach types and force levels.

| **Peak Velocity (PV)** | | | | | | | | | | | | | | | | |
|---|---|---|---|---|---|---|---|---|---|---|---|---|---|---|---|---|
| **Symmetric Long Reach<br><br>Low Force:<br>30N Elastic, -15N Viscous** | | | | | **Symmetric Short Reach<br><br>Low Force:<br>30N Elastic, -15N Viscous** | | | | **Asymmetric Left Long. Right Short Reach<br><br>Low Force:<br>.30N Elastic, -15N Viscous** | | | | **Asymmetric Left Short, Right Long Reach<br><br>Low Force:<br>30N Elastic, -15N Viscous** | | | |
| **Loads Applied: Left/Right Arm** | ***β*** | **SE** | **Z-Ratio** | **P Value** | ***β*** | **SE** | **Z-Ratio** | **P Value** | ***β*** | **SE** | **Z-Ratio** | **P Value** | **Estimate** | **SE** | **Z-Ratio** | **P Value** |
| NL/NL | 4.63 | 1.23 | 3.77 | **<0.001** | 5.72 | 1.10 | 5.22 | **<0.001** | 40.35 | 1.14 | 35.37 | **<0.001** | -30.86 | 1.16 | -26.63 | **<0.001** |
| V/V | 4.59 | 1.10 | 4.19 | **<0.001** | 2.38 | 1.10 | 2.16 | **0.031** | 36.59 | 1.10 | 33.35 | **<0.001** | -28.92 | 1.11 | -26.06 | **<0.001** |
| E/E | 6.35 | 1.11 | 5.71 | **<0.001** | 3.78 | 1.10 | 3.44 | **<0.001** | 37.89 | 1.10 | 34.39 | **<0.001** | -27.52 | 1.09 | -25.20 | **<0.001** |
| NL/E | -17.40 | 1.18 | -14.76 | **<0.001** | -12.23 | 1.10 | -11.15 | **<0.001** | 20.14 | 1.10 | 18.26 | **<0.001** | -49.78 | 1.15 | -43.11 | **<0.001** |
| E/NL | 28.92 | 1.18 | 24.51 | **<0.001** | 21.27 | 1.10 | 19.32 | **<0.001** | 56.01 | 1.14 | 48.96 | **<0.001** | -9.84 | 1.10 | -8.93 | **<0.001** |
| NL/V | -17.36 | 1.14 | -15.20 | **<0.001** | -18.89 | 1.10 | -17.16 | **<0.001** | 17.29 | 1.10 | 15.68 | **<0.001** | -53.19 | 1.15 | -46.22 | **<0.001** |
| V/NL | 27.32 | 1.14 | 23.91 | **<0.001** | 28.02 | 1.10 | 25.58 | **<0.001** | 59.08 | 1.14 | 51.85 | **<0.001** | -9.02 | 1.10 | -8.17 | **<0.001** |
| V/E | 4.90 | 1.11 | 4.42 | **<0.001** | 8.08 | 1.10 | 7.38 | **<0.001** | 39.09 | 1.11 | 35.29 | **<0.001** | -23.49 | 1.10 | -21.34 | **<0.001** |
| E/V | 5.15 | 1.11 | 4.65 | **<0.001** | -2.58 | 1.10 | -2.36 | **0.019** | 32.45 | 1.10 | 29.53 | **<0.001** | -34.25 | 1.11 | -30.92 | **<0.001** |
| **Medium Force:<br>36.25N Elastic, -20N Viscous** | | | | | **Medium Force:<br>36.25N Elastic, -20N Viscous** | | | | **Medium Force:<br>36.25N Elastic, -20N Viscous** | | | | **Medium Force:<br>36.25N Elastic, -20N Viscous** | | | |
| NL/NL | 4.63 | 1.23 | 3.77 | **<0.001** | 5.72 | 1.10 | 5.22 | **<0.001** | 40.35 | 1.14 | 35.37 | **<0.001** | -30.86 | 1.16 | -26.63 | **<0.001** |
| V/V | 3.08 | 1.10 | 2.81 | **0.005** | 2.49 | 1.10 | 2.27 | **0.023** | 33.43 | 1.09 | 30.55 | **<0.001** | -26.93 | 1.10 | -24.46 | **<0.001** |
| E/E | 4.73 | 1.10 | 4.29 | **<0.001** | 4.24 | 1.10 | 3.86 | **<0.001** | 34.38 | 1.10 | 31.23 | **<0.001** | -25.97 | 1.10 | -23.51 | **<0.001** |
| NL/E | -19.15 | 1.18 | -16.26 | **<0.001** | -16.58 | 1.10 | -15.11 | **<0.001** | 18.32 | 1.10 | 16.65 | **<0.001** | -49.47 | 1.17 | -42.32 | **<0.001** |
| E/NL | 31.81 | 1.17 | 27.17 | **<0.001** | 23.66 | 1.10 | 21.57 | **<0.001** | 58.04 | 1.16 | 50.15 | **<0.001** | -7.65 | 1.11 | -6.91 | **<0.001** |
| NL/V | -23.97 | 1.16 | -20.64 | **<0.001** | -23.13 | 1.10 | -20.98 | **<0.001** | 11.90 | 1.10 | 10.84 | **<0.001** | -57.95 | 1.15 | -50.56 | **<0.001** |
| V/NL | 30.87 | 1.14 | 27.04 | **<0.001** | 27.88 | 1.10 | 25.41 | **<0.001** | 62.04 | 1.14 | 54.34 | **<0.001** | -3.75 | 1.10 | -3.41 | **<0.001** |
| V/E | 7.72 | 1.10 | 7.01 | **<0.001** | 8.59 | 1.09 | 7.85 | **<0.001** | 41.43 | 1.10 | 37.66 | **<0.001** | -18.89 | 1.11 | -17.06 | **<0.001** |
| E/V | 2.09 | 1.10 | 1.89 | 0.058 | -4.16 | 1.10 | -3.77 | **<0.001** | 29.60 | 1.10 | 27.00 | **<0.001** | -33.12 | 1.12 | -29.63 | **<0.001** |

#### Supplementary Table 15. Between-Hand Peak Velocity (PV) Difference Across Loads (Relative to NL/NL)

**T15. Post-Hoc Comparisons of Peak Velocity (PV) Differences between Hands by Load Condition and Reach Type.** Pairwise comparisons assess whether hand differences in PV significantly vary across load conditions, using the No Load/No Load condition as the baseline.

| Peak Velocity (PV) | | | | | | | | | | | | | | | | |
|---|---|---|---|---|---|---|---|---|---|---|---|---|---|---|---|---|
| Symmetric Long Reach<br><br>Low Force:<br>30N Elastic, -15N Viscous | | | | | Symmetric Short Reach<br><br>Low Force:<br>30N Elastic, -15N Viscous | | | | Asymmetric Left Long. Right Short Reach<br><br>Low Force:<br>.30N Elastic, -15N Viscous | | | | Asymmetric Left Short, Right Long Reach<br><br>Low Force:<br>30N Elastic, -15N Viscous | | | |
| Loads Applied: Left/Right Arm | *β* | SE | Z-Ratio | P Value | *β* | SE | Z-Ratio | P Value | *β* | SE | Z-Ratio | P Value | Estimate | SE | Z-Ratio | P Value |
| V/V | 0.56 | 1.48 | 0.38 | 0.940 | 3.40 | 1.40 | 2.42 | **0.021** | 3.81 | 1.43 | 2.67 | **0.010** | -2.19 | 1.45 | -1.51 | 0.132 |
| E/E | -1.06 | 1.49 | -0.71 | 0.766 | 1.80 | 1.41 | 1.28 | 0.200 | 2.46 | 1.43 | 1.72 | 0.098 | -4.05 | 1.44 | -2.81 | **0.007** |
| NL/E | 22.18 | 1.53 | 14.48 | **<0.001** | 17.90 | 1.40 | 12.76 | **<0.001** | 19.83 | 1.44 | 13.80 | **<0.001** | 18.19 | 1.47 | 12.34 | **<0.001** |
| E/NL | -23.59 | 1.53 | -15.37 | **<0.001** | -15.75 | 1.41 | -11.19 | **<0.001** | -15.95 | 1.46 | -10.95 | **<0.001** | -21.66 | 1.45 | -14.95 | **<0.001** |
| NL/V | 22.17 | 1.51 | 14.69 | **<0.001** | 24.48 | 1.41 | 17.39 | **<0.001** | 22.81 | 1.43 | 15.93 | **<0.001** | 22.06 | 1.47 | 14.96 | **<0.001** |
| V/NL | -21.94 | 1.51 | -14.49 | **<0.001** | -22.27 | 1.41 | -15.83 | **<0.001** | -19.13 | 1.45 | -13.18 | **<0.001** | -22.08 | 1.45 | -15.26 | **<0.001** |
| V/E | -0.15 | 1.49 | -0.10 | 0.985 | -2.23 | 1.40 | -1.59 | 0.128 | 1.41 | 1.44 | 0.98 | 0.328 | -7.71 | 1.44 | -5.35 | **<0.001** |
| E/V | -0.03 | 1.49 | -0.02 | 0.985 | 8.36 | 1.40 | 5.96 | **<0.001** | 7.92 | 1.43 | 5.53 | **<0.001** | 2.63 | 1.45 | 1.81 | 0.080 |
| Medium Force:<br>36.25N Elastic, -20N Viscous | | | | | Medium Force:<br>36.25N Elastic, -20N Viscous | | | | Medium Force:<br>36.25N Elastic, -20N Viscous | | | | Medium Force:<br>36.25N Elastic, -20N Viscous | | | |
| V/V | 1.90 | 1.48 | 1.28 | 0.228 | 3.21 | 1.40 | 2.29 | **0.029** | 6.82 | 1.42 | 4.79 | **<0.001** | -4.25 | 1.44 | -2.94 | **0.004** |
| E/E | 0.14 | 1.49 | 0.09 | 0.925 | 1.37 | 1.40 | 0.98 | 0.328 | 6.15 | 1.43 | 4.29 | **<0.001** | -5.32 | 1.45 | -3.67 | **<0.001** |
| NL/E | 23.33 | 1.53 | 15.21 | **<0.001** | 21.94 | 1.40 | 15.63 | **<0.001** | 22.26 | 1.44 | 15.50 | **<0.001** | 18.03 | 1.49 | 12.12 | **<0.001** |
| E/NL | -26.31 | 1.53 | -17.15 | **<0.001** | -17.98 | 1.40 | -12.81 | **<0.001** | -18.20 | 1.47 | -12.42 | **<0.001** | -23.88 | 1.45 | -16.43 | **<0.001** |
| NL/V | 28.74 | 1.52 | 18.88 | **<0.001** | 28.50 | 1.41 | 20.26 | **<0.001** | 27.59 | 1.43 | 19.26 | **<0.001** | 26.66 | 1.47 | 18.12 | **<0.001** |
| V/NL | -26.06 | 1.51 | -17.23 | **<0.001** | -22.14 | 1.40 | -15.76 | **<0.001** | -22.18 | 1.45 | -15.27 | **<0.001** | -27.78 | 1.45 | -19.19 | **<0.001** |
| V/E | -2.77 | 1.49 | -1.86 | 0.084 | -2.93 | 1.40 | -2.09 | **0.042** | -1.58 | 1.43 | -1.11 | 0.268 | -12.62 | 1.45 | -8.70 | **<0.001** |
| E/V | 2.88 | 1.48 | 1.94 | 0.083 | 9.76 | 1.41 | 6.94 | **<0.001** | 10.53 | 1.43 | 7.36 | **<0.001** | 1.92 | 1.46 | 1.31 | 0.190 |

#### Supplementary Table 16. Left/Right Pairwise Comparison, Time to Peak Velocity (TPV)

**T16. Post-Hoc Comparisons of Time to Peak Velocity (TPV) between hands.** Tukey-adjusted pairwise comparisons of left- and right-hand TPV within each load condition, across all reach types and force levels.

| Time to Peak Velocity (TPV) | | | | | | | | | | | | | | | | |
|---|---|---|---|---|---|---|---|---|---|---|---|---|---|---|---|---|
| | **Symmetric Long Reach<br><br>Low Force:<br>30N Elastic, -15N Viscous** | | | | **Symmetric Short Reach<br><br>Low Force:<br>30N Elastic, -15N Viscous** | | | | **Asymmetric Left Long. Right Short Reach<br><br>Low Force:<br>.30N Elastic, -15N Viscous** | | | | **Asymmetric Left Short, Right Long Reach<br><br>Low Force:<br>30N Elastic, -15N Viscous** | | | |
| **Loads Applied: Left/Right Arm** | ***β*** | **SE** | **Z-Ratio** | **P Value** | ***β*** | **SE** | **Z-Ratio** | **P Value** | ***β*** | **SE** | **Z-Ratio** | **P Value** | **Estimate** | **SE** | **Z-Ratio** | **P Value** |
| NL/NL | -4.85 | 13.13 | -0.37 | 0.712 | 3.06 | 11.34 | 0.27 | 0.787 | 28.01 | 15.52 | 1.80 | 0.071 | -27.41 | 14.67 | -1.87 | 0.062 |
| V/V | 2.60 | 10.15 | 0.26 | 0.798 | 4.57 | 10.15 | 0.45 | 0.652 | 109.68 | 10.10 | 10.86 | **<0.001** | -117.51 | 10.19 | -11.53 | **<0.001** |
| E/E | 2.03 | 10.25 | 0.20 | 0.843 | 1.53 | 10.16 | 0.15 | 0.880 | 171.65 | 10.16 | 16.90 | **<0.001** | -144.43 | 10.13 | -14.26 | **<0.001** |
| NL/E | 83.34 | 10.96 | 7.60 | **<0.001** | 0.23 | 10.09 | 0.02 | 0.982 | 202.28 | 10.16 | 19.92 | **<0.001** | -82.92 | 10.77 | -7.70 | **<0.001** |
| E/NL | -116.87 | 10.96 | -10.66 | **<0.001** | 18.05 | 10.16 | 1.78 | 0.076 | 115.73 | 10.59 | 10.93 | **<0.001** | -185.70 | 10.10 | -18.38 | **<0.001** |
| NL/V | 200.57 | 10.69 | 18.77 | **<0.001** | 146.90 | 10.08 | 14.58 | **<0.001** | 292.42 | 10.16 | 28.79 | **<0.001** | 75.44 | 10.71 | 7.05 | **<0.001** |
| V/NL | -186.83 | 10.57 | -17.67 | **<0.001** | -145.85 | 10.07 | -14.49 | **<0.001** | -66.00 | 10.70 | -6.17 | **<0.001** | -275.28 | 10.16 | -27.08 | **<0.001** |
| V/E | -93.67 | 10.22 | -9.17 | **<0.001** | -166.33 | 10.17 | -16.36 | **<0.001** | 37.78 | 10.22 | 3.70 | **<0.001** | -256.97 | 10.21 | -25.17 | **<0.001** |
| E/V | 109.78 | 10.26 | 10.70 | **<0.001** | 172.07 | 10.11 | 17.02 | **<0.001** | 293.47 | 10.18 | 28.82 | **<0.001** | -25.09 | 10.16 | -2.47 | **0.014** |
| | **Medium Force:<br>36.25N Elastic, -20N Viscous** | | | | **Medium Force:<br>36.25N Elastic, -20N Viscous** | | | | **Medium Force:<br>36.25N Elastic, -20N Viscous** | | | | **Medium Force:<br>36.25N Elastic, -20N Viscous** | | | |
| NL/NL | -4.85 | 13.13 | -0.37 | 0.712 | 3.06 | 11.34 | 0.27 | 0.787 | 28.01 | 15.52 | 1.80 | 0.071 | -27.41 | 14.67 | -1.87 | 0.062 |
| V/V | 9.31 | 10.13 | 0.92 | 0.358 | -0.66 | 10.12 | -0.06 | 0.948 | 152.69 | 10.14 | 15.06 | **<0.001** | -137.09 | 10.18 | -13.46 | **<0.001** |
| E/E | 7.72 | 10.13 | 0.76 | 0.446 | 0.92 | 10.13 | 0.09 | 0.928 | 179.06 | 10.17 | 17.60 | **<0.001** | -165.51 | 10.20 | -16.23 | **<0.001** |
| NL/E | 125.84 | 10.94 | 11.50 | **<0.001** | -5.85 | 10.09 | -0.58 | 0.562 | 205.69 | 10.12 | 20.33 | **<0.001** | -57.83 | 10.80 | -5.35 | **<0.001** |
| E/NL | -104.15 | 10.79 | -9.65 | **<0.001** | 14.44 | 10.10 | 1.43 | 0.153 | 110.43 | 10.77 | 10.25 | **<0.001** | -173.91 | 10.22 | -17.02 | **<0.001** |
| NL/V | 244.25 | 10.77 | 22.68 | **<0.001** | 182.24 | 10.16 | 17.94 | **<0.001** | 360.20 | 10.18 | 35.40 | **<0.001** | 96.73 | 10.62 | 9.11 | **<0.001** |
| V/NL | -226.12 | 10.66 | -21.20 | **<0.001** | -172.23 | 10.10 | -17.05 | **<0.001** | -86.70 | 10.70 | -8.11 | **<0.001** | -332.64 | 10.15 | -32.78 | **<0.001** |
| V/E | -119.24 | 10.18 | -11.71 | **<0.001** | -168.23 | 10.12 | -16.62 | **<0.001** | 6.50 | 10.16 | 0.64 | 0.523 | -289.10 | 10.20 | -28.34 | **<0.001** |
| E/V | 121.64 | 10.17 | 11.96 | **<0.001** | 182.61 | 10.06 | 18.15 | **<0.001** | 332.53 | 10.13 | 32.83 | **<0.001** | -6.09 | 10.15 | -0.60 | 0.548 |

#### Supplementary Table 17. Between-Hand Time to Peak Velocity (TPV) Difference Across Loads (Relative to NL/NL)

**T17. Post-Hoc Comparisons of Time to Peak Velocity (TPV) Differences between Hands by Load Condition and Reach Type.** Pairwise comparisons assess whether hand differences in TPV significantly vary across load conditions, using the No Load/No Load condition as the baseline.

| Time to Peak Velocity (TPV) | | | | | | | | | | | | | | | | |
|---|---|---|---|---|---|---|---|---|---|---|---|---|---|---|---|---|
| Symmetric Long Reach<br><br>Low Force:<br>30N Elastic, -15N Viscous | | | | | Symmetric Short Reach<br><br>Low Force:<br>30N Elastic, -15N Viscous | | | | Asymmetric Left Long. Right Short Reach<br><br>Low Force:<br>.30N Elastic, -15N Viscous | | | | Asymmetric Left Short, Right Long Reach<br><br>Low Force:<br>30N Elastic, -15N Viscous | | | |
| Loads Applied: Left/Right Arm | β | SE | Z-Ratio | P Value | β | SE | Z-Ratio | P Value | β | SE | Z-Ratio | P Value | Estimate | SE | Z-Ratio | P Value |
| V/V | -7.38 | 13.70 | -0.54 | 0.674 | 4.58 | 12.55 | 0.36 | 0.817 | -68.79 | 15.28 | -4.50 | **<0.001** | 78.62 | 14.74 | 5.33 | **<0.001** |
| E/E | -5.69 | 13.75 | -0.41 | 0.679 | 1.85 | 12.60 | 0.15 | 0.883 | -122.18 | 15.30 | -7.98 | **<0.001** | 103.17 | 14.73 | 7.00 | **<0.001** |
| NL/E | -83.86 | 14.14 | -5.93 | **<0.001** | 6.88 | 12.54 | 0.55 | 0.778 | -158.52 | 15.31 | -10.35 | **<0.001** | 47.24 | 15.05 | 3.14 | **0.002** |
| E/NL | 106.90 | 14.13 | 7.57 | **<0.001** | -15.94 | 12.61 | -1.26 | 0.330 | -73.85 | 15.50 | -4.76 | **<0.001** | 144.71 | 14.71 | 9.84 | **<0.001** |
| NL/V | -206.50 | 14.02 | -14.73 | **<0.001** | -140.86 | 12.49 | -11.28 | **<0.001** | -247.48 | 15.30 | -16.18 | **<0.001** | -114.43 | 15.00 | -7.63 | **<0.001** |
| V/NL | 183.92 | 13.94 | 13.20 | **<0.001** | 152.04 | 12.48 | 12.18 | **<0.001** | 108.84 | 15.58 | 6.98 | **<0.001** | 235.44 | 14.72 | 15.99 | **<0.001** |
| V/E | 89.85 | 13.75 | 6.53 | **<0.001** | 169.40 | 12.60 | 13.44 | **<0.001** | 4.45 | 15.34 | 0.29 | 0.772 | 217.43 | 14.77 | 14.72 | **<0.001** |
| E/V | -115.54 | 13.80 | -8.37 | **<0.001** | -164.62 | 12.54 | -13.13 | **<0.001** | -252.55 | 15.35 | -16.46 | **<0.001** | -17.94 | 14.72 | -1.22 | 0.223 |
| Medium Force:<br>36.25N Elastic, -20N Viscous | | | | | Medium Force:<br>36.25N Elastic, -20N Viscous | | | | Medium Force:<br>36.25N Elastic, -20N Viscous | | | | Medium Force:<br>36.25N Elastic, -20N Viscous | | | |
| V/V | -13.93 | 13.70 | -1.02 | 0.354 | 8.52 | 12.53 | 0.68 | 0.567 | -104.76 | 15.31 | -6.84 | **<0.001** | 98.45 | 14.74 | 6.68 | **<0.001** |
| E/E | -11.72 | 13.71 | -0.85 | 0.393 | 6.12 | 12.56 | 0.49 | 0.626 | -129.62 | 15.35 | -8.45 | **<0.001** | 123.94 | 14.76 | 8.40 | **<0.001** |
| NL/E | -126.32 | 14.12 | -8.95 | **<0.001** | 12.37 | 12.52 | 0.99 | 0.517 | -164.07 | 15.27 | -10.74 | **<0.001** | 23.88 | 15.07 | 1.58 | 0.113 |
| E/NL | 92.61 | 14.08 | 6.58 | **<0.001** | -8.74 | 12.55 | -0.70 | 0.567 | -65.69 | 15.62 | -4.21 | **<0.001** | 129.93 | 14.77 | 8.80 | **<0.001** |
| NL/V | -248.28 | 14.04 | -17.69 | **<0.001** | -176.80 | 12.57 | -14.06 | **<0.001** | -313.04 | 15.36 | -20.38 | **<0.001** | -136.83 | 14.97 | -9.14 | **<0.001** |
| V/NL | 216.43 | 14.02 | 15.44 | **<0.001** | 176.60 | 12.56 | 14.06 | **<0.001** | 126.53 | 15.59 | 8.12 | **<0.001** | 288.31 | 14.75 | 19.55 | **<0.001** |
| V/E | 118.01 | 13.74 | 8.59 | **<0.001** | 174.91 | 12.57 | 13.91 | **<0.001** | 34.13 | 15.31 | 2.23 | **0.026** | 251.97 | 14.76 | 17.07 | **<0.001** |
| E/V | -135.44 | 13.74 | -9.86 | **<0.001** | -172.23 | 12.50 | -13.78 | **<0.001** | -285.21 | 15.31 | -18.62 | **<0.001** | -34.68 | 14.70 | -2.36 | **0.021** |

#### Supplementary Table 18. Left-Right Hand Difference LME Model, Peak Acceleration & Time at Peak Acceleration

**T18. Results from Linear Mixed-Effects Models Examining Right-Left Hand Differences in (a) Peak Acceleration (PA) and (b) Time to Peak Acceleration (TPA).** Fixed effects estimates (β), standard errors (SE), and p-values are reported for each model, which included random intercepts for participant and trial number. Fixed effects included Load Condition, Reach Configuration, and Force Magnitude. Significant effects ($p < 0.05$) are shown in bold.

| | **Linear Mixed-Effects Model Results: Right–Left Hand Differences in Peak Acceleration (PA) and Time to Peak Acceleration (TPA)** | | | | | |
|---|---|---|---|---|---|---|
| | **(a) PA** | | | **(b) TPA** | | |
| **Fixed effects L=left arm/R=right arm)** | ***β*** | **SE** | ***p(χ²)*** | ***β*** | **SE** | ***p(χ²)*** |
| Intercept (No Load (L)/No Load (R)) | -29.02 | 6.92 | **<0.001** | -4.79 | 5.08 | 0.345 |
| Elastic (L)/Elastic (R) | -2.59 | 3.17 | 0.414 | -2.05 | 3.30 | 0.534 |
| Viscous (L)/Viscous (R) | 10.05 | 3.17 | 0.002 | 2.74 | 3.24 | 0.398 |
| No Load (L)/Elastic (R) | 64.10 | 3.17 | **<0.001** | 4.46 | 3.27 | 0.173 |
| Elastic (L)/No Load (R) | -63.51 | 3.17 | **<0.001** | -8.52 | 3.28 | **0.009** |
| No Load (L)/Viscous (R) | 206.14 | 3.18 | **<0.001** | -77.35 | 3.25 | **<0.001** |
| Viscous (L)/No Load (R) | -195.25 | 3.17 | **<0.001** | 91.59 | 3.25 | **<0.001** |
| Viscous (L)/Elastic (R) | -132.77 | 3.17 | **<0.001** | 74.19 | 3.28 | **<0.001** |
| Elastic (L)/Viscous (R) | 141.60 | 3.18 | **<0.001** | -85.95 | 3.28 | **<0.001** |
| Reach Configuration [Symmetric Short Reach] | 7.86 | 2.11 | **<0.001** | -1.00 | 2.19 | 0.648 |
| Reach Configuration [Asymmetric Left-Long/Right-Short Reach] | -79.72 | 2.11 | **<0.001** | -50.32 | 2.20 | **<0.001** |
| Reach Configuration [Asymmetric Left-Short/Right-Long Reach] | 93.49 | 2.11 | **<0.001** | 38.80 | 2.20 | **<0.001** |
| Force Magnitude [Medium Force: 36.25N Elastic, -20N Viscous] | 0.69 | 1.49 | 0.644 | -0.79 | 1.55 | 0.609 |
| **Random effects** | **Groups** | | **SD** | **Groups** | | **SD** |
| | Trial Number | Intercept | 7.71 | Trial Number | Intercept | |
| | Participant | Intercept | 34.82 | Participant | Intercept | |
| | Residual | | 104.80 | Residual | | |
| | Observations: | 19848 | | Observations: | | |
| **Full Model:** Right-Left Difference in PA or TPA ~ Load Condition + Reach Configuration + Force Magnitude + (1 \| Trial Number) + (1 \| Participant) | | | | | | |

#### Supplementary Table 19. Left/Right Pairwise Comparison, Peak Acceleration Time (PA)

**T19. Post-Hoc Comparisons of Peak Acceleration (PA) between hands.** Tukey-adjusted pairwise comparisons of left- and right-hand PA within each load condition, across all reach types and force levels.

| Peak Acceleration (PA) | | | | | | | | | | | | | | | | |
|---|---|---|---|---|---|---|---|---|---|---|---|---|---|---|---|---|
| Symmetric Long Reach<br><br>Low Force:<br>30N Elastic, -15N Viscous | | | | | Symmetric Short Reach<br><br>Low Force:<br>30N Elastic, -15N Viscous | | | | Asymmetric Left Long. Right Short Reach<br><br>Low Force:<br>.30N Elastic, -15N Viscous | | | | Asymmetric Left Short, Right Long Reach<br><br>Low Force:<br>30N Elastic, -15N Viscous | | | |
| **Loads Applied: Left/Right Arm** | ***β*** | **SE** | **Z-Ratio** | **P Value** | ***β*** | **SE** | **Z-Ratio** | **P Value** | ***β*** | **SE** | **Z-Ratio** | **P Value** | **Estimate** | **SE** | **Z-Ratio** | **P Value** |
| NL/NL | 28.67 | 7.33 | 3.91 | **<0.001** | 26.31 | 7.33 | 3.59 | **<0.001** | 132.15 | 7.30 | 18.09 | **<0.001** | -89.64 | 7.32 | -12.24 | **<0.001** |
| V/V | 16.40 | 7.28 | 2.25 | **0.024** | 9.68 | 7.36 | 1.31 | 0.189 | 91.14 | 7.33 | 12.44 | **<0.001** | -56.05 | 7.29 | -7.69 | **<0.001** |
| E/E | 34.35 | 7.28 | 4.72 | **<0.001** | 19.34 | 7.32 | 2.64 | **0.008** | 119.57 | 7.29 | 16.40 | **<0.001** | -68.83 | 7.30 | -9.42 | **<0.001** |
| NL/E | -36.90 | 7.30 | -5.05 | **<0.001** | -28.44 | 7.30 | -3.90 | **<0.001** | 58.68 | 7.30 | 8.04 | **<0.001** | -138.78 | 7.32 | -18.97 | **<0.001** |
| E/NL | 110.33 | 7.26 | 15.20 | **<0.001** | 73.02 | 7.30 | 10.01 | **<0.001** | 177.15 | 7.28 | 24.32 | **<0.001** | -17.61 | 7.29 | -2.42 | **0.016** |
| NL/V | -182.58 | 7.28 | -25.07 | **<0.001** | -158.80 | 7.30 | -21.75 | **<0.001** | -77.17 | 7.31 | -10.56 | **<0.001** | -260.38 | 7.30 | -35.68 | **<0.001** |
| V/NL | 238.81 | 7.30 | 32.70 | **<0.001** | 206.09 | 7.32 | 28.17 | **<0.001** | 294.41 | 7.30 | 40.31 | **<0.001** | 110.16 | 7.26 | 15.18 | **<0.001** |
| V/E | 165.53 | 7.30 | 22.68 | **<0.001** | 151.78 | 7.28 | 20.84 | **<0.001** | 225.09 | 7.33 | 30.71 | **<0.001** | 67.55 | 7.27 | 9.29 | **<0.001** |
| E/V | -109.90 | 7.30 | -15.05 | **<0.001** | -111.91 | 7.29 | -15.35 | **<0.001** | -37.99 | 7.29 | -5.21 | **<0.001** | -206.36 | 7.30 | -28.26 | **<0.001** |
| **Medium Force:<br>36.25N Elastic, -20N Viscous** | | | | | **Medium Force:<br>36.25N Elastic, -20N Viscous** | | | | **Medium Force:<br>36.25N Elastic, -20N Viscous** | | | | **Medium Force:<br>36.25N Elastic, -20N Viscous** | | | |
| NL/NL | 28.67 | 7.33 | 3.91 | **<0.001** | 26.31 | 7.33 | 3.59 | **<0.001** | 132.15 | 7.30 | 18.09 | **<0.001** | -89.64 | 7.32 | -12.24 | **<0.001** |
| V/V | 9.97 | 7.28 | 1.37 | 0.171 | 15.09 | 7.31 | 2.07 | **0.039** | 81.04 | 7.33 | 11.06 | **<0.001** | -51.31 | 7.28 | -7.05 | **<0.001** |
| E/E | 36.77 | 7.28 | 5.05 | **<0.001** | 26.82 | 7.31 | 3.67 | **<0.001** | 107.74 | 7.30 | 14.77 | **<0.001** | -58.83 | 7.30 | -8.05 | **<0.001** |
| NL/E | -50.26 | 7.28 | -6.91 | **<0.001** | -38.56 | 7.31 | -5.28 | **<0.001** | 53.90 | 7.26 | 7.43 | **<0.001** | -136.76 | 7.30 | -18.74 | **<0.001** |
| E/NL | 105.89 | 7.30 | 14.51 | **<0.001** | 76.55 | 7.32 | 10.46 | **<0.001** | 193.79 | 7.33 | 26.44 | **<0.001** | -9.58 | 7.26 | -1.32 | 0.187 |
| NL/V | -218.23 | 7.32 | -29.83 | **<0.001** | -184.45 | 7.34 | -25.12 | **<0.001** | -109.10 | 7.30 | -14.94 | **<0.001** | -288.66 | 7.28 | -39.63 | **<0.001** |
| V/NL | 261.24 | 7.28 | 35.89 | **<0.001** | 211.84 | 7.28 | 29.09 | **<0.001** | 316.96 | 7.27 | 43.59 | **<0.001** | 136.39 | 7.25 | 18.80 | **<0.001** |
| V/E | 178.64 | 7.30 | 24.48 | **<0.001** | 150.26 | 7.30 | 20.59 | **<0.001** | 245.98 | 7.30 | 33.71 | **<0.001** | 90.07 | 7.30 | 12.34 | **<0.001** |
| E/V | -133.33 | 7.32 | -18.23 | **<0.001** | -127.17 | 7.32 | -17.38 | **<0.001** | -43.93 | 7.32 | -6.00 | **<0.001** | -192.12 | 7.33 | -26.21 | **<0.001** |

#### Supplementary Table 20. Between-Hand Peak Acceleration (PA) Difference Across Loads (Relative to NL/NL)

**T20. Post-Hoc Comparisons of Peak Acceleration (PA) Differences between Hands by Load Condition and Reach Type.** Pairwise comparisons assess whether hand differences in PA significantly vary across load conditions, using the No Load/No Load condition as the baseline.

| **Peak Acceleration (PA)** | | | | | | | | | | | | | | | | |
|---|---|---|---|---|---|---|---|---|---|---|---|---|---|---|---|---|
| **Symmetric Long Reach<br><br>Low Force:<br>30N Elastic, -15N Viscous** | | | | | **Symmetric Short Reach<br><br>Low Force:<br>30N Elastic, -15N Viscous** | | | | **Asymmetric Left Long. Right Short Reach<br><br>Low Force:<br>.30N Elastic, -15N Viscous** | | | | **Asymmetric Left Short, Right Long Reach<br><br>Low Force:<br>30N Elastic, -15N Viscous** | | | |
| **Loads Applied: Left/Right Arm** | ***β*** | **SE** | **Z-Ratio** | **P Value** | ***β*** | **SE** | **Z-Ratio** | **P Value** | ***β*** | **SE** | **Z-Ratio** | **P Value** | **Estimate** | **SE** | **Z-Ratio** | **P Value** |
| V/V | 11.83 | 8.86 | 1.33 | 0.208 | 16.52 | 8.91 | 1.86 | 0.073 | 36.68 | 8.91 | 4.12 | **<0.001** | -29.22 | 8.93 | -3.27 | **0.001** |
| E/E | -7.56 | 8.88 | -0.85 | 0.395 | 8.21 | 8.90 | 0.92 | 0.356 | 10.05 | 8.88 | 1.13 | 0.258 | -18.00 | 8.94 | -2.01 | **0.044** |
| NL/E | 66.71 | 8.91 | 7.49 | **<0.001** | 54.29 | 8.88 | 6.11 | **<0.001** | 69.66 | 8.89 | 7.84 | **<0.001** | 52.48 | 8.92 | 5.88 | **<0.001** |
| E/NL | -81.52 | 8.87 | -9.19 | **<0.001** | -46.32 | 8.90 | -5.21 | **<0.001** | -49.46 | 8.87 | -5.57 | **<0.001** | -68.96 | 8.91 | -7.74 | **<0.001** |
| NL/V | 206.86 | 8.88 | 23.29 | **<0.001** | 184.92 | 8.89 | 20.80 | **<0.001** | 202.90 | 8.90 | 22.80 | **<0.001** | 171.63 | 8.91 | 19.26 | **<0.001** |
| V/NL | -208.32 | 8.90 | -23.39 | **<0.001** | -177.16 | 8.94 | -19.82 | **<0.001** | -162.80 | 8.89 | -18.32 | **<0.001** | -193.86 | 8.89 | -21.81 | **<0.001** |
| V/E | -136.42 | 8.90 | -15.33 | **<0.001** | -124.08 | 8.87 | -13.98 | **<0.001** | -91.73 | 8.93 | -10.27 | **<0.001** | -150.48 | 8.89 | -16.94 | **<0.001** |
| E/V | 133.13 | 8.91 | 14.94 | **<0.001** | 136.96 | 8.87 | 15.45 | **<0.001** | 163.31 | 8.88 | 18.39 | **<0.001** | 117.13 | 8.92 | 13.13 | **<0.001** |
| **Medium Force:<br>36.25N Elastic, -20N Viscous** | | | | | **Medium Force:<br>36.25N Elastic, -20N Viscous** | | | | **Medium Force:<br>36.25N Elastic, -20N Viscous** | | | | **Medium Force:<br>36.25N Elastic, -20N Viscous** | | | |
| V/V | 17.05 | 8.86 | 1.92 | 0.062 | 10.66 | 8.87 | 1.20 | 0.262 | 47.39 | 8.91 | 5.32 | **<0.001** | -31.04 | 8.89 | -3.49 | **<0.001** |
| E/E | -8.84 | 8.89 | -0.99 | 0.320 | -0.18 | 8.88 | -0.02 | 0.984 | 20.02 | 8.88 | 2.25 | **0.024** | -24.85 | 8.91 | -2.79 | **0.005** |
| NL/E | 75.90 | 8.89 | 8.54 | **<0.001** | 63.34 | 8.89 | 7.13 | **<0.001** | 77.06 | 8.87 | 8.68 | **<0.001** | 52.91 | 8.89 | 5.95 | **<0.001** |
| E/NL | -76.74 | 8.90 | -8.62 | **<0.001** | -50.04 | 8.90 | -5.62 | **<0.001** | -60.56 | 8.91 | -6.79 | **<0.001** | -74.88 | 8.88 | -8.44 | **<0.001** |
| NL/V | 241.54 | 8.91 | 27.09 | **<0.001** | 208.16 | 8.89 | 23.41 | **<0.001** | 233.26 | 8.90 | 26.19 | **<0.001** | 199.51 | 8.91 | 22.40 | **<0.001** |
| V/NL | -231.76 | 8.90 | -26.03 | **<0.001** | -183.68 | 8.85 | -20.75 | **<0.001** | -184.88 | 8.86 | -20.86 | **<0.001** | -219.95 | 8.90 | -24.72 | **<0.001** |
| V/E | -148.59 | 8.88 | -16.73 | **<0.001** | -121.93 | 8.89 | -13.72 | **<0.001** | -117.64 | 8.88 | -13.24 | **<0.001** | -171.42 | 8.91 | -19.23 | **<0.001** |
| E/V | 157.83 | 8.90 | 17.73 | **<0.001** | 151.09 | 8.93 | 16.92 | **<0.001** | 169.54 | 8.91 | 19.02 | **<0.001** | 103.01 | 8.96 | 11.49 | **<0.001** |

#### Supplementary Table 21. Left/Right Pairwise Comparison, Time at Peak Acceleration (TPA)

**T21. Post-Hoc Comparisons of Time at Peak Acceleration (TPA) between hands.** Tukey-adjusted pairwise comparisons of left- and right-hand TPA within each load condition, across all reach types and force levels.

| Time at Peak Acceleration (TPA) | | | | | | | | | | | | | | | | |
|---|---|---|---|---|---|---|---|---|---|---|---|---|---|---|---|---|
| **Symmetric Long Reach<br><br>Low Force:<br>30N Elastic, -15N Viscous** | | | | | **Symmetric Short Reach<br><br>Low Force:<br>30N Elastic, -15N Viscous** | | | | **Asymmetric Left Long. Right Short Reach<br><br>Low Force:<br>.30N Elastic, -15N Viscous** | | | | **Asymmetric Left Short, Right Long Reach<br><br>Low Force:<br>30N Elastic, -15N Viscous** | | | |
| **Loads Applied: Left/Right Arm** | ***β*** | **SE** | **Z-Ratio** | **P Value** | ***β*** | **SE** | **Z-Ratio** | **P Value** | ***β*** | **SE** | **Z-Ratio** | **P Value** | **Estimate** | **SE** | **Z-Ratio** | **P Value** |
| NL/NL | 0.88 | 7.83 | 0.11 | 0.911 | 14.31 | 7.86 | 1.82 | 0.069 | 36.64 | 7.83 | 4.68 | **<0.001** | -17.93 | 7.87 | -2.28 | **0.023** |
| V/V | -14.33 | 7.83 | -1.83 | 0.067 | 6.56 | 7.77 | 0.84 | 0.399 | 93.56 | 7.79 | 12.01 | **<0.001** | -65.78 | 7.83 | -8.40 | **<0.001** |
| E/E | 25.29 | 7.97 | 3.17 | **0.002** | 6.41 | 7.93 | 0.81 | 0.419 | 41.98 | 7.97 | 5.27 | **<0.001** | -23.30 | 7.98 | -2.92 | **0.004** |
| NL/E | -5.73 | 7.92 | -0.72 | 0.470 | 5.38 | 7.88 | 0.68 | 0.495 | 47.29 | 7.86 | 6.02 | **<0.001** | -29.36 | 7.90 | -3.72 | **<0.001** |
| E/NL | 16.61 | 7.90 | 2.10 | **0.036** | 15.55 | 7.90 | 1.97 | **0.049** | 43.49 | 7.96 | 5.46 | **<0.001** | -4.48 | 7.88 | -0.57 | 0.570 |
| NL/V | 104.96 | 7.81 | 13.45 | **<0.001** | 51.87 | 7.83 | 6.63 | **<0.001** | 118.61 | 7.86 | 15.09 | **<0.001** | 29.03 | 7.82 | 3.71 | **<0.001** |
| V/NL | -112.44 | 7.87 | -14.29 | **<0.001** | -46.71 | 7.83 | -5.96 | **<0.001** | -16.39 | 7.79 | -2.10 | **0.035** | -127.73 | 7.85 | -16.26 | **<0.001** |
| V/E | -101.51 | 7.99 | -12.71 | **<0.001** | -52.78 | 7.84 | -6.73 | **<0.001** | -8.66 | 7.95 | -1.09 | 0.276 | -91.57 | 7.88 | -11.62 | **<0.001** |
| E/V | 114.61 | 7.91 | 14.49 | **<0.001** | 63.08 | 7.89 | 8.00 | **<0.001** | 139.76 | 7.93 | 17.62 | **<0.001** | 17.65 | 7.90 | 2.23 | **0.026** |
| **Medium Force:<br>36.25N Elastic, -20N Viscous** | | | | | **Medium Force:<br>36.25N Elastic, -20N Viscous** | | | | **Medium Force:<br>36.25N Elastic, -20N Viscous** | | | | **Medium Force:<br>36.25N Elastic, -20N Viscous** | | | |
| NL/NL | 0.88 | 7.83 | 0.11 | 0.911 | 14.31 | 7.86 | 1.82 | 0.069 | 36.64 | 7.83 | 4.68 | **<0.001** | -17.93 | 7.87 | -2.28 | **0.023** |
| V/V | 4.31 | 7.94 | 0.54 | 0.587 | -10.72 | 7.79 | -1.38 | 0.169 | 80.46 | 7.85 | 10.25 | **<0.001** | -51.25 | 7.82 | -6.55 | **<0.001** |
| E/E | 0.72 | 7.94 | 0.09 | 0.927 | 15.32 | 8.02 | 1.91 | 0.056 | 51.58 | 7.91 | 6.52 | **<0.001** | -37.02 | 8.00 | -4.63 | **<0.001** |
| NL/E | 19.70 | 7.87 | 2.50 | **0.012** | -4.65 | 7.87 | -0.59 | 0.555 | 43.58 | 7.90 | 5.52 | **<0.001** | -21.86 | 7.90 | -2.77 | **0.006** |
| E/NL | 13.97 | 7.90 | 1.77 | 0.077 | 17.09 | 7.94 | 2.15 | **0.031** | 42.27 | 7.97 | 5.30 | **<0.001** | -34.07 | 7.89 | -4.32 | **<0.001** |
| NL/V | 118.70 | 7.88 | 15.06 | **<0.001** | 77.65 | 7.82 | 9.93 | **<0.001** | 147.12 | 7.87 | 18.68 | **<0.001** | 48.41 | 7.84 | 6.18 | **<0.001** |
| V/NL | -128.28 | 7.88 | -16.28 | **<0.001** | -69.31 | 7.79 | -8.90 | **<0.001** | -40.76 | 7.85 | -5.20 | **<0.001** | -127.84 | 7.86 | -16.26 | **<0.001** |
| V/E | -98.28 | 7.92 | -12.40 | **<0.001** | -77.16 | 7.88 | -9.80 | **<0.001** | 1.32 | 7.85 | 0.17 | 0.866 | -104.63 | 7.90 | -13.25 | **<0.001** |
| E/V | 125.07 | 7.93 | 15.77 | **<0.001** | 89.09 | 7.91 | 11.27 | **<0.001** | 143.52 | 7.95 | 18.06 | **<0.001** | 51.56 | 7.86 | 6.56 | **<0.001** |

#### Supplementary Table 22. Between-Hand Time at Peak Acceleration (TPA) Difference Across Loads (Relative to NL/NL)

**T22. Post-Hoc Comparisons of Time at Peak Acceleration (TPA) Differences between Hands by Load Condition and Reach Type.** Pairwise comparisons assess whether hand differences in TPA significantly vary across load conditions, using the No Load/No Load condition as the baseline.

| **Time at Peak Acceleration (TPA)** | | | | | | | | | | | | | | | | |
|---|---|---|---|---|---|---|---|---|---|---|---|---|---|---|---|---|
| **Symmetric Long Reach<br><br>Low Force:<br>30N Elastic, -15N Viscous** | | | | | **Symmetric Short Reach<br><br>Low Force:<br>30N Elastic, -15N Viscous** | | | | **Asymmetric Left Long. Right Short Reach<br><br>Low Force:<br>.30N Elastic, -15N Viscous** | | | | **Asymmetric Left Short, Right Long Reach<br><br>Low Force:<br>30N Elastic, -15N Viscous** | | | |
| **Loads Applied: Left/Right Arm** | ***β*** | **SE** | **Z-Ratio** | **P Value** | ***β*** | **SE** | **Z-Ratio** | **P Value** | ***β*** | **SE** | **Z-Ratio** | **P Value** | **Estimate** | **SE** | **Z-Ratio** | **P Value** |
| V/V | 14.13 | 9.04 | 1.56 | 0.135 | 8.84 | 8.99 | 0.98 | 0.372 | -58.88 | 9.02 | -6.53 | **<0.001** | 46.56 | 9.06 | 5.14 | **<0.001** |
| E/E | -20.57 | 9.19 | -2.24 | **0.040** | 9.18 | 9.16 | 1.00 | 0.372 | -10.20 | 9.20 | -1.11 | 0.306 | 8.23 | 9.27 | 0.89 | 0.375 |
| NL/E | 9.39 | 9.11 | 1.03 | 0.303 | 11.48 | 9.14 | 1.26 | 0.334 | -7.92 | 9.11 | -0.87 | 0.385 | 15.03 | 9.18 | 1.64 | 0.135 |
| E/NL | -19.74 | 9.13 | -2.16 | **0.041** | -6.32 | 9.15 | -0.69 | 0.490 | -13.00 | 9.22 | -1.41 | 0.211 | -14.28 | 9.14 | -1.56 | 0.135 |
| NL/V | -100.03 | 9.01 | -11.10 | **<0.001** | -39.67 | 9.04 | -4.39 | **<0.001** | -81.40 | 9.07 | -8.97 | **<0.001** | -48.82 | 9.05 | -5.40 | **<0.001** |
| V/NL | 114.43 | 9.05 | 12.65 | **<0.001** | 63.18 | 9.04 | 6.99 | **<0.001** | 50.19 | 9.01 | 5.57 | **<0.001** | 106.96 | 9.12 | 11.73 | **<0.001** |
| V/E | 103.05 | 9.23 | 11.16 | **<0.001** | 65.82 | 9.09 | 7.24 | **<0.001** | 42.64 | 9.17 | 4.65 | **<0.001** | 76.76 | 9.16 | 8.38 | **<0.001** |
| E/V | -115.68 | 9.16 | -12.63 | **<0.001** | -48.66 | 9.12 | -5.34 | **<0.001** | -105.89 | 9.17 | -11.54 | **<0.001** | -39.39 | 9.19 | -4.29 | **<0.001** |
| **Medium Force:<br>36.25N Elastic, -20N Viscous** | | | | | **Medium Force:<br>36.25N Elastic, -20N Viscous** | | | | **Medium Force:<br>36.25N Elastic, -20N Viscous** | | | | **Medium Force:<br>36.25N Elastic, -20N Viscous** | | | |
| V/V | -1.00 | 9.10 | -0.11 | 0.916 | 25.42 | 9.00 | 2.83 | **0.008** | -46.68 | 9.08 | -5.14 | **<0.001** | 33.46 | 9.05 | 3.70 | **<0.001** |
| E/E | 0.97 | 9.16 | 0.11 | 0.916 | -0.68 | 9.27 | -0.07 | 0.942 | -16.57 | 9.14 | -1.81 | 0.093 | 13.18 | 9.27 | 1.42 | 0.177 |
| NL/E | -13.46 | 9.09 | -1.48 | 0.202 | 19.74 | 9.10 | 2.17 | **0.040** | -4.08 | 9.15 | -0.45 | 0.656 | 6.02 | 9.18 | 0.66 | 0.512 |
| E/NL | -13.07 | 9.12 | -1.43 | 0.202 | -8.37 | 9.18 | -0.91 | 0.413 | -6.15 | 9.19 | -0.67 | 0.576 | 14.40 | 9.14 | 1.57 | 0.154 |
| NL/V | -111.99 | 9.10 | -12.30 | **<0.001** | -64.20 | 9.03 | -7.11 | **<0.001** | -109.55 | 9.10 | -12.03 | **<0.001** | -63.77 | 9.07 | -7.03 | **<0.001** |
| V/NL | 132.47 | 9.09 | 14.58 | **<0.001** | 82.44 | 9.01 | 9.15 | **<0.001** | 74.52 | 9.08 | 8.21 | **<0.001** | 111.15 | 9.11 | 12.20 | **<0.001** |
| V/E | 95.88 | 9.16 | 10.47 | **<0.001** | 90.79 | 9.10 | 9.97 | **<0.001** | 36.51 | 9.10 | 4.01 | **<0.001** | 84.62 | 9.18 | 9.22 | **<0.001** |
| E/V | -123.70 | 9.16 | -13.51 | **<0.001** | -72.90 | 9.14 | -7.98 | **<0.001** | -106.57 | 9.18 | -11.61 | **<0.001** | -74.99 | 9.14 | -8.21 | **<0.001** |

#### Supplementary Table 23. Left-Right Hand Difference LME Model, Peak Deceleration & Time at Peak Deceleration

**T23. Results from Linear Mixed-Effects Models Examining Right-Left Hand Differences in (a) Peak Deceleration (PD) and (b) Time to Peak Deceleration (TPD).** Fixed effects estimates (β), standard errors (SE), and p-values are reported for each model, which included random intercepts for participant and trial number. Fixed effects included Load Condition, Reach Configuration, and Force Magnitude. Significant effects ($p < 0.05$) are shown in bold.

| | **Linear Mixed-Effects Model Results: Right–Left Hand Differences in Peak Deceleration (PD) and Time to Peak Deceleration (TPD)** | | | | | |
|---|---|---|---|---|---|---|
| | **(a) PD** | | | **(b) TPD** | | |
| **Fixed effects L=left arm/R=right arm)** | ***β*** | **SE** | ***p(χ²)*** | ***β*** | **SE** | ***p(χ²)*** |
| Intercept (No Load (L)/No Load (R)) | -49.03 | 8.20 | **<0.001** | 1.19 | 9.67 | 0.902 |
| Elastic (L)/Elastic (R) | 5.29 | 5.27 | 0.316 | -0.14 | 6.13 | 0.982 |
| Viscous (L)/Viscous (R) | 32.57 | 5.35 | **<0.001** | -2.33 | 6.28 | 0.711 |
| No Load (L)/Elastic (R) | 63.60 | 5.28 | **<0.001** | -87.07 | 6.20 | **<0.001** |
| Elastic (L)/No Load (R) | -44.68 | 5.28 | **<0.001** | 88.75 | 6.20 | **<0.001** |
| No Load (L)/Viscous (R) | 227.76 | 5.33 | **<0.001** | -234.76 | 6.28 | **<0.001** |
| Viscous (L)/No Load (R) | -195.77 | 5.32 | **<0.001** | 227.19 | 6.26 | **<0.001** |
| Viscous (L)/Elastic (R) | -156.70 | 5.31 | **<0.001** | 140.60 | 6.24 | **<0.001** |
| Elastic (L)/Viscous (R) | 191.57 | 5.33 | **<0.001** | -150.04 | 6.24 | **<0.001** |
| Reach Configuration [Symmetric Short Reach] | 9.19 | 3.55 | **0.010** | 0.32 | 4.13 | 0.938 |
| Reach Configuration [Asymmetric Left-Long/Right-Short Reach] | -85.97 | 3.56 | **<0.001** | -177.72 | 4.16 | **<0.001** |
| Reach Configuration [Asymmetric Left-Short/Right-Long Reach] | 99.43 | 3.57 | **<0.001** | 179.46 | 4.16 | **<0.001** |
| Force Magnitude [Medium Force: 36.25N Elastic, -20N Viscous] | 0.72 | 2.51 | 0.773 | -3.47 | 2.94 | 0.237 |
| **Random effects** | **Groups** | | **SD** | **Groups** | | **SD** |
| | Trial Number | Intercept | 14.67 | Trial Number | Intercept | 14.06 |
| | Participant | Intercept | 37.32 | Participant | Intercept | 44.23 |
| | Residual | | 174.48 | Residual | | 204.33 |
| | Observations: 19421 | | | Observations: | | |
| **Full Model:** Right-Left Difference in PD or TPD ~ Load Condition + Reach Configuration + Force Magnitude + (1 \| Trial Number) + (1 \| Participant) | | | | | | |

#### Supplementary Table 24. Left/Right Pairwise Comparison, Peak Deceleration (PD)

**T24. Post-Hoc Comparisons of Peak Deceleration (PD) between hands.** Tukey-adjusted pairwise comparisons of left- and right-hand PD within each load condition, across all reach types and force levels.

| Peak Deceleration (PD) | | | | | | | | | | | | | | | | |
|---|---|---|---|---|---|---|---|---|---|---|---|---|---|---|---|---|
| Symmetric Long Reach<br><br>Low Force:<br>30N Elastic, -15N Viscous | | | | | Symmetric Short Reach<br><br>Low Force:<br>30N Elastic, -15N Viscous | | | | Asymmetric Left Long. Right Short Reach<br><br>Low Force:<br>.30N Elastic, -15N Viscous | | | | Asymmetric Left Short, Right Long Reach<br><br>Low Force:<br>30N Elastic, -15N Viscous | | | |
| **Loads Applied: Left/Right Arm** | ***β*** | **SE** | **Z-Ratio** | **P Value** | ***β*** | **SE** | **Z-Ratio** | **P Value** | ***β*** | **SE** | **Z-Ratio** | **P Value** | **Estimate** | **SE** | **Z-Ratio** | **P Value** |
| NL/NL | 48.93 | 11.05 | 4.43 | **<0.001** | 43.58 | 10.98 | 3.97 | **<0.001** | 135.57 | 10.97 | 12.35 | **<0.001** | -56.59 | 10.94 | -5.17 | **<0.001** |
| V/V | 17.86 | 11.24 | 1.59 | 0.112 | 14.03 | 11.08 | 1.27 | 0.205 | 85.82 | 11.38 | 7.54 | **<0.001** | -58.49 | 11.53 | -5.07 | **<0.001** |
| E/E | 47.19 | 11.05 | 4.27 | **<0.001** | 27.61 | 11.03 | 2.50 | **0.012** | 165.56 | 11.00 | 15.05 | **<0.001** | -66.99 | 10.96 | -6.11 | **<0.001** |
| NL/E | -0.25 | 11.12 | -0.02 | 0.982 | -27.74 | 10.98 | -2.53 | **0.012** | 102.13 | 10.91 | 9.36 | **<0.001** | -133.99 | 11.02 | -12.16 | **<0.001** |
| E/NL | 78.00 | 10.99 | 7.10 | **<0.001** | 106.41 | 11.02 | 9.65 | **<0.001** | 183.98 | 10.96 | 16.78 | **<0.001** | -29.56 | 11.06 | -2.67 | **0.008** |
| NL/V | -158.37 | 11.14 | -14.22 | **<0.001** | -184.42 | 11.05 | -16.69 | **<0.001** | -75.39 | 11.38 | -6.63 | **<0.001** | -270.68 | 11.08 | -24.43 | **<0.001** |
| V/NL | 221.88 | 11.24 | 19.74 | **<0.001** | 244.29 | 11.04 | 22.13 | **<0.001** | 307.30 | 11.01 | 27.90 | **<0.001** | 148.42 | 11.31 | 13.12 | **<0.001** |
| V/E | 212.83 | 11.32 | 18.79 | **<0.001** | 164.68 | 11.03 | 14.93 | **<0.001** | 288.97 | 11.02 | 26.22 | **<0.001** | 99.38 | 11.37 | 8.74 | **<0.001** |
| E/V | -149.15 | 11.30 | -13.20 | **<0.001** | -123.68 | 11.13 | -11.11 | **<0.001** | -40.64 | 11.30 | -3.60 | **<0.001** | -247.35 | 11.18 | -22.12 | **<0.001** |
| **Medium Force:<br>36.25N Elastic, -20N Viscous** | | | | | **Medium Force:<br>36.25N Elastic, -20N Viscous** | | | | **Medium Force:<br>36.25N Elastic, -20N Viscous** | | | | **Medium Force:<br>36.25N Elastic, -20N Viscous** | | | |
| NL/NL | 48.93 | 11.05 | 4.43 | **<0.001** | 43.58 | 10.98 | 3.97 | **<0.001** | 135.57 | 10.97 | 12.35 | **<0.001** | -56.59 | 10.94 | -5.17 | **<0.001** |
| V/V | 11.19 | 11.45 | 0.98 | 0.328 | 13.65 | 11.19 | 1.22 | 0.222 | 80.87 | 11.84 | 6.83 | **<0.001** | -56.32 | 11.59 | -4.86 | **<0.001** |
| E/E | 45.50 | 11.02 | 4.13 | **<0.001** | 37.11 | 11.04 | 3.36 | **<0.001** | 116.67 | 11.01 | 10.60 | **<0.001** | -59.06 | 11.00 | -5.37 | **<0.001** |
| NL/E | -2.82 | 11.12 | -0.25 | 0.800 | -57.06 | 11.06 | -5.16 | **<0.001** | 73.96 | 11.00 | 6.72 | **<0.001** | -121.03 | 10.99 | -11.01 | **<0.001** |
| E/NL | 79.67 | 11.02 | 7.23 | **<0.001** | 110.11 | 10.99 | 10.02 | **<0.001** | 209.88 | 11.00 | 19.08 | **<0.001** | -27.94 | 11.05 | -2.53 | **0.011** |
| NL/V | -195.92 | 11.47 | -17.08 | **<0.001** | -199.05 | 11.08 | -17.96 | **<0.001** | -98.28 | 11.64 | -8.44 | **<0.001** | -293.03 | 11.25 | -26.04 | **<0.001** |
| V/NL | 261.66 | 11.34 | 23.06 | **<0.001** | 244.01 | 11.08 | 22.02 | **<0.001** | 342.87 | 11.10 | 30.89 | **<0.001** | 169.02 | 11.51 | 14.69 | **<0.001** |
| V/E | 230.99 | 11.32 | 20.40 | **<0.001** | 179.69 | 11.05 | 16.26 | **<0.001** | 306.07 | 11.08 | 27.63 | **<0.001** | 146.84 | 11.48 | 12.79 | **<0.001** |
| E/V | -151.98 | 11.31 | -13.43 | **<0.001** | -149.98 | 11.11 | -13.50 | **<0.001** | -58.20 | 11.54 | -5.04 | **<0.001** | -258.55 | 11.15 | -23.19 | **<0.001** |

#### Supplementary Table 25. Between-Hand Peak Deceleration (PD) Difference Across Loads (Relative to NL/NL)

**T25. Post-Hoc Comparisons of Peak Deceleration (PD) Differences between Hands by Load Condition and Reach Type.** Pairwise comparisons assess whether hand differences in PD significantly vary across load conditions, using the No Load/No Load condition as the baseline.

| Peak Deceleration (PD) | | | | | | | | | | | | | | | | |
|---|---|---|---|---|---|---|---|---|---|---|---|---|---|---|---|---|
| Symmetric Long Reach<br><br>Low Force:<br>30N Elastic, -15N Viscous | | | | | Symmetric Short Reach<br><br>Low Force:<br>30N Elastic, -15N Viscous | | | | Asymmetric Left Long. Right Short Reach<br><br>Low Force:<br>.30N Elastic, -15N Viscous | | | | Asymmetric Left Short, Right Long Reach<br><br>Low Force:<br>30N Elastic, -15N Viscous | | | |
| **Loads Applied: Left/Right Arm** | ***β*** | **SE** | **Z-Ratio** | **P Value** | ***β*** | **SE** | **Z-Ratio** | **P Value** | ***β*** | **SE** | **Z-Ratio** | **P Value** | **Estimate** | **SE** | **Z-Ratio** | **P Value** |
| V/V | 14.13 | 9.04 | 1.56 | 0.135 | 8.84 | 8.99 | 0.98 | 0.372 | -58.88 | 9.02 | -6.53 | **<0.001** | 46.56 | 9.06 | 5.14 | **<0.001** |
| E/E | -20.57 | 9.19 | -2.24 | **0.040** | 9.18 | 9.16 | 1.00 | 0.372 | -10.20 | 9.20 | -1.11 | 0.306 | 8.23 | 9.27 | 0.89 | 0.375 |
| NL/E | 9.39 | 9.11 | 1.03 | 0.303 | 11.48 | 9.14 | 1.26 | 0.334 | -7.92 | 9.11 | -0.87 | 0.385 | 15.03 | 9.18 | 1.64 | 0.135 |
| E/NL | -19.74 | 9.13 | -2.16 | **0.041** | -6.32 | 9.15 | -0.69 | 0.490 | -13.00 | 9.22 | -1.41 | 0.211 | -14.28 | 9.14 | -1.56 | 0.135 |
| NL/V | -100.03 | 9.01 | -11.10 | **<0.001** | -39.67 | 9.04 | -4.39 | **<0.001** | -81.40 | 9.07 | -8.97 | **<0.001** | -48.82 | 9.05 | -5.40 | **<0.001** |
| V/NL | 114.43 | 9.05 | 12.65 | **<0.001** | 63.18 | 9.04 | 6.99 | **<0.001** | 50.19 | 9.01 | 5.57 | **<0.001** | 106.96 | 9.12 | 11.73 | **<0.001** |
| V/E | 103.05 | 9.23 | 11.16 | **<0.001** | 65.82 | 9.09 | 7.24 | **<0.001** | 42.64 | 9.17 | 4.65 | **<0.001** | 76.76 | 9.16 | 8.38 | **<0.001** |
| E/V | -115.68 | 9.16 | -12.63 | **<0.001** | -48.66 | 9.12 | -5.34 | **<0.001** | -105.89 | 9.17 | -11.54 | **<0.001** | -39.39 | 9.19 | -4.29 | **<0.001** |
| **Medium Force:<br>36.25N Elastic, -20N Viscous** | | | | | **Medium Force:<br>36.25N Elastic, -20N Viscous** | | | | **Medium Force:<br>36.25N Elastic, -20N Viscous** | | | | **Medium Force:<br>36.25N Elastic, -20N Viscous** | | | |
| V/V | -1.00 | 9.10 | -0.11 | 0.916 | 25.42 | 9.00 | 2.83 | **0.008** | -46.68 | 9.08 | -5.14 | **<0.001** | 33.46 | 9.05 | 3.70 | **<0.001** |
| E/E | 0.97 | 9.16 | 0.11 | 0.916 | -0.68 | 9.27 | -0.07 | 0.942 | -16.57 | 9.14 | -1.81 | 0.093 | 13.18 | 9.27 | 1.42 | 0.177 |
| NL/E | -13.46 | 9.09 | -1.48 | 0.202 | 19.74 | 9.10 | 2.17 | **0.040** | -4.08 | 9.15 | -0.45 | 0.656 | 6.02 | 9.18 | 0.66 | 0.512 |
| E/NL | -13.07 | 9.12 | -1.43 | 0.202 | -8.37 | 9.18 | -0.91 | 0.413 | -6.15 | 9.19 | -0.67 | 0.576 | 14.40 | 9.14 | 1.57 | 0.154 |
| NL/V | -111.99 | 9.10 | -12.30 | **<0.001** | -64.20 | 9.03 | -7.11 | **<0.001** | -109.55 | 9.10 | -12.03 | **<0.001** | -63.77 | 9.07 | -7.03 | **<0.001** |
| V/NL | 132.47 | 9.09 | 14.58 | **<0.001** | 82.44 | 9.01 | 9.15 | **<0.001** | 74.52 | 9.08 | 8.21 | **<0.001** | 111.15 | 9.11 | 12.20 | **<0.001** |
| V/E | 95.88 | 9.16 | 10.47 | **<0.001** | 90.79 | 9.10 | 9.97 | **<0.001** | 36.51 | 9.10 | 4.01 | **<0.001** | 84.62 | 9.18 | 9.22 | **<0.001** |
| E/V | -123.70 | 9.16 | -13.51 | **<0.001** | -72.90 | 9.14 | -7.98 | **<0.001** | -106.57 | 9.18 | -11.61 | **<0.001** | -74.99 | 9.14 | -8.21 | **<0.001** |

#### Supplementary Table 26. Left/Right Pairwise Comparison, Time at Peak Deceleration (TPD)

**T26. Post-Hoc Comparisons of Time at Peak Deceleration (PD) between hands.** Tukey-adjusted pairwise comparisons of left- and right-hand TPD within each load condition, across all reach types and force levels.

| Time at Peak Deceleration (TPD) | | | | | | | | | | | | | | | | |
|---|---|---|---|---|---|---|---|---|---|---|---|---|---|---|---|---|
| | Symmetric Long Reach<br><br>Low Force:<br>30N Elastic, -15N Viscous | | | | Symmetric Short Reach<br><br>Low Force:<br>30N Elastic, -15N Viscous | | | | Asymmetric Left Long. Right Short Reach<br><br>Low Force:<br>.30N Elastic, -15N Viscous | | | | Asymmetric Left Short, Right Long Reach<br><br>Low Force:<br>30N Elastic, -15N Viscous | | | |
| **Loads Applied: Left/Right Arm** | ***β*** | **SE** | **Z-Ratio** | **P Value** | ***β*** | **SE** | **Z-Ratio** | **P Value** | ***β*** | **SE** | **Z-Ratio** | **P Value** | **Estimate** | **SE** | **Z-Ratio** | **P Value** |
| NL/NL | 14.24 | 13.36 | 1.07 | 0.286 | -0.60 | 13.32 | -0.04 | 0.964 | 126.27 | 13.33 | 9.48 | **<0.001** | -145.60 | 13.28 | -10.97 | **<0.001** |
| V/V | 14.03 | 13.37 | 1.05 | 0.294 | 2.05 | 13.36 | 0.15 | 0.878 | 183.46 | 13.67 | 13.42 | **<0.001** | -188.05 | 13.86 | -13.57 | **<0.001** |
| E/E | -2.40 | 13.14 | -0.18 | 0.855 | -4.84 | 13.17 | -0.37 | 0.713 | 192.24 | 13.09 | 14.68 | **<0.001** | -189.37 | 13.11 | -14.45 | **<0.001** |
| NL/E | 119.68 | 13.29 | 9.00 | **<0.001** | 34.32 | 13.23 | 2.59 | **0.009** | 266.71 | 13.22 | 20.17 | **<0.001** | -83.88 | 13.29 | -6.31 | **<0.001** |
| E/NL | -138.52 | 13.20 | -10.49 | **<0.001** | -45.40 | 13.22 | -3.43 | **<0.001** | 117.45 | 13.27 | 8.85 | **<0.001** | -309.56 | 13.21 | -23.43 | **<0.001** |
| NL/V | 218.45 | 13.42 | 16.27 | **<0.001** | 173.01 | 13.37 | 12.94 | **<0.001** | 385.50 | 13.67 | 28.21 | **<0.001** | 41.38 | 13.43 | 3.08 | **0.002** |
| V/NL | -223.04 | 13.47 | -16.56 | **<0.001** | -185.29 | 13.35 | -13.88 | **<0.001** | -11.66 | 13.12 | −0.89 | 0.374 | -385.55 | 13.65 | -28.24 | **<0.001** |
| V/E | -82.18 | 13.50 | -6.09 | **<0.001** | -144.45 | 13.38 | -10.79 | **<0.001** | 83.28 | 13.21 | 6.30 | **<0.001** | -311.33 | 13.63 | -22.84 | **<0.001** |
| E/V | 92.07 | 13.38 | 6.88 | **<0.001** | 166.91 | 13.35 | 12.50 | **<0.001** | 333.07 | 13.52 | 24.64 | **<0.001** | -81.04 | 13.38 | -6.06 | **<0.001** |
| **Medium Force:<br>36.25N Elastic, -20N Viscous** | | | | | **Medium Force:<br>36.25N Elastic, -20N Viscous** | | | | **Medium Force:<br>36.25N Elastic, -20N Viscous** | | | | **Medium Force:<br>36.25N Elastic, -20N Viscous** | | | |
| NL/NL | 14.24 | 13.36 | 1.07 | 0.286 | -0.60 | 13.32 | -0.04 | 0.964 | 126.27 | 13.33 | 9.48 | **<0.001** | -145.60 | 13.28 | -10.97 | **<0.001** |
| V/V | 8.50 | 13.65 | 0.62 | 0.534 | 5.73 | 13.53 | 0.42 | 0.672 | 180.92 | 14.18 | 12.76 | **<0.001** | -187.44 | 13.97 | -13.42 | **<0.001** |
| E/E | 2.82 | 13.06 | 0.22 | 0.829 | -20.37 | 13.13 | -1.55 | 0.121 | 183.04 | 13.14 | 13.93 | **<0.001** | -159.45 | 13.03 | -12.24 | **<0.001** |
| NL/E | 116.42 | 13.33 | 8.73 | **<0.001** | 55.21 | 13.33 | 4.14 | **<0.001** | 281.42 | 13.18 | 21.36 | **<0.001** | -61.77 | 13.24 | -4.66 | **<0.001** |
| E/NL | -117.94 | 13.21 | -8.93 | **<0.001** | -55.32 | 13.23 | -4.18 | **<0.001** | 97.76 | 13.35 | 7.32 | **<0.001** | -276.86 | 13.27 | -20.86 | **<0.001** |
| NL/V | 270.05 | 13.78 | 19.59 | **<0.001** | 253.84 | 13.43 | 18.89 | **<0.001** | 459.27 | 14.04 | 32.72 | **<0.001** | 93.99 | 13.55 | 6.93 | **<0.001** |
| V/NL | -253.39 | 13.67 | -18.54 | **<0.001** | -229.43 | 13.43 | -17.08 | **<0.001** | -92.16 | 13.48 | −6.84 | **<0.001** | -458.76 | 13.97 | -32.84 | **<0.001** |
| V/E | -147.00 | 13.52 | -10.87 | **<0.001** | -166.95 | 13.34 | -12.52 | **<0.001** | 8.34 | 13.37 | 0.62 | 0.533 | -389.90 | 13.82 | -28.21 | **<0.001** |
| E/V | 143.92 | 13.49 | 10.67 | **<0.001** | 197.48 | 13.42 | 14.71 | **<0.001** | 353.70 | 13.93 | 25.39 | **<0.001** | 8.94 | 13.36 | 0.67 | 0.503 |

#### Supplementary Table 27. Between-Hand Time at Peak Deceleration (TPD) Difference Across Loads (Relative to NL/NL)

**T27. Post-Hoc Comparisons of Time at Peak Deceleration (TPD) Differences between Hands by Load Condition and Reach Type.** Pairwise comparisons assess whether hand differences in TPD significantly vary across load conditions, using the No Load/No Load condition as the baseline.

| Time at Peak Deceleration (TPD) | | | | | | | | | | | | | | | | |
|---|---|---|---|---|---|---|---|---|---|---|---|---|---|---|---|---|
| **Symmetric Long Reach<br><br>Low Force:<br>30N Elastic, -15N Viscous** | | | | | **Symmetric Short Reach<br><br>Low Force:<br>30N Elastic, -15N Viscous** | | | | **Asymmetric Left Long. Right Short Reach<br><br>Low Force:<br>.30N Elastic, -15N Viscous** | | | | **Asymmetric Left Short, Right Long Reach<br><br>Low Force:<br>30N Elastic, -15N Viscous** | | | |
| **Loads Applied: Left/Right Arm** | **β** | **SE** | **Z-Ratio** | **P Value** | **β** | **SE** | **Z-Ratio** | **P Value** | **β** | **SE** | **Z-Ratio** | **P Value** | **Estimate** | **SE** | **Z-Ratio** | **P Value** |
| V/V | 0.58 | 17.32 | 0.03 | 0.973 | -1.47 | 17.27 | -0.08 | 0.932 | -55.55 | 17.65 | -3.15 | **0.002** | 45.11 | 17.67 | 2.55 | **0.011** |
| E/E | 17.74 | 17.19 | 1.03 | 0.345 | 7.61 | 17.16 | 0.44 | 0.751 | -65.97 | 17.20 | -3.83 | **<0.001** | 49.26 | 17.09 | 2.88 | **0.005** |
| NL/E | -98.34 | 17.40 | -5.65 | **<0.001** | -24.84 | 17.22 | -1.44 | 0.199 | -138.54 | 17.37 | -7.98 | **<0.001** | -48.99 | 17.32 | -2.83 | **0.005** |
| E/NL | 148.33 | 17.31 | 8.57 | **<0.001** | 39.70 | 17.22 | 2.30 | **0.034** | 16.50 | 17.44 | 0.95 | 0.344 | 166.56 | 17.21 | 9.68 | **<0.001** |
| NL/V | -198.91 | 17.50 | -11.37 | **<0.001** | -174.51 | 17.29 | -10.09 | **<0.001** | -257.52 | 17.68 | -14.56 | **<0.001** | -182.03 | 17.37 | -10.48 | **<0.001** |
| V/NL | 233.41 | 17.51 | 13.33 | **<0.001** | 186.03 | 17.27 | 10.77 | **<0.001** | 139.11 | 17.23 | 8.07 | **<0.001** | 246.31 | 17.50 | 14.07 | **<0.001** |
| V/E | 97.93 | 17.47 | 5.61 | **<0.001** | 141.46 | 17.32 | 8.17 | **<0.001** | 36.05 | 17.32 | 2.08 | **0.043** | 168.30 | 17.46 | 9.64 | **<0.001** |
| E/V | -74.34 | 17.32 | -4.29 | **<0.001** | -167.62 | 17.30 | -9.69 | **<0.001** | -208.31 | 17.51 | -11.90 | **<0.001** | -60.00 | 17.34 | -3.46 | **<0.001** |
| **Medium Force:<br>36.25N Elastic, -20N Viscous** | | | | | **Medium Force:<br>36.25N Elastic, -20N Viscous** | | | | **Medium Force:<br>36.25N Elastic, -20N Viscous** | | | | **Medium Force:<br>36.25N Elastic, -20N Viscous** | | | |
| V/V | 8.16 | 17.50 | 0.47 | 0.641 | -6.18 | 17.38 | -0.36 | 0.722 | -56.46 | 17.96 | -3.14 | **0.002** | 46.92 | 17.69 | 2.65 | **0.009** |
| E/E | 12.33 | 17.10 | 0.72 | 0.538 | 16.63 | 17.09 | 0.97 | 0.378 | -58.12 | 17.26 | -3.37 | **0.001** | 17.40 | 16.99 | 1.02 | 0.306 |
| NL/E | -96.63 | 17.45 | -5.54 | **<0.001** | -53.92 | 17.29 | -3.12 | **0.002** | -162.24 | 17.32 | -9.36 | **<0.001** | -75.88 | 17.23 | -4.40 | **<0.001** |
| E/NL | 128.85 | 17.31 | 7.44 | **<0.001** | 54.67 | 17.22 | 3.17 | **0.002** | 23.20 | 17.50 | 1.33 | 0.185 | 128.85 | 17.28 | 7.46 | **<0.001** |
| NL/V | -250.24 | 17.74 | -14.11 | **<0.001** | -254.89 | 17.39 | -14.66 | **<0.001** | -333.87 | 17.92 | -18.63 | **<0.001** | -234.30 | 17.51 | -13.38 | **<0.001** |
| V/NL | 263.02 | 17.65 | 14.90 | **<0.001** | 227.75 | 17.39 | 13.10 | **<0.001** | 208.77 | 17.61 | 11.85 | **<0.001** | 320.09 | 17.82 | 17.96 | **<0.001** |
| V/E | 164.09 | 17.44 | 9.41 | **<0.001** | 162.73 | 17.29 | 9.41 | **<0.001** | 115.59 | 17.47 | 6.62 | **<0.001** | 241.73 | 17.65 | 13.70 | **<0.001** |
| E/V | -128.82 | 17.43 | -7.39 | **<0.001** | -195.03 | 17.42 | -11.19 | **<0.001** | -229.19 | 17.87 | -12.83 | **<0.001** | -143.17 | 17.29 | -8.28 | **<0.001** |
